# *E. coli* phylogeny drives co-amoxiclav resistance through variable expression of *bla*_TEM-1_

**DOI:** 10.1101/2024.08.12.607562

**Authors:** William Matlock, Gillian Rodger, Emma Pritchard, Matthew Colpus, Natalia Kapel, Lucinda Barrett, Marcus Morgan, Sarah Oakley, Katie L. Hopkins, Aysha Roohi, Drosos Karageorgopoulos, Matthew B. Avison, A. Sarah Walker, Samuel Lipworth, Nicole Stoesser

## Abstract

Co-amoxiclav resistance in *E. coli* is a clinically important phenotype associated with increased mortality. The class A beta-lactamase *bla*_TEM-1_ is often carried by co- amoxiclav-resistant pathogens, but exhibits high phenotypic heterogeneity, making genotype-phenotype predictions challenging. We present a curated dataset of *n*=377 *E. coli* isolates representing all 8 known phylogroups, where the only acquired beta- lactamase is *bla*_TEM-1_. For all isolates, we generate hybrid assemblies and co-amoxiclav MICs, and for a subset (*n*=67/377), *bla*_TEM-1_ qPCR expression data. First, we test whether certain *E. coli* lineages are intrinsically better or worse at expressing *bla*_TEM-1_, for example, due to lineage differences in regulatory systems, which are challenging to directly quantify. Using genotypic features of the isolates (*bla*_TEM-1_ promoter variants and copy number), we develop a hierarchical Bayesian model for *bla*_TEM-1_ expression that controls for phylogeny. We establish that *bla*_TEM-1_ expression intrinsically varies across the phylogeny, with some lineages (e.g. phylogroups B1 and C, ST12) better at expression than others (e.g. phylogroups E and F, ST372). Next, we test whether phylogenetic variation in expression influences the resistance of the isolates. With a second model, we use genotypic features (*bla*_TEM-1_ promoter variants, copy number, duplications; *ampC* promoter variants; efflux pump AcrF presence) to predict isolate MIC, again controlling for phylogeny. Lastly, we use a third model to demonstrate that the phylogenetic influence on *bla*_TEM-1_ expression causally drives the variation in co- amoxiclav MIC. This underscores the importance of incorporating phylogeny into genotype-phenotype predictions, and the study of resistance more generally.

## Introduction

The class A beta-lactamase *bla*_TEM-1_ was first identified in 1965 in a clinical *Escherichia coli* isolate^1^. Originally, it was mobilised by two of the earliest named transposons, Tn*2* and Tn*3*, located on plasmids^2^. In the decades since the genetic context of *bla*_TEM-1_ has evolved^3^, and other mobile genetic elements such as IS*26*^4^ and a diverse array of plasmids^5^ contribute to its dissemination. At the time of writing, NCBI contains over 170,000 unique isolates carrying *bla*_TEM-1_ distributed across 28 genera, including the common clinical pathogens *Escherichia coli*, *Klebsiella pneumoniae*, and *Acinetobacter baumannii*^6^. The emergence and dissemination of beta-lactam resistance has been a major healthcare challenge^7^, and *bla*_TEM-1_ represents a key example.

In the UK, beta-lactam and beta-lactamase inhibitor combinations such as co-amoxiclav (amoxicillin and clavulanic acid), are commonly used as a first-line treatment for severe infections^8^. For Enterobacterales, the current EUCAST co-amoxiclav Minimum Inhibitory Concentration (MIC) breakpoint for resistance is 8/2 μg/mL^9^ across all indications, with a recent study concluding that empiric co-amoxiclav treatment of *E. coli* bacteraemia with MICs[ >[32/2 μg/mL was associated with significantly higher mortality in *E. coli* bacteraemia^10^. However, the carriage of *bla*_TEM-1_ is associated with high phenotypic heterogeneity, making genotype-phenotype predictions challenging.

Small-scale, experimental *bla*_TEM-1_ systems have demonstrated that the interplay of location (plasmid or chromosome) and copies in the genome, through varying dosage, contributes to variable resistance^11,12^. In addition, other determinants such as mutations in the promoter of *bla*_TEM-1_^13^ and the chromosomally intrinsic *ampC* gene^14^, and efflux pumps^15^, are associated with *E. coli* beta-lactam resistance. Moreover, different regulatory systems^16,17^, epistasis (interaction between genes)^18,19^, and epigenetics (heritable phenotypic changes without alterations to the underlying DNA sequence)^20^, might also influence co-amoxiclav resistance. For example, five different *E. coli* strains carrying the same pLL35 plasmid (which carries *bla*_CTX-M-15_ and *bla*_TEM-112_) varied in cefotaxime resistance^21^. Likewise, the introduction of a pOXA-48 plasmid to six different *E. coli* strains resulted in variable co-amoxiclav resistance^22^. This indicates that strain background plays a role in resistance.

Successful genotype-to-phenotype prediction requires a comprehensive understanding of not only individual resistant determinants but also their combined effects. Moreover, this understanding must be translated to clinically relevant pathogens. Yet, to accurately model resistance in these systems, a large sample of linked genomic and phenotypic data is required, which until recently has been limited by sequencing technology and costs.

In this study, we curated and completely reconstructed the genomes of nearly 400 clinical *E. coli* bacteraemia isolates to reflect a “real-world” but relatively simple genetic scenario where the only known acquired beta-lactamase gene identified was *bla*_TEM-1_, all identical at the amino acid level. We quantified the co-amoxiclav MICs for these isolates and generated *bla*_TEM-1_ qPCR expression data for a subset. We then modelled *bla*_TEM-1_ expression and co-amoxiclav MIC whilst controlling confounding genetic mechanisms and chromosomal phylogeny.

## Materials and methods

### Isolate selection

We considered *n*=548 candidate *E. coli* bacteraemia isolates cultured from patients presenting to Oxford University Hospitals NHS Foundation Trust between 2013–2018, and selected from a larger study of systematically sequenced isolates based on screening their short-read only assemblies with NCBIAMRFinder (v. 3.11.2) for *bla*_TEM-1_, and the absence of other beta-lactamases^23^.

### DNA extraction and sequencing

Sub-cultures of isolate stocks, stored at −80°C in 10[% glycerol nutrient broth, were grown on Columbia blood agar (CBA) overnight at 37°C. DNA was extracted using the EasyMag system (bioMerieux) and quantified using the Broad Range DNA Qubit kit (Thermo Fisher Scientific, UK). DNA extracts were multiplexed as 24 samples per sequencing run using the Oxford Nanopore Technologies (ONT) Rapid Barcoding kit (SQK-RBK110.96) according to the manufacturer’s protocol. Sequencing was performed on a GridION using version FLO- MIN106 R9.4.1 flow cells with MinKNOW software (v. 21.11.7) and basecalled using Guppy (v. 3.84). Short-read sequencing was performed on the Illumina HiSeq 4000 pooling 192 isolates per lane, generating 150bp paired end-reads^24^.

### Dataset curation and genome assembly

Full details are given in Supplementary File 1. Briefly, short- and long-read quality control used fastp (v. 0.23.4) and filtlong (v. 0.2.1), respectively^25,26^. We also used Rasusa (v. 0.7.1) on *n*=3/548 long-read sets due to memory constraints^27^. Genome assembly used Flye (v. 2.9.2-b1786) with bwa (v. 0.7.17-r1188) and Polypolish (v. 0.5.0), and Unicycler (v. 0.5.0) which uses SPAdes (v. 3.15.5), miniasm (v. 0.3-r179), and Racon (v. 1.5.0)^28–34^. Plasmid contig validation used Mash screen (v. 2.3) with PLSDB (v. 2023_06_23_v2)^35,36^. All assemblies were annotated with NCBIAMRFinder (v. 3.11.26 and database v. 2023-11- 15.1)^23^. Alongside, we validated the presence of *bla*_TEM-1_ using tblastn (v. 2.15.0+) with the NCBI Reference Gene Catalog TEM-1 RefSeq protein WP_000027057.1 and 100% amino acid identity^37^. Following genome assembly, we removed *n*=171/548 isolates, either because (i) the chromosome did not circularise (116/171), (ii) it carried a non-*bla*_TEM-1_ *bla*_TEM_ variant and/or an additional acquired beta-lactamase (54/171), or (iii) the chromosome was too short consistent with misassembly (∼3.5Mbp; 1/171). This left a final dataset of *n*=377 isolates.

### Antibiotic susceptibility testing

Antibiotic susceptibility testing was performed using the BD Phoenix™ system in accordance with the manufacturers’ instructions, generating MICs for co-amoxiclav.

### Generation of cDNA template

RNA extraction and DNase treatment were performed on replicates of each isolate (*n*=3 biological/*n*=3 technical) as described previously^38^. RNA was quantified post DNase treatment using Broad Range RNA Qubit kit (Thermo Fisher Scientific, UK), normalised to 1 μg and reverse transcribed to cDNA using SuperScript IV VILO (Thermo Fisher Scientific, UK) under the following conditions: 25°C for 10 minutes, 42°C for 60 minutes and 85°C for 5 minutes.

### qPCR quantification of *bla*_TEM-1_ expression

*bla*_TEM-1_ expression was quantified in a selection of isolates: initially *n*=35 isolates in triplicate, referred to as batch 1; then a further *n*=48 isolates in duplicate referred to as batch 2. Batch 1 were randomly selected stratified by MICs (*n*=2 MIC <=2, *n*=5 MIC 4/2, *n*=9 MIC 8/2, *n*=10 MIC 16/2, *n*=4 MIC 32/2, *n*=5 MIC >32/2). Batch 2 was enhanced for specific *bla*_TEM-1_ promoter mutations, selecting all isolates with a single *bla*_TEM-1_ gene with C32T (with or without a G146A mutation) that had not already been tested, and then randomly selecting from other wildtype and G146A, single *bla*_TEM-1_ gene isolates. For all qPCR reactions, *E. coli* cDNA was normalised to 1ng and amplified in a duplex qPCR reaction targeting *bla*_TEM-1_ and 16S. qPCR standard curves were prepared for both *bla*_TEM-1_ (Genbank Accession: DQ221255.1) and 16S (Genbank Accession: LC747145.1) sequences cloned into pMX vectors (Thermo Fisher Scientific, UK). Tenfold dilutions of linearised plasmids (1- 1x10^7^ copies/reaction) were used as a standard curve for each experiment. Both curves were linear in the range tested (16S: R^2^>0.991; TEM-1: R^2^>0.91,). The slopes of the standard curves for 16S and *bla*_TEM-1_ were -3.607 and -3.522 respectively. qPCR was performed using a custom 20 μl TaqMan gene expression assay consisting of TaqMan™ Multiplex Master Mix, TaqMan unlabelled primers and a TaqMan probe with dye label (FAM for TEM-1 and VIC for 16S) carried out on the QuantStudio5^TM^ real-time PCR system (Thermo Fisher Scientific, UK). Cycling conditions were 95°C for 20 seconds, followed by 40 cycles of 95°C for 3[seconds and 60°C for 30 seconds, with Mustang purple as the passive reference. For batch 1, triplicate samples were analysed and standardized against 16S rRNA gene expression. Triplicate reactions for each isolate demonstrated good reproducibility for batch 1 (Figure S1). Of note, for isolate OXEC-75, TEM-1 expression was very low-level, and 5 reactions (1 technical replicate for biological replicate 1, 1 technical replicate for biological replicate 2, and all 3 technical replicates for biological replicate 3) failed to amplify any product. Due to resource constraints, we reduced replicates for batch 2 (*n*=1 biological/*n*=2 technical; Figure S2). To reduce model complexity, we omitted some batch 1 isolates (*n*=16/35) which carried more than one copy of *bla*_TEM-1_ in the genome, leaving a total of *n*=67 isolates. ΔCt values were calculated by subtracting mean 16S Ct from mean TEM-1 Ct.

### Assembly annotations

We annotated the chromosomes using Prokka (v . 1.14.6) with default parameters except `-- centre X --compliant` (see `annotate.sh`)^39^. Abricate (v. 1.0.1) was used with default parameters and the PlasmidFinder database (v. 2023-Nov-4) to annotate for plasmid replicons^40,41^. Plasmid mobilities were predicted using MOB-suite’s MOB-typer (v. 3.1.4) with default parameters^41^. Briefly, a plasmid was labelled as putatively conjugative if it had both a relaxase and mating pair formation (MPF) complex, mobilisable if it had either a relaxase or an origin of transfer (oriT) but no MPF, and non-mobilisable if it had no relaxase and oriT. Lastly, we assigned sequence types (STs) and phylogroups to our *E. coli* chromosomes using mlst (v. 2.23.0) with default parameters and EzClermont (v. 0.7.0) default parameters, respectively^42,43^. We used blastn (v. 2.15.0+) with a custom database of known *bla*_TEM-1_ promoters^13,44,45^. Due to the high similarity between the P3, Pa/b, P4, and P5 reference sequences, we chose the top hit in each position.

### SNV analysis

We first determined the sets of sequences we wanted to align: (i) *bla*_TEM-1_ (*n*=451; some genomes carried multiple copies), (ii) *bla*_TEM-1_ promoters (*n*=409), (iii) *ampC* (*n*=377), and (iv) *ampC* promoters (*n*=377). For *bla*_TEM-1_ and *ampC*, we extracted the relevant sequences using the coordinate and strand information from the NCBIAMRFinder output (see `extractGene.py`). For the *bla*_TEM-1_ promoters, we used coordinate and strand information from the earlier blastn results. For the *ampC* promoters, we took the sequence 200bp upstream of the *ampC* gene then manually excised the −42 to +37 region in AliView^46^. Sets of sequences were aligned using MAFFT (v. 7.520) with default parameters except `--auto`^47^. Variable sites were examined using snp-sites (v. 2.5.1) with default parameters and in `-v` mode^48^.

### Contig copy number

We used BWA (v. 0.7.17-r1188) to map the quality-controlled short-reads to each contig, then SAMtools (v. 1.18) for subsequent processing (see `copyNumber.sh`)^29,49^. For each contig, we calculated the mean depth over its length, then within each assembly, normalised by the mean depth of the chromosome.

### Chromosomal core gene phylogeny

Building the chromosomal phylogeny involved four main steps: annotating the chromosomes, identifying the core genes, aligning them, and building a phylogeny. Initially, all the chromosomes carried a copy of *ampC*, meaning it was a core gene and would be included in the phylogeny. Since we wanted to manually verify EzClermont phylogroup classifications with the phylogeny and then compare phylogroups to the distribution of *ampC* gene variants, we excised the *ampC* sequence from all the chromosomes beforehand to avoid confounding our analysis (see `removeGene.py`). To identify the core genes (those with ≥98% frequency in the sample), we used Panaroo (v. 1.4.2) with default parameters except `--clean-mode sensitive --aligner mafft -a core --core_threshold 0.98`^50^. Panaroo also aligned our core genes using MAFFT (v. 7.520; see `runPanaroo.sh`)^47^. Lastly, we built the core gene maximum-likelihood phylogeny using IQ-Tree (v. 2.3.0) with default parameters except -`m GTR + F + I + R4 -keep-ident -B 1000 -mem 10G` using `-s core_gene_alignment_filtered.aln` from Panaroo (see `runIQTREE.sh`)^51^. The substitution model used was general time reversible (GTR) using empirical base frequencies from the alignment (F), allowing for invariant sites (I) and variable rates of substitution (R4).

### Statistical analysis and visualisation

All statistical analysis was performed in R (v. 4.4.0) using RStudio (v. 2024.04.2+764)^52,53^.We implemented MCMC generalised linear mixed models using the MCMCglmm library in R^54^. Model specifications, convergence diagnostics, parameter estimations, and outputs are reported in Supplementary File 2; see `modelExpression.R`, `modelMIC.R`, and `modelCombined.R`, to reproduce the *bla*_TEM-1_ expression, co-amoxiclav MIC, and causal models, respectively). Homogeneity and completeness are defined in Rosenberg, A. and Hirschberg, J (2007) and were also implemented in R^55^. A 95% highest posterior density (HPD) credible interval finds the closest points (*a* and *b*) for which *F(b) - F(a)* = 0.95, where *F* is the empirical density of the posterior. Figures were plotted with the ggplot2 library^56^.

### Computational reproducibility

See `buildResults.R` to reproduce all statistics and figures in the manuscript. All scripts referenced in the Materials and methods can be found in https://github.com/wtmatlock/tem.

### Data availability

Metadata for all *n*=377 genomes included in the final analysis is given in Supplementary Table 1. Metadata for all *n*=451 *bla*_TEM-1_ annotations identified in these genomes is given in Supplementary Table 2. qPCR expression data for all replicates is given in Supplementary Table 3. NCBI accessions for short- and long-read sets and assemblies are given in Supplementary Table 4.

## Results

### A curated dataset of *E. coli* isolates with hybrid assemblies and co-amoxiclav MICs

We began with *n*=548 candidate *E. coli* isolates, which following hybrid assembly, were curated into a final dataset of *n*=377/548 (see Materials and methods and Supplementary File 1). In total, 77% (291/377) of assemblies were complete (all contigs were circularised), with the remaining 27% (86/377) having at least a circularised chromosome to confidently distinguish between chromosomal and plasmid-associated *bla*_TEM-1_. Assemblies contained median=3 (IQR=2-5) plasmid contigs.

We identified *n*=451 *bla*_TEM-1_ genes on 431 contigs (13% [58/431] chromosomal versus 87% [373/431] plasmid). Isolates carried a median=1 copy of *bla*_TEM-1_ (range=1-6). Carrying more than one copy of *bla*_TEM-1_ on a single contig was rare: of all *bla*_TEM-1_-positive contigs, 97% (400/412) versus 3% (12/412) had no duplications versus at least one. The *bla*_TEM-1_ genes had synonymous single nucleotide polymorphisms (SNPs) in positions 18, 138, 228, 396, 474, 705, and 717, totalling *n*=7 single nucleotide variant (SNV) profiles across the replicons, yet diversity was dominated by *bla*_TEM-1b_ at 73% (329/451; SNV profile TATTTCG; see Materials and methods)^57^. Where *bla*_TEM-1_-positive contigs had at least two copies of *bla*_TEM-1_, they were almost always the same SNV duplicated (11/12).

By examining the “genomic arrangement” of *bla*_TEM-1_ (namely the replicons it was found on as well as any copies), we found most isolates carried a single non-chromosomal copy (73.5% [277/377]; Figure 1a-b). More generally, whilst the plasmid contigs totalled only 3.6% of the total sequence length (bp) across the assemblies (71,126,646bp/1,969,804,202bp), they carried 85.8% of the *bla*_TEM-1_ genes (387/451). Such *bla*_TEM-1_-carrying plasmids were represented across the *E. coli* phylogeny (Figure 1c).

**Figure 1.**
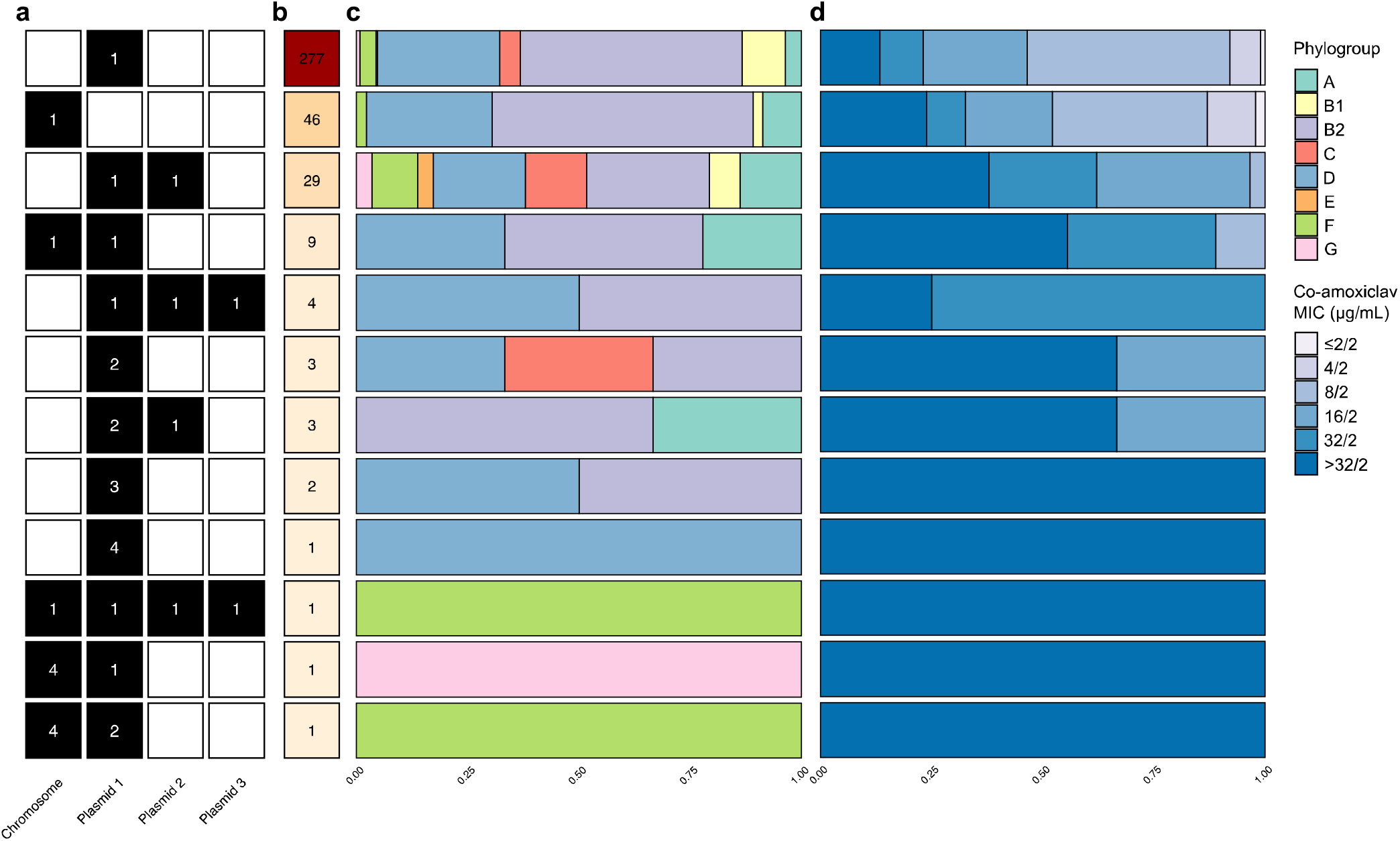
A genotypically and phenotypically heterogenous population of *bla*_TEM-1_-carrying *E. coli*. **(a)** nomic arrangement of *bla*_TEM-1_ in the genomes ordered in descending **(b**) frequency. **(c)** Phylogroup co-amoxiclav MIC distribution for each genomic arrangement.

Overall, the dataset comprised 5.6% (21/377) phylogroup A, 8.0% (30/377) B1, 48.5% (183/377) B2, 4.8% (18/377) C, 27.3% (103/377) D, 0.5% (2/377) E, 4.2% (16/377) F, and 1.1% (4/377) G. In total, we manually corrected *n*=5 EzClermont phylogroup classifications using a core-gene phylogeny (see Materials and methods): OXEC-108 (G to D), OXEC-317 (B2 to D), OXEC-333 (U to B1), OXEC-344 (U to B1), and OXEC-406 (U to B1). The EzClermont publication presented a 98.4% (123/125) true-positive rate on their validation set, which is in line with our ∼98.7% (372/377)^43^.

Five known upstream promoters modulate the expression of *bla*_TEM-1_: P3, Pa/Pb, P4, P5, and Pc/d ^13,44^. We linked 91% (409/451) *bla*_TEM-1_ genes to a promoter immediately upstream, of which a majority, 64% (262/409), were identical to the P3 reference. More generally, excluding *n*=2 different Pc/Pd-like promoters which have large deletions, we identified SNPs in positions 32, 43, 65, 141, 162, and 175, totalling *n*=8 SNV profiles (by Sutcliffe numbering^58^). Notably, 15% (39/262) of promoters had the Pa/Pb-associated C32T mutation which produces two overlapping promoter sequences.

Isolates were associated with a diverse range of co-amoxiclav MICs for the 377 isolates (μg/mL; ≤2/2 [4 (1.1%)], 4/2 [24 (6.4%)], 8/2 [144 (38.2%)], 16/2 [86 (22.8%)], 32/2 [44 (11.7%)], and >32/2 [75 (19.9%)]; Figure 1d; see Materials and methods).

For a genome carrying *bla*_TEM-1_ on the chromosome, the gene’s copy number and total number of genes in the genome are equivalent. For a genome carrying *bla*_TEM-1_ on a plasmid, this might not be the case. This is because plasmids can exist as multiple copies. The calculated copy number of all plasmidic contigs (*n*=1,512) was median=3.13 (range=0.04- 57.00). Of these, *n*=19/1,512 contigs (with 6/19 circularised) had calculated copy numbers less than one (see Materials and methods). This was potentially due to uneven short-read coverage. However, none carried *bla*_TEM-1_ and so were not used in the later modelling. Taking the circularised plasmids with copy number at least one (1,030/1,512), longer plasmids (>10kbp; 668/1,030) were generally low copy number (median=2.36), whilst shorter plasmids (≤10kbp; 362/1,030) were generally high copy number (median=11.01, see Figure S3), consistent with previous studies^59^.

### *E. coli* phylogeny shapes *bla*_TEM-1_ expression

Within different *E. coli* lineages, *bla*_TEM-1_ and its promoter are potentially subject to different regulatory systems and epigenetic interactions, which may in turn affect *bla*_TEM-1_ expression. To test this, we first selected a random subsample of *n*=67/377 isolates with a single copy of *bla*_TEM-1_ in the genome, either on a chromosome (15/67) or a plasmid (52/67). Moreover, we only selected isolates with zero, one, or two mutations in the *bla*_TEM-1_ promoter sequence: C32T, which produces two overlapping promoters and is known to increase expression^13^, and G175A (according to Sutcliffe numbering based on the PBR322 plasmid^58^; see Table 1).

**Table 1.** Replicon distribution of *n*=67 *bla*_TEM-1_ promoter variants. Single nucleotide variants (SNVs) are for the 32^nd^ and 175^th^ positions by Sutcliffe numbering^58^.

| Promoter SNV / Replicon | Chromosome | Plasmid | Total |
| --- | --- | --- | --- |
| CG (wildtype) | 1 | 22 | 23 |
| G175A | 2 | 21 | 23 |
| C32T | 3 | 5 | 7 |
| C32T, G175A | 9 | 4 | 13 |
| Total | 15 | 52 | 67 |

Isolates were distributed across the entire *E. coli* phylogeny (phylogroup A [6/67], B1 [4/67], B2 [40/67], C [4/67], D [8/67], E [1/67], and F ([4/67]). We then performed qPCR to evaluate for *bla*_TEM-1_ expression (see Materials and methods). Every isolate had at least two replicates (2 [48/67], 4 [1/67], or 9 [18/67), giving a total of *n*=262 *bla*_TEM-1_ ΔCt observations (TEM-1 Ct – 16S Ct; see Materials and methods) for modelling.

To test for the effects of *E. coli* lineage, we built a maximum likelihood core gene phylogeny for all *n*=377 chromosomes (see Materials and methods). In total, we identified 17,836 gene clusters, of which 18.7% (3,342/17,836) were core genes (those found in ≥98% of chromosomes). The phylogeny (midpoint rooted and restricted to the *n*=67/377 isolates in the expression analysis) is given in Figure 2a. Using *b*=1000 ultrafast bootstraps, all phylogroup node supports were 100%, and more generally, 76.7% (287/374) of internal node supports were 100%, and 87.4% (327/374) were at least 95% (see Materials and methods).

**Figure 2.**
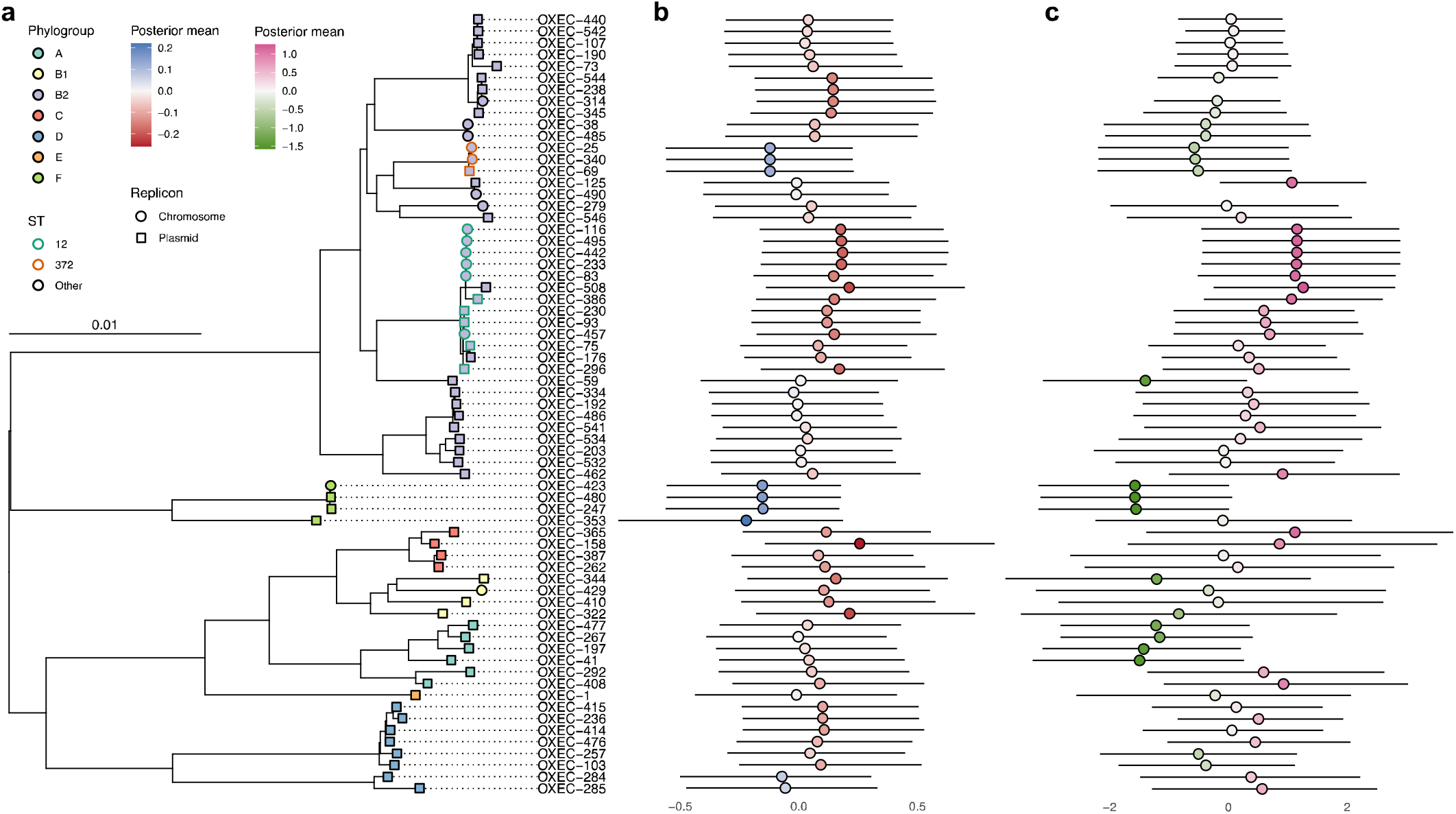
Intrinsic expression of *bla*_TEM-1_ shapes co-amoxiclav MIC across the *E. coli* phylogeny. **(a)** A midpoint-rooted core gene phylogeny of *E. coli* chromosomes from the 67 isolates with expression quantified. Tips are coloured by phylogroup. ST12 (higher than average effect) and ST372 (lower than average effect) are highlighted by green and orange outlines, respectively. Tip shape distinguishes location of *bla*_TEM-1_ on chromosomes (circles) and plasmids (squares). **(b)** Posterior means (coloured circles) and 95% HPD intervals (horizontal lines) for phylogenetic effect on negative *bla*_TEM-1_ ΔCt for each tip (multiplied by −1 for ease of comparison). Red indicates above average expression and blue indicates below average expression. **(c)** Posterior means (coloured circles) and 95% HPD intervals (horizontal lines) for phylogenetic effect on co- amoxiclav MIC for each tip. Pink indicates above average MIC and green indicates below average MIC.

Moreover, the Robinson-Foulds distance between the ML tree and consensus tree was 4, indicating nearly identical topology.

Briefly, the expression linear mixed model employed Markov Chain Monte Carlo (MCMC) to estimate parameters. The response variable *bla*_TEM-1_ ΔCt (normalised and 95^th^ percentile truncated) was related to the fixed effects (i) *bla*_TEM-1_ cell copy number (normalised), (ii) presence of the C32T promoter mutation, (iii) presence of the G175A mutation, and (iv) their interaction. Random effects were incorporated to account for qPCR replicates and phylogenetic relationships between isolates. See Supplementary File 2 for model specification, outputs, and diagnostics.

In decreasing order of effect size, C32T, G175A, and a one unit increase in contig copy number all increased expression (decreased ΔCt; Table 2). There was no additional effect of G175A if C32T was also present (-1.69 < −1.71). The posterior for contribution of variance from phylogeny demonstrated a long right tail (mean=0.07; 95% highest posterior density, HPD=[0.00, 0.21]; see Materials and methods), suggestive of high heterogeneity. For qPCR replicates, the contribution of variance exhibited minimal skew (mean=0.15; 95% HPD=[0.08, 0.23]). To investigate this further, we computed the posterior mean and 95% HPD credible interval for each tip in the phylogeny (Figure 2b). Compared to the average across the *E. coli* phylogeny, some phylogroups (B1, C) and STs (12) were associated with increased *bla*_TEM-1_ expression, whilst some phylogroups (E, F) and STs (372) were associated with decreased *bla*_TEM-1_ expression.

**Table 2.** Parameter estimates for *bla*_TEM-1_ ΔCt genotype-phenotype model. All values are taken from chain 1. For Replicates and Phylogeny, estimates represent the average contribution to variance across the sample. Effect of both C32T* and G175A* is −1.71- 0.31+0.33=-1.69.

| Variable | Beta coefficient<br>posterior mean | 95% HPD (l, u) | *p*<sub>MCMC</sub> |
| --- | --- | --- | --- |
| Intercept | 0.28 | 0.03, 0.59 | 0.0245 |
| C32T* | -1.71 | -2.08, -1.34 | <1e-05 |
| G175A* | -0.31 | -0.57, -0.04 | 0.0202 |
| C32T*:G175A* interaction | 0.33 | -0.14, 0.81 | 0.1779 |
| *bla*<sub>TEM-1</sub> cell copy number | -0.12 | -0.24, 0.01 | 0.0679 |

\* Numbering by Sutcliffe<sup>58</sup>

### *ampC* gene variation is highly concordant with *E. coli* lineage

In *E. coli*, the chromosomally intrinsic *ampC* encodes a class C beta-lactamase with expression typically induced from an external stimulus, namely the presence of a penicillin such as amoxicillin, or a beta-lactamase inhibitor such as clavulanic acid. At the time of writing, the beta-lactamase database (BLDB) contains *n*=2,281 non-synonymous variants of the gene.

To quantify how well *ampC* variants agree with phylogroup and ST, we calculated the homogeneity (*h*) and completeness (*c*; both range from 0 to 1; see Materials and methods). Briefly, *h* = 1 means that a phylogroup or ST contains a single *ampC* variant. Conversely, *c* = 1 means that all instances of an *ampC* variant fall within the same phylogroup or ST. For phylogroups, we found *h* = 0.489 and *c* = 0.964, and for STs (excluding 38/377 which were unassigned), *h* = 0.938 and *c* = 0.877. Overall, this suggests that phylogroups tend to contain distinct *ampC* variants, which are generally ST-specific, and overall, that *E. coli* phylogeny is a suitable proxy for *ampC* variation.

Whilst many *E. coli ampC* variants present a narrow spectrum of hydrolytic activity, some can potentially hydrolyse third-generation cephalosporins following mutations in the promoter sequence. To explore promoter variation, we aligned all *n*=377 *ampC* promoter sequences. Mutations outside positions -42 to +37 (according to Jaurin numbering^14^) were disregarded based on existing characterisations^60,61^. In total, *n*=12 *ampC* promoter SNVs were identified, with variation dominated by the *E. coli* K12 wildtype at 47% (177/377). Table 3 documents all *n*=11 mutations identified. A given *ampC* variant associated almost uniquely with an *ampC* promoter variant, yet *ampC* promoter variants were associated with multiple *ampC* variants (*h* = 0.483 and *c* = 0.941).

**Table 3.** Variation in *n*=377 *ampC* promoters. Positions are according to Jaurin numbering^14^.

| Region | Position | *E. coli* K12 (n) | Mutation (n) |
| --- | --- | --- | --- |
| Spacer | -28 | G (243) | A (134) |
|  | -18 | G (329) | A (48) |
| Between -10 box and attenuator | -1 | C (329) | T (48) |
|  | +11 | T (371) | - (6) |
| Attenuator | +17 | C (316) | T (61) |
|  | +22 | C (367) | T (10) |
|  | +26 | T (367) | G (10) |
|  | +27 | A (367) | T (10) |
|  | +30 | G (376) | A (1) |
|  | +32 | G (364) | A (13) |
|  | +37 | G (376) | T (1) |

### Efflux pump AcrF is not encoded by all *E. coli*

Efflux pumps can confer resistance to antibiotics by transporting molecules outside the cell^15^. In particular, members of the resistance-nodulation-cell division superfamily (RND), including AcrF (encoded by *acrF*), can export penams. We annotated our *n*=377 genomes for efflux pump genes, totalling *n*=1,067 hits (at least 100% coverage and at least 95% identity; see Materials and methods), invariably on the chromosome. Chromosomes had median=3 annotations (range=1-4). AcrF was encoded by 65% (244/377) of isolates, always as a single copy. Moreover, it was widely distributed across the phylogeny: 8.2% (20/244) phylogroup A, 11.1% (27/244) B1, 29.9% (73/244) B2, 7.0% (17/244) C, 38.9% (95/244) D, 0.8% (2/244) E, 2.5% (6/244) F, and 1.6% (4/244) G. All other efflux pump annotations are given in Supplementary Table 1.

### *E. coli* phylogeny drives co-amoxiclav resistance through expression

We next investigated whether the *E. coli* lineages with intrinsically higher *bla*_TEM-1_ expression also had intrinsically higher co-amoxiclav MICs. This would be consistent with lineage differences in regulatory regions and epigenetic interactions driving increased resistance.

We employed an MCMC to estimate parameters in an ordinal mixed model. The response variable isolate co-amoxiclav MIC (μg/mL; levels ≤2/2, 4/22, 8/22, 16/2, 32/2, >32/2) was predicted by the fixed effects (i) *bla*_TEM-1_ cell copy number (normalised and 95^th^ percentile truncated), (ii) *bla*_TEM-1_ genome copy number (>1 vs. 1), (iii) non-wildtype *bla*_TEM-1_ promoter SNVs, (iv) non-wildtype *ampC* promoter SNVs, and (v) presence of AcrF. For the model, we only used isolates for which every *bla*_TEM-1_ gene was linked to a promoter, and all the promoters were the same variant. We then filtered out isolates with *bla*_TEM-1_ promoter and *ampC* promoter variants that appeared less than 10 times. This left *n*=292/377 isolates. Full model specification, convergence diagnostics, and outputs are given in Supplementary File 2.

In decreasing order of effect size, the presence of C32T and G175A in the *bla*_TEM-1_ promoter, the presence of just C32T, *bla*_TEM-1_ cell copy number, and *bla*_TEM-1_ genome copy number all increased co-amoxiclav MIC (Table 4); the remaining effects were compatible with chance. As with the expression model, the posterior distribution for contribution of variance from phylogeny demonstrated a long right tail (mean=2.85; 95% HPD=[0.75, 5.25]). The phylogeny for the *n*=292 isolates with co-amoxiclav MIC tip effects is given in Figure S4.

**Table 4.** Parameter estimates for co-amoxiclav MIC genotype-phenotype model. All values are taken from chain 1.

| Variable | | Beta coefficient<br>posterior mean | 95% HPD (l, u) | $p_{\text{MCMC}}$ |
| --- | --- | --- | --- | --- |
| Intercept |  | 3.89 | 2.89, 4.89 | <1e-05 |
| *bla*<sub>TEM-1</sub> cell copy number |  | 2.01 | 1.34, 2.68 | <1e-05 |
| *bla*<sub>TEM-1</sub> genome copy number (>1 vs. 1) |  | 0.99 | 0.04, 1.93 | 0.0399 |
| *bla*<sub>TEM-1</sub> promoter<br>SNV vs. wildtype | G175A* | 0.16 | -0.37, 0.67 | 0.5435 |
|  | C32T* | 6.06 | 4.14, 8.06 | <1e-05 |

|  |  |  |  |  |
| --- | --- | --- | --- | --- |
|  | C32T* and G175A* | 5.88 | 3.91, 7.94 | <1e-05 |
| *ampC* promoter<br>SNV vs. wildtype | G-28A<sup>†</sup> and C17T<sup>†</sup> | 0.44 | -0.62, 1.51 | 0.4096 |
|  | G-28A<sup>†</sup> | 0.87 | -0.93, 2.60 | 0.3152 |
|  | G-18A<sup>†</sup> and C-1T<sup>†</sup> | 0.83 | -1.45, 3.07 | 0.4444 |
| AcrF |  | 0.02 | -0.51, 0.54 | 0.9475 |

\* Numbering by Sutcliffe<sup>58</sup>
† Numbering by Jaurin<sup>14</sup>

Lastly, we developed a combined model to test whether the phylogenetic influence on *bla*_TEM-1_ expression casually varies co-amoxiclav MIC (see Supplementary File 2). Here, we only used the predictors identified as significant from previous expression and MIC models (*bla*_TEM-1_ cell and genome copy numbers, and *bla*_TEM-1_ promoter SNV). Briefly, the model estimates a parameter that scales the phylogenetic and non-phylogenetic random effects from expression to MIC. Under causality, the scaling parameter should be constant across all random effect terms. We found the scaling parameter had a posterior mean=-1.13 (95% HPD=[-0.72, −1.49]; *p*_MCMC_=0.002; values taken from chain 1). This supports a direct and substantial influence of expression on MIC, mediated by phylogenetic relationships.

## Discussion

In our dataset of clinical *E. coli*, *bla*_TEM-1_ was overwhelmingly carried by conjugative plasmids. These means it can spread between bacterial hosts and different genetic backgrounds. We demonstrated that different bacterial hosts intrinsically vary in their ability to express *bla*_TEM-1_ when accounting for variation in promoters (C32T and G175A mutations) and contig copy number. Moreover, our findings suggest that some clinically successful lineages (e.g. ST12) are better at expressing *bla*_TEM-1_ than less clinically successful lineages (e.g. phylogroup F). With a second model, we also found that different *E. coli* lineages vary intrinsically in co-amoxiclav MIC (accounting for *bla*_TEM-1_ genome and cell copies, *bla*_TEM-1_ and *ampC* promoter variants, and AcrF presence). Again, we observed that some clinically successful lineages (e.g. ST12) had higher resistance than less clinically successful lineages (e.g. phylogroup F). A third model demonstrated that these two traits were casually linked: *E. coli* phylogeny drives co-amoxiclav resistance through variable expression of *bla*_TEM-1_. We believe this finding should generalise to other AMR genes, and underscores the necessity of fully resolving bacterial genomes to incorporate accurate genetic, genomic, and phylogenetic information in resistance prediction models. Future work could include evaluations of single amino acid substitutions in TEM-1 (that hydrolyse third-generation cephalosporins and carbapenemases^62^) which are typically carried in more complex genetic backgrounds.

This study has limitations. Firstly, it is possible that, due to fragmented plasmid assemblies, some isolates identified as having multiple copies of *bla*_TEM-1_ on multiple plasmids instead had multiple copies on the same plasmid. Nonetheless, our expression analysis only considered isolates with a single copy of *bla*_TEM-1_ in the genome, mitigating this concern.

Secondly, we only examined expression for a subsample of our isolates due to resource limitations. Thirdly, whilst there is not an agreed upon standard reference gene for quantifying beta-lactamase expression^63–65^, previous work has shown 16S to be stable^64^.

Crucially, our delta Ct values were consistent within isolates. Fourthly, whilst we found no signal for the AcrF pump influencing co-amoxiclav MIC, recent work on ST11 suggests that not all *E. coli* possess a functional copy of the gene^66^. However, it was out of scope of this work to characterise all AcrF gene variants. We also only observed two potentially relevant porin mutations (a premature stop codon in OmpC on OXEC-40’s chromosome and in OmpF on OXEC-423’s chromosome; see Supplementary Table 1), limiting our ability to investigate their effects on phenotype. Penultimately, automated susceptibility testing methods, like the BD Phoenix™ used here, may not agree completely with reference methods; yet previous work has shown strong agreement with the EUCAST agar dilution method^67^. Lastly, plasmid copy number is not static. Moreover, in the presence of antibiotics, it has been demonstrated that resistance gene-carrying plasmids can increase their copy number to increase the chance of survival^68^. Our point estimates of plasmid copy number were derived from genome assemblies sequenced in the absence of antibiotics, which likely represent a lower bound.

Nonetheless, we found strong signal to suggest the import of plasmid copy number on resistance, even if under our sensitivity testing, plasmid copy number potentially increased within isolates.

Future studies should examine specific regulatory pathways, epistatic interactions, and epigenetic mechanisms such as DNA methylation, that link phylogenetic background with *bla*_TEM-1_ expression and resistance. Additionally, examining other resistance genes and their expression patterns in a similar phylogenetic framework could provide a broader understanding and prediction of resistance mechanisms across different bacterial species and antibiotics. Expression of a resistance gene like *bla*_TEM-1_ might carry a fitness penalty to the host cell. In the absence of antibiotics, this can select against cells with resistance gene-carrying plasmids. We speculate that in bacterial populations, conjugative plasmids could escape purging by moving from bacterial lineages with high intrinsic expression, which bear the fitness cost, to low intrinsic expression, where the fitness cost is reduced. This process could help maintain the presence of resistance genes within the population even in the absence of antibiotic pressure.

## Acknowledgements

The authors thank Jarrod Hadfield for providing guidance on implementing the MCMCglmm library.

