## Supplementary File 1 for "*E. coli* phylogeny drives co-amoxiclav resistance through variable expression of *bla*_TEM-1_"

**Genome assembly**

We first performed quality control on both the short- and long-read sets (see readQC.sh). For the short-reads, we ran fastp (v. 0.23.4) with default parameters to remove any unpaired short-reads. For the long-reads, we ran filtlong (v. 0.2.1) with default parameters except --min_length 1000 --keep_percent 95 .This removed any reads shorter than 1kbp and excluded the worst 5% of reads.

We then assembled the genomes using two methods:

1. **Flye.** We first built a long-read assembly graph with Flye (v. 2.9.2-b1786; default parameters except --plasmids --nano-raw; see runFlye.sh). Next, we polished the graph with long-reads using medaka (v. 1.8.0; default parameters except -m r941_min_high_g360). Medaka did not provide a model which perfectly matched our set-up (MinION with a R9.4.1 flow cell and Guppy [v. 3.84] in SUP mode), so we used r941_min_high_g360 as the closest match. Lastly, we polished the graph with short-reads. To do this, we first aligned our short-reads to the assembly with bwa (v. 0.7.17-r1188; index mode followed mem mode for both halves of the short-read sets; default parameters), then Polypolish (v. 0.5.0; polypolish_insert_filter.py followed by running with default parameters). Proceeding the polishing with a size filter excluded any excessive alignments. We polished with long-reads first because they are less likely to align to multiple positions. The short-read polishing then only made changes with unanimous agreement.
2. **Unicycler.** We produced a hybrid assembly with Unicycler (v. 0.5.0; see runUnicycler.sh). This method produces a short-read assembly graph with SPAdes (v. 3.15.5), and then uses the long-reads to bridge between contigs with miniasm (v. 0.3-r179) and Racon (v. 1.5.0). We ran Unicycler with default settings except --mode conservative min_component_size 500 --min_dead_end_size 500.

Next, we approximated the depth of the short- and long-reads by dividing the total size of each read set by 5Mbp, the average size of an *E. coli* genome (see approxDepth.py). Taking any value less than 50 to be ‘shallow’ and anything larger to be ‘deep’, 0.5% (3/548) had shallow short- and long-reads, 1.5% (8/548) had shallow short-reads and deep long-reads, 26.5% (145/548) has deep short-reads and shallow long-reads, and 71.5% (392/548) had deep short- and long-reads. We found that when short-reads were deep, regardless of long-read depth, the Unicycler method was better at recovering a “complete” assembly (all contigs circularised; 46.7% [251/537] for Unicycler versus 33.0% [177/537] for Flye), but the Flye method was better at circularising the chromosome (57.9% [311/537] for Unicycler versus 66.9% [359/537] for Flye).

For *n*=3 isolates (OXEC-60, OXEC-247, and OXEC-273), we had to subsample their long-reads due to memory constraints. Before, isolates OXEC-60, OXEC-247, and OXEC-273 had approximate long-read depths of 519, 485, and 561, respectively. We used Rasusa (v. 0.7.1) with default parameters except --coverage 200 --genome-size 5mb to reduce them to an approximate depth of 200.

Long-read-first assemblies are better at resolving repetitive regions in genomes than short-read-first assemblies. Also, for this study, having a complete assembly was preferable to only having a circularised chromosome. This determined our order of preference in the final assembly choice: 1^st^ choice a complete Flye assembly (177/548); 2^nd^ choice a complete Unicycler assembly (148/548); 3^rd^ choice a circularised-chromosome Flye assembly (85/548); 4^th^ choice a circularised-chromosome Unicycler assembly (22/548). In total, we kept 78.8% (432/548) of genome assemblies, discarding the remaining 21.2% (116/548).

**Dataset curation**

Firstly, we observed one isolate (OXEC-153) for which the assembly chromosome length was 3,560,788bp < 4.5Mbp, which we removed, leaving *n*=431 assemblies. Then, to identify *bla*_TEM-1_ and other AMR genes, all assemblies were annotated with NCBIAMRFinder (v. 3.11.26 and database v. 2023-11-15.1) with default parameters except --plus --organism Escherichia. Alongside, we validated the presence of *bla*_TEM-1_ using tblastn (v. 2.15.0+) with the NCBI Reference Gene Catalog TEM-1 RefSeq protein WP_000027057.1 and 100% amino acid identity. At this stage, *n*=42 assemblies were found to not carry *bla*_TEM-1_, and a further *n*=12 were found to carry additional beta-lactamases, leaving a total of *n*=377 assemblies in our final dataset.

**Validating plasmid contigs**

To confirm the origin of the non-chromosomal contigs, we used Mash screen (v. 2.3) to score their containment in plasmids from PLSDB (v. 2023_06_23_v2), a curated database of 50,554 plasmid sequences curated from NCBI (see screenPlsdb.sh). For each of our assembly contigs, we kept the top hit.
