## Supplementary File 2 for "*E. coli* phylogeny drives co-amoxiclav resistance through variable expression of *bla*_TEM-1_"

### **Single phenotype model specifications**

Both of the genotype-phenotype models are generalised linear models with mixed effects (GLMMs), structured as

$$\boldsymbol{Y}=X\boldsymbol{\beta}+\sum_{i} Z_{i}\boldsymbol{u}_{\boldsymbol{i}}+ \boldsymbol{\varepsilon}$$

where $\boldsymbol{Y}$ is the response vector, is the $X$ design matrix for the fixed effects, $\boldsymbol{\beta}$ is the fixed effects coefficients, $Z_{i}$ is the design matrix for the random effect $i$, is $\boldsymbol{u}_{\boldsymbol{i}}$ the random effects coefficients, and $\boldsymbol{\varepsilon}$ is the residual errors.

In the *bla*_TEM-1_ expression model, the response variable $\boldsymbol{Y}$ is assumed to follow a multivariate normal distribution

$$\boldsymbol{Y} \sim\text{N}(X\boldsymbol{\beta}+\Sigma_{i}Z_{i}\boldsymbol{u}_{\boldsymbol{i}}, \sigma^{2}I)$$

where$\sigma^{2}$ is the residual variance. The fixed effects are assumed to follow a multivariate normal distribution

$$\boldsymbol{\beta} \sim\text{N}(\boldsymbol{\mu}_{\boldsymbol{\beta}}, \boldsymbol{\sigma}_{\boldsymbol{\beta}}^{\boldsymbol{2}}\boldsymbol{)}$$

The random effects for qPCR replicates are given by ${Z_{1}\boldsymbol{u}}_{\boldsymbol{1}}$ and

$$\boldsymbol{u}_{\boldsymbol{1}} \sim\text{N}\left( \boldsymbol{0}, \sigma_{u_{1}}^{2}I \right)$$

The phylogenetic random effects are given by ${Z_{2}\boldsymbol{u}}_{\boldsymbol{2}}$ and

$$\boldsymbol{u}_{\boldsymbol{2}} \sim\text{N}\left( \boldsymbol{0}, \sigma_{u_{2}}^{2}A^{-1} \right)$$

where$A^{-1}$ is the inverse of the phylogenetic relationship matrix (see later). The residuals $\boldsymbol{\varepsilon}$ are assumed to follow a multivariate normal distribution

$$\boldsymbol{\varepsilon} \sim\text{N}(\boldsymbol{0}, \sigma^{2}I)$$

The co-amoxiclav MIC model is specified similarly, except there is no random effect for qPCR replicate, and $\boldsymbol{Y}$ is assumed to follow an ordinal distribution modelled through underlying continuous latent variables

$$Y_{i}^{*} \sim\text{N}(\mu_{i}, \sigma^{2})$$

and the observed ordinal response $Y_{i}$ is determined by cutpoints $\theta_{i}$ applied to $Y_{i}^{*}$

$$Y_{i}= \left\{ \begin{aligned} &1\text{ if }Y_{i}^{*} \leq\theta_{1} \\ &2\text{ if }\theta_{1} < Y_{i}^{*} \leq\theta_{2} \\ &\vdots\\ &k \text{if} \theta_{k-1}< Y_{i}^{*} \end{aligned} \right.$$

### **Parameter estimation**

We used Markov Chain Monte Carlo (MCMC) to sample posterior distributions for the fixed effects, random effects, variance components, and for the ordinal model, cutpoints. Fixed effects use normal priors $\text{N}\left( \boldsymbol{0}, {10}^{10}\boldsymbol{\times I} \right)$. Priors for the variance components of qPCR replicate random effect and residual errors were $\text{Inverse-Wishart}(1, 0.02)$. For the phylogenetic random effect variance, to improve the MCMC mixing, we defined it as the product $u_{i}=\alpha\eta_{i}$ where $\eta_{i}\sim\text{N}(0, V_{\eta})$, which yields the two priors $\alpha\sim\text{N}(0, V\alpha)$ and $V\eta\sim\text{Inverse-Gamma}(V, \nu)$. We set $\alpha\sim\text{N}(0, 1000)$ and $\text{Inverse-Gamma}\left( 0.001, 0.001 \right)$. In the co-amoxiclav MIC model, we fixed the residual variance at 1.

For both models, we ran two chains for 10 million iterations with 10% burn-in and a thinning interval of 100. To test for convergence, we calculated the Gelman-Rubin statistic for each model to confirm it was invariably 1 for all parameters. It assumes that if the chains have converged, the between-chain variance should be similar to the within-chain variance. We also verified that effective sample sizes were comparable between chains. We calculated the autocorrelation function for all chains and visually inspected the trace plots for global trends. For all parameters, we calculated the posterior means and 95% high density intervals.

Parameter values reported in the manuscript refer to the first chain.

For a parameter $\beta$, $p_{\text{MCMC}}\left( \beta\right)=2 \cdot\text{min}\left( \mathbb{P}\left( \beta>0 \right)\mathbb{, P}\left( \beta<0 \right) \right)$ and measures the probability that $\beta$ is in the more extreme tail of its posterior distribution. If $\beta$ were truly centred around zero, we would expect $\mathbb{P}\left( \beta>0 \right)\mathbb{\approx P}\left( \beta<0 \right)\approx0.5$.

### **Phylogeny as a random effect**

Here we assume that more closely related isolates have similar responses due to their shared evolutionary history. With the *E. coli* chromosomal phylogeny, we took the midpoint root, then coerced the tree into ultrametricity using a penalised likelihood method which assumed that assumed that rates of branch evolution were correlated. Then, taking the transformed phylogeny as a variance-covariance matrix $A$, we inverted it to generate the precision matrix $A^{-1}$. Using the precision matrix is generally more numerically stable.

Best Linear Unbiased Predictors (BLUPs) represent the deviation of individual isolates from the average effect across the entire phylogeny. The total variance of all BLUPs is the phylogenetic random effect variance.

### ***bla*_TEM-1_ expression model outputs**

```
> summary(chain.1)

 Iterations = 1000001:9999901
 Thinning interval  = 100
 Sample size  = 90000

 DIC: -84.31782

 G-structure:  ~phylo

      post.mean  l-95% CI u-95% CI eff.samp
phylo   0.06803 6.734e-12   0.2066    48985

               ~isolate.assembly

                 post.mean l-95% CI u-95% CI eff.samp
isolate.assembly    0.1498  0.07779   0.2272    76382

 R-structure:  ~units

      post.mean l-95% CI u-95% CI eff.samp
units   0.03356  0.02711  0.04045    90000

 Location effects: exp.scaled ~ pos1.bool * pos2.bool + contig.copy.number.scaled

                            post.mean  l-95% CI  u-95% CI eff.samp  pMCMC
(Intercept)                  0.297258  0.031125  0.585104    87300 0.0245 *
pos1.boolTRUE               -1.713268 -2.079701 -1.343259    90000 <1e-05 ***
pos2.boolTRUE               -0.313958 -0.570731 -0.042875    90000 0.0202 *
contig.copy.number.scaled   -0.116605 -0.242749  0.007042    87138 0.0679 .
pos1.boolTRUE:pos2.boolTRUE  0.325914 -0.143755  0.811333    88934 0.1779
---
Signif. codes:  0 ‘***’ 0.001 ‘**’ 0.01 ‘*’ 0.05 ‘.’ 0.1 ‘ ’ 1

> autocorr.diag(chain.1$VCV)
               phylo isolate.assembly         units
Lag 0    1.000000000     1.0000000000  1.0000000000
Lag 100  0.147970235     0.0373112895  0.0019415565
Lag 500  0.027018030     0.0048930316 -0.0007288071
Lag 1000 0.005572987    -0.0014661450  0.0003393882
Lag 5000 0.000294595     0.0007567522  0.0024936229

> plot(chain.1)
```

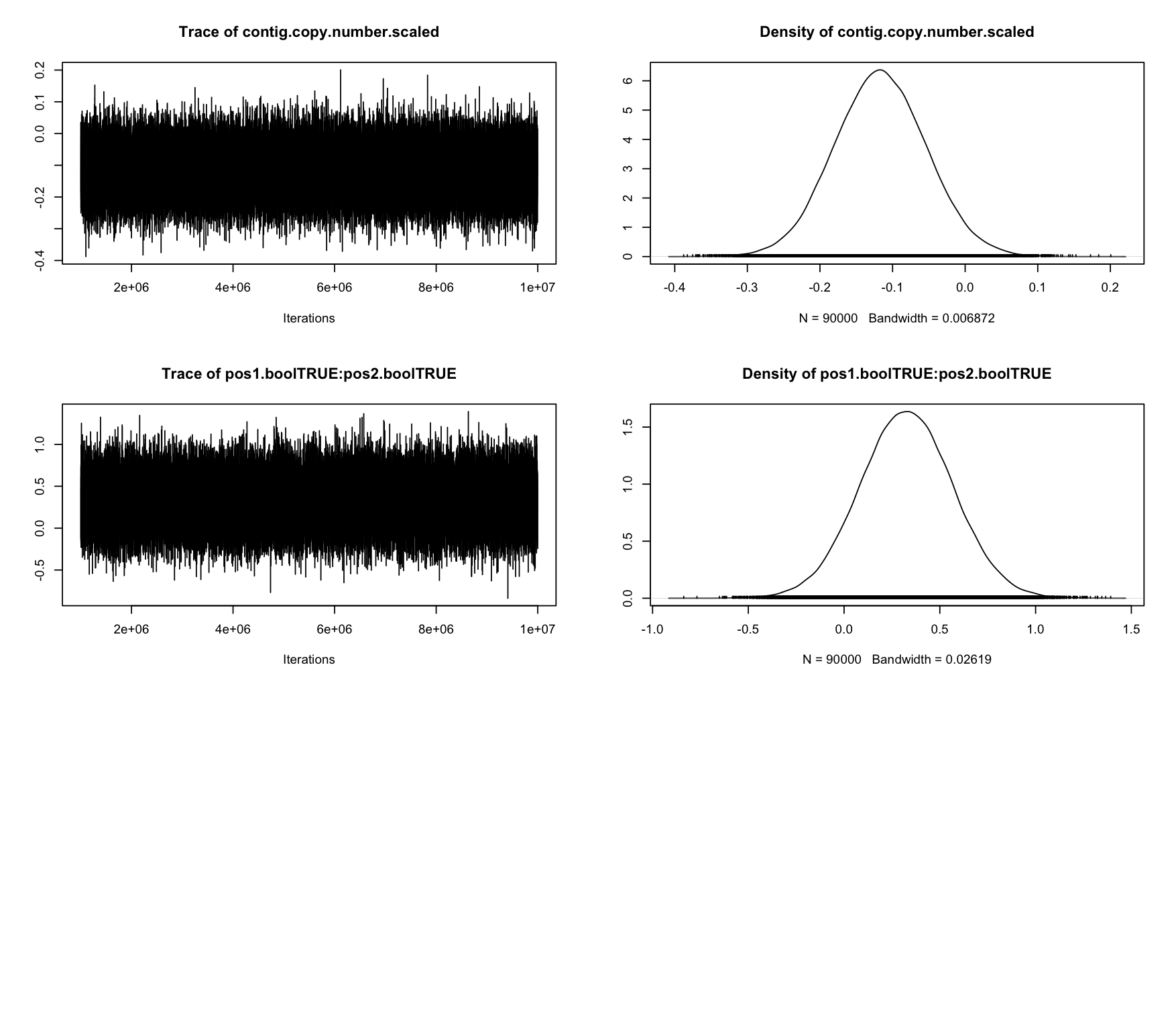

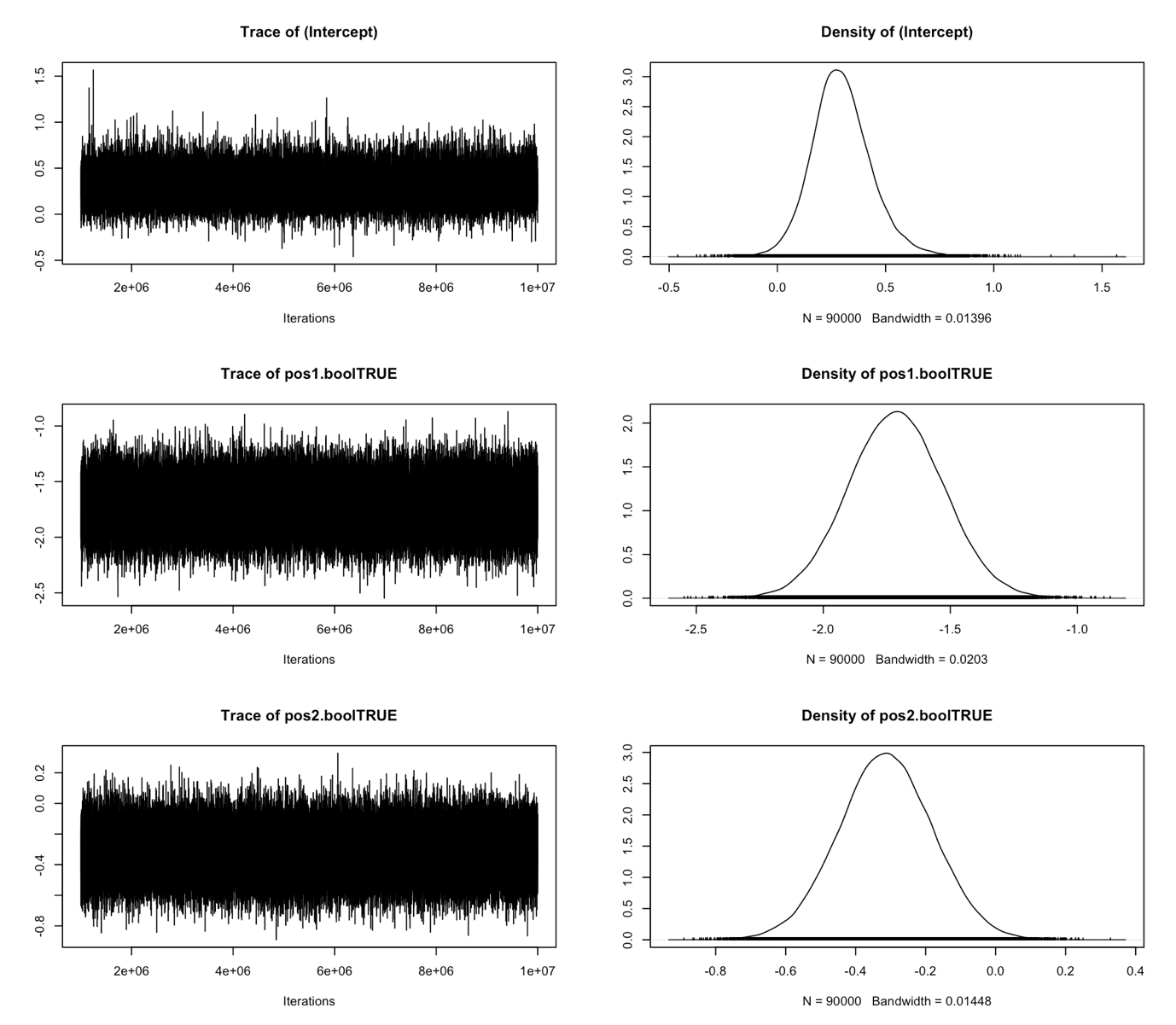

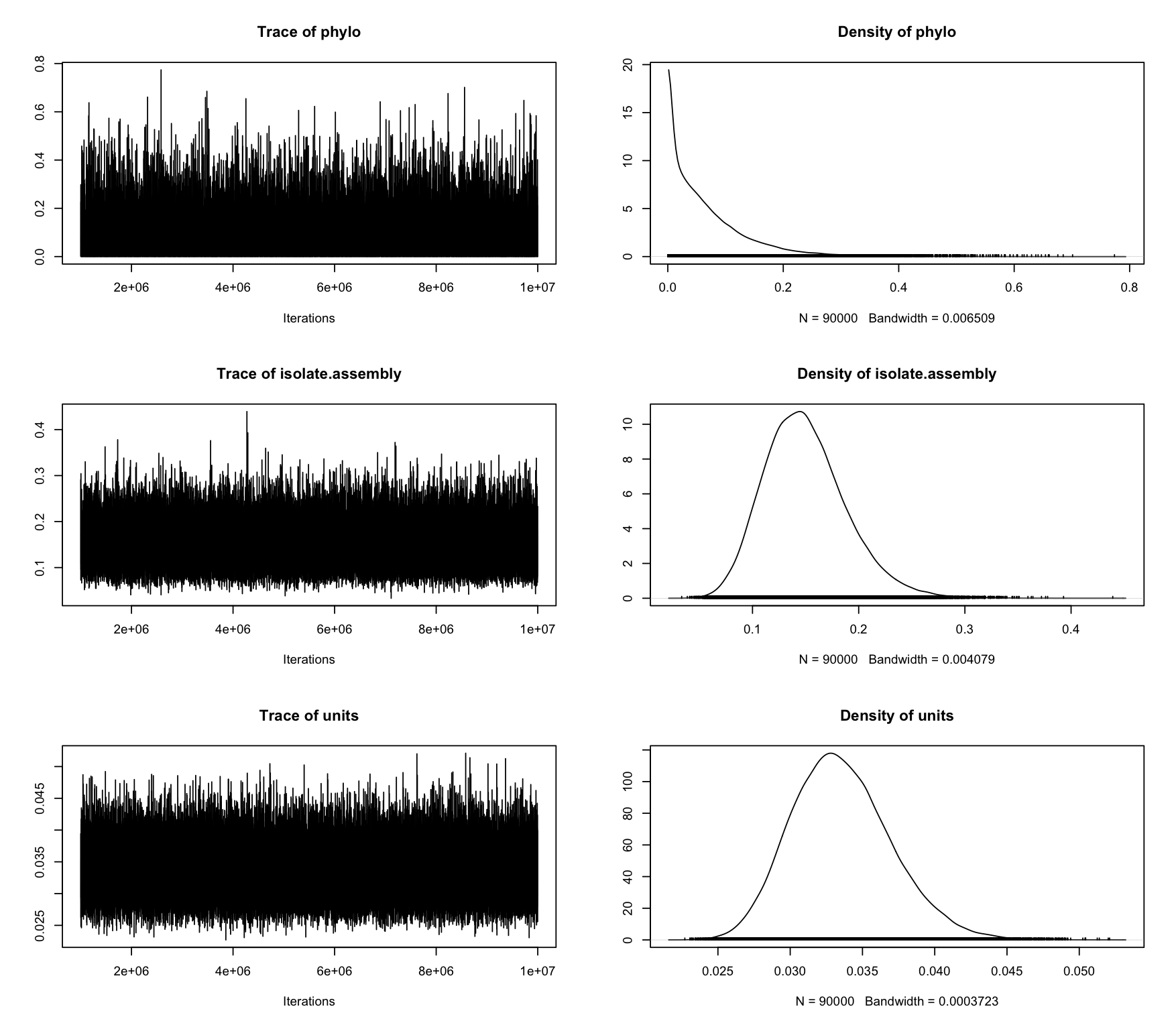
> summary(chain.2)

```
 Iterations = 1000001:9999901
 Thinning interval  = 100
 Sample size  = 90000

 DIC: -84.35054

 G-structure:  ~phylo

      post.mean  l-95% CI u-95% CI eff.samp
phylo   0.06815 3.565e-10   0.2062    48048

               ~isolate.assembly

                 post.mean l-95% CI u-95% CI eff.samp
isolate.assembly    0.1499  0.07934   0.2281    74765

 R-structure:  ~units

      post.mean l-95% CI u-95% CI eff.samp
units   0.03355   0.0272  0.04045    91734

 Location effects: exp.scaled ~ pos1.bool * pos2.bool + contig.copy.number.scaled

                            post.mean  l-95% CI  u-95% CI eff.samp  pMCMC
(Intercept)                  0.296815  0.038728  0.590741    88946 0.0247 *
pos1.boolTRUE               -1.713476 -2.083985 -1.344730    90000 <1e-05 ***
pos2.boolTRUE               -0.313962 -0.585371 -0.054027    90000 0.0210 *
contig.copy.number.scaled   -0.117043 -0.242901  0.008085    87127 0.0684 .
pos1.boolTRUE:pos2.boolTRUE  0.326406 -0.146791  0.812865    90000 0.1787
---
Signif. codes:  0 ‘***’ 0.001 ‘**’ 0.01 ‘*’ 0.05 ‘.’ 0.1 ‘ ’ 1

> autocorr.diag(chain.2$VCV)
               phylo isolate.assembly         units
Lag 0    1.000000000     1.0000000000  1.0000000000
Lag 100  0.144760011     0.0425125177 -0.0040162838
Lag 500  0.027608837     0.0107320749  0.0041080909
Lag 1000 0.007409151    -0.0038731455 -0.0026025067
Lag 5000 0.002530502     0.0006590854  0.0004411566

> plot(chain.2)
```

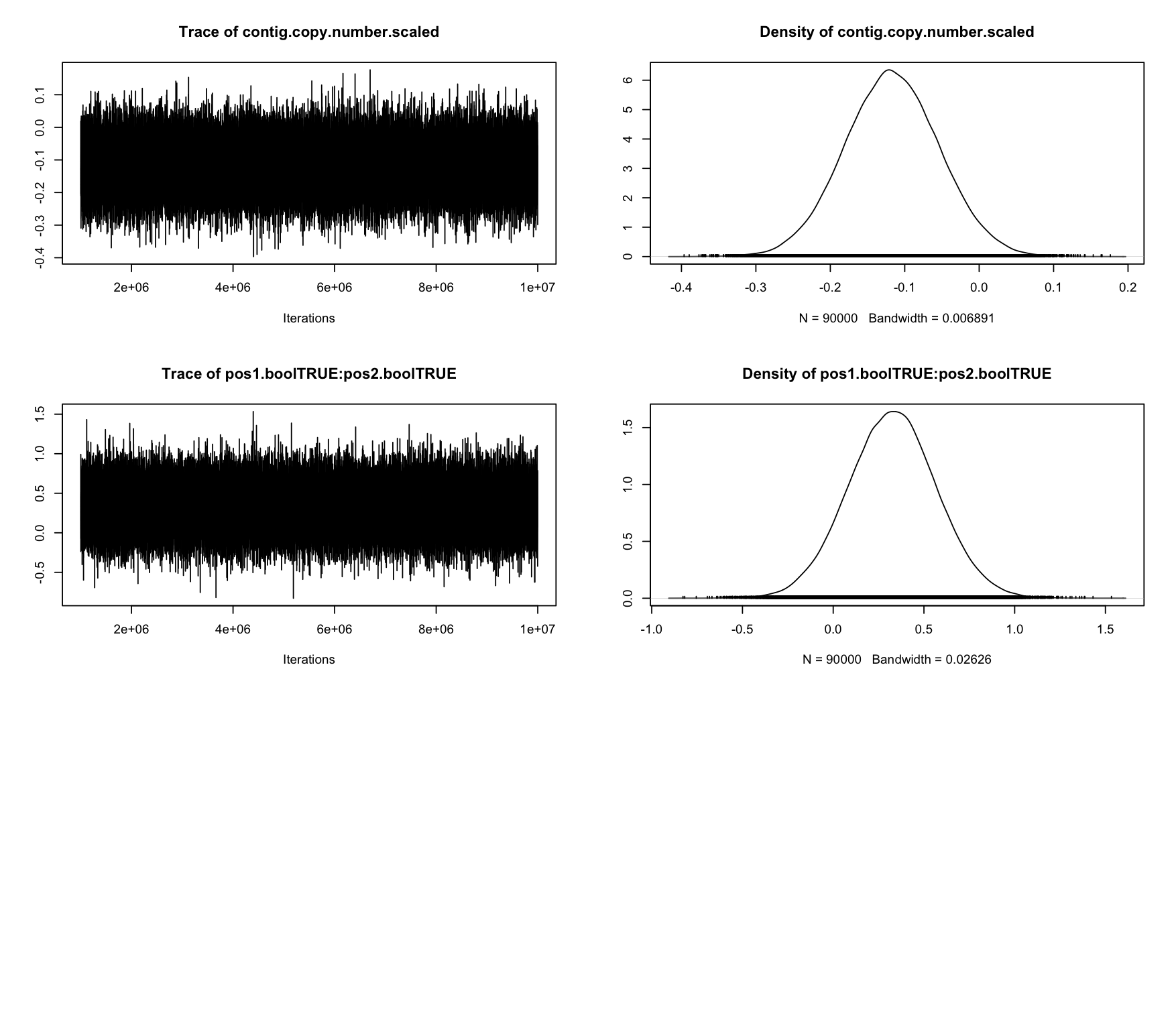

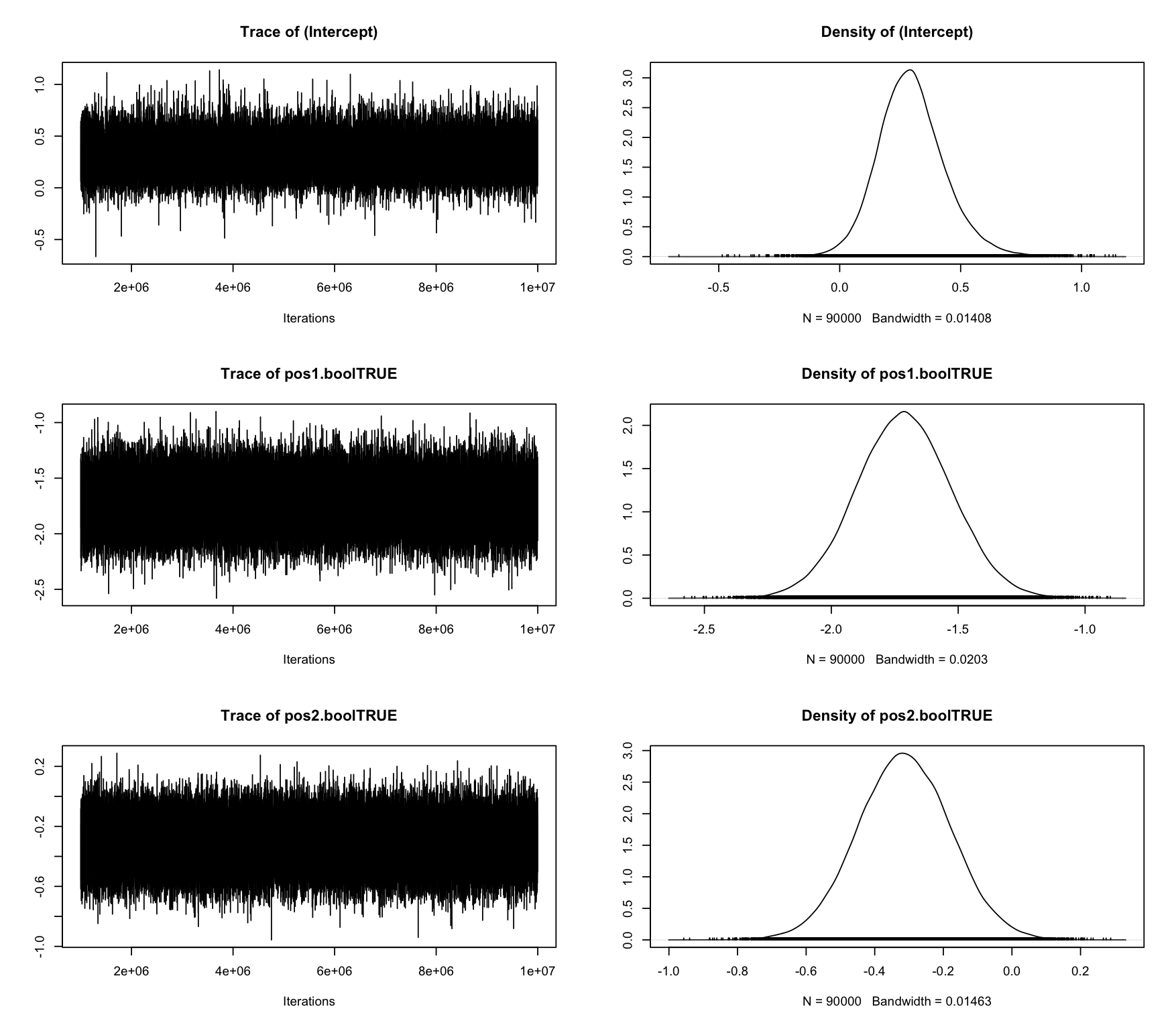

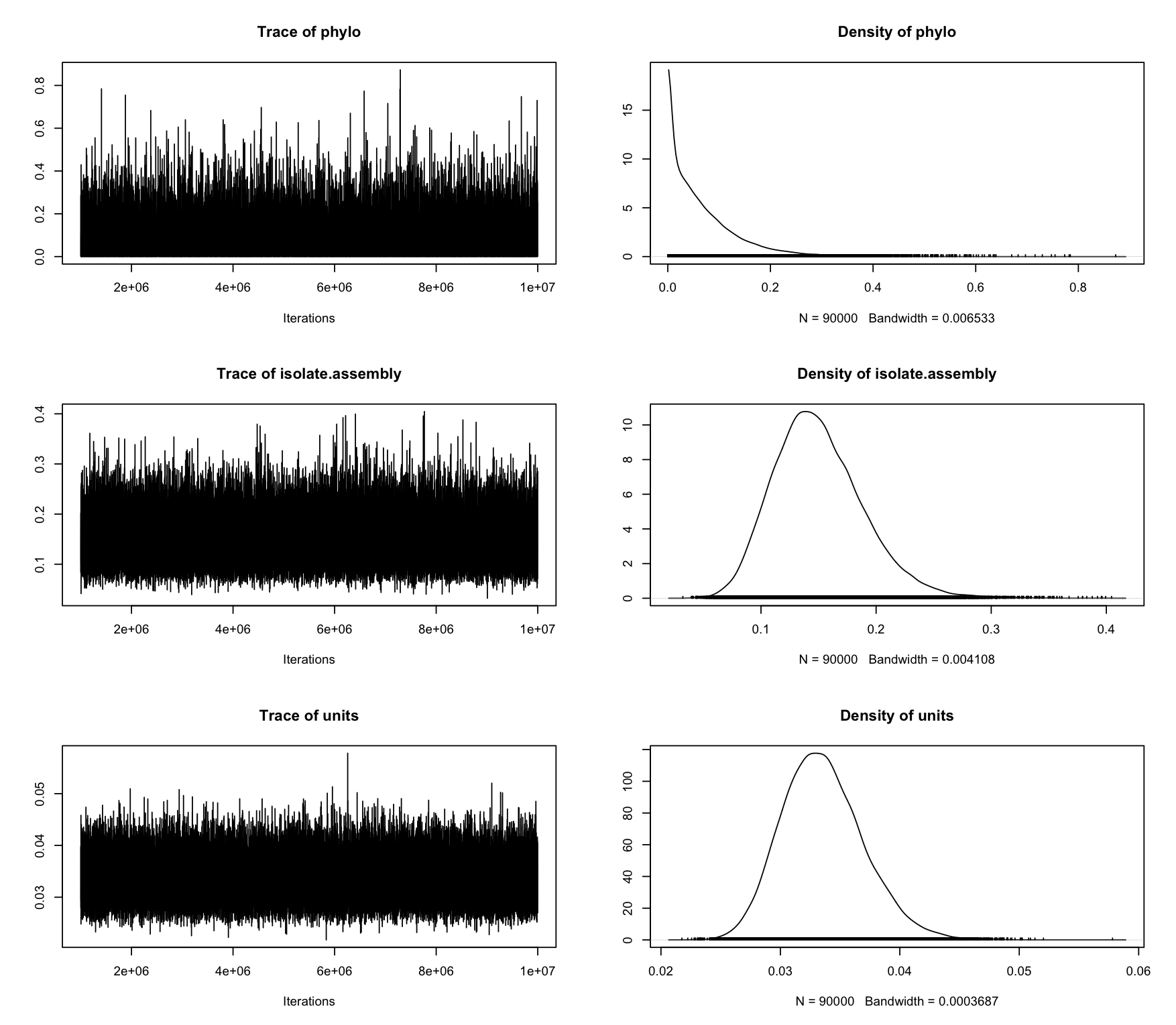
> mclist <- mcmc.list(chain.1$Sol, chain.2$Sol)

```
> gelman.diag(mclist)
Potential scale reduction factors:

                            Point est. Upper C.I.
(Intercept)                          1          1
pos1.boolTRUE                        1          1
pos2.boolTRUE                        1          1
contig.copy.number.scaled            1          1
pos1.boolTRUE:pos2.boolTRUE          1          1
phylo.542                            1          1
phylo.414                            1          1
phylo.236                            1          1
phylo.415                            1          1
phylo.476                            1          1
phylo.257                            1          1
phylo.103                            1          1
phylo.285                            1          1
phylo.284                            1          1
phylo.410                            1          1
phylo.344                            1          1
phylo.429                            1          1
phylo.322                            1          1
phylo.262                            1          1
phylo.387                            1          1
phylo.158                            1          1
phylo.365                            1          1
phylo.267                            1          1
phylo.477                            1          1
phylo.197                            1          1
phylo.41                             1          1
phylo.408                            1          1
phylo.292                            1          1
phylo.1                              1          1
phylo.480                            1          1
phylo.423                            1          1
phylo.247                            1          1
phylo.353                            1          1
phylo.532                            1          1
phylo.203                            1          1
phylo.534                            1          1
phylo.192                            1          1
phylo.334                            1          1
phylo.486                            1          1
phylo.541                            1          1
phylo.462                            1          1
phylo.93                             1          1
phylo.230                            1          1
phylo.457                            1          1
phylo.176                            1          1
phylo.75                             1          1
phylo.296                            1          1
phylo.442                            1          1
phylo.495                            1          1
phylo.116                            1          1
phylo.233                            1          1
phylo.83                             1          1
phylo.508                            1          1
phylo.386                            1          1
phylo.59                             1          1
phylo.490                            1          1
phylo.125                            1          1
phylo.69                             1          1
phylo.340                            1          1
phylo.25                             1          1
phylo.546                            1          1
phylo.279                            1          1
phylo.485                            1          1
phylo.38                             1          1
phylo.238                            1          1
phylo.544                            1          1
phylo.314                            1          1
phylo.345                            1          1
phylo.73                             1          1
phylo.190                            1          1
phylo.107                            1          1
phylo.440                            1          1
isolate.assembly.1                   1          1
isolate.assembly.25                  1          1
isolate.assembly.38                  1          1
isolate.assembly.41                  1          1
isolate.assembly.59                  1          1
isolate.assembly.69                  1          1
isolate.assembly.73                  1          1
isolate.assembly.75                  1          1
isolate.assembly.83                  1          1
isolate.assembly.93                  1          1
isolate.assembly.103                 1          1
isolate.assembly.107                 1          1
isolate.assembly.116                 1          1
isolate.assembly.125                 1          1
isolate.assembly.158                 1          1
isolate.assembly.176                 1          1
isolate.assembly.190                 1          1
isolate.assembly.192                 1          1
isolate.assembly.197                 1          1
isolate.assembly.203                 1          1
isolate.assembly.230                 1          1
isolate.assembly.233                 1          1
isolate.assembly.236                 1          1
isolate.assembly.238                 1          1
isolate.assembly.247                 1          1
isolate.assembly.257                 1          1
isolate.assembly.262                 1          1
isolate.assembly.267                 1          1
isolate.assembly.279                 1          1
isolate.assembly.284                 1          1
isolate.assembly.285                 1          1
isolate.assembly.292                 1          1
isolate.assembly.296                 1          1
isolate.assembly.314                 1          1
isolate.assembly.322                 1          1
isolate.assembly.334                 1          1
isolate.assembly.340                 1          1
isolate.assembly.344                 1          1
isolate.assembly.345                 1          1
isolate.assembly.353                 1          1
isolate.assembly.365                 1          1
isolate.assembly.386                 1          1
isolate.assembly.387                 1          1
isolate.assembly.408                 1          1
isolate.assembly.410                 1          1
isolate.assembly.414                 1          1
isolate.assembly.415                 1          1
isolate.assembly.423                 1          1
isolate.assembly.429                 1          1
isolate.assembly.440                 1          1
isolate.assembly.442                 1          1
isolate.assembly.457                 1          1
isolate.assembly.462                 1          1
isolate.assembly.476                 1          1
isolate.assembly.477                 1          1
isolate.assembly.480                 1          1
isolate.assembly.485                 1          1
isolate.assembly.486                 1          1
isolate.assembly.490                 1          1
isolate.assembly.495                 1          1
isolate.assembly.508                 1          1
isolate.assembly.532                 1          1
isolate.assembly.534                 1          1
isolate.assembly.541                 1          1
isolate.assembly.542                 1          1
isolate.assembly.544                 1          1
isolate.assembly.546                 1          1

Multivariate psrf

1
```

### **Co-amoxiclav MIC model outputs**

```
> summary(chain.1)

 Iterations = 1000001:9999901
 Thinning interval  = 100
 Sample size  = 90000

 DIC: NaN

 G-structure:  ~phylo

      post.mean l-95% CI u-95% CI eff.samp
phylo     2.848   0.7516    5.247    68960

 R-structure:  ~units

      post.mean l-95% CI u-95% CI eff.samp
units         1        1        1        0

 Location effects: coamox.mic ~ tem1.isolate.copy.number.scaled + tem1.isolate.scaled + ampc.promoter.snv + promoter.snv + acrf

                                post.mean l-95% CI u-95% CI eff.samp  pMCMC
(Intercept)                       3.88700  2.88511  4.88655    68180 <1e-05 ***
tem1.isolate.copy.number.scaled   2.00863  1.34212  2.68362    87781 <1e-05 ***
tem1.isolate.scaledTRUE           0.99087  0.03854  1.92603    90000 0.0399 *
ampc.promoter.snvAGCTTCTAGGG      0.43731 -0.61580  1.50588    90000 0.4096
ampc.promoter.snvAGCTCCTAGGG      0.86638 -0.92864  2.59683    90000 0.3152
ampc.promoter.snvGATTCCTAGGG      0.83345 -1.44783  3.06506    90000 0.4444
promoter.snvCGGCGA                0.16276 -0.36927  0.67495    90862 0.5435
promoter.snvTGGCGA                6.05584  4.14118  8.06071    74795 <1e-05 ***
promoter.snvTGGCGG                5.87888  3.90570  7.93776    80057 <1e-05 ***
acrfTRUE                          0.01684 -0.50850  0.53567    90000 0.9475
---
Signif. codes:  0 ‘***’ 0.001 ‘**’ 0.01 ‘*’ 0.05 ‘.’ 0.1 ‘ ’ 1

 Cutpoints:

                           post.mean l-95% CI u-95% CI eff.samp
cutpoint.traitcoamox.mic.1     1.225   0.7772    1.710    47669
cutpoint.traitcoamox.mic.2     4.069   3.5440    4.556    31875
cutpoint.traitcoamox.mic.3     5.621   5.0860    6.113    30912
cutpoint.traitcoamox.mic.4     6.661   6.1760    7.000    32452

> autocorr.diag(chain.1$VCV)
                 phylo units
Lag 0     1.0000000000   NaN
Lag 100   0.0819682926   NaN
Lag 500   0.0075573098   NaN
Lag 1000  0.0008024054   NaN
Lag 5000 -0.0064707638   NaN

> plot(chain.1)
```

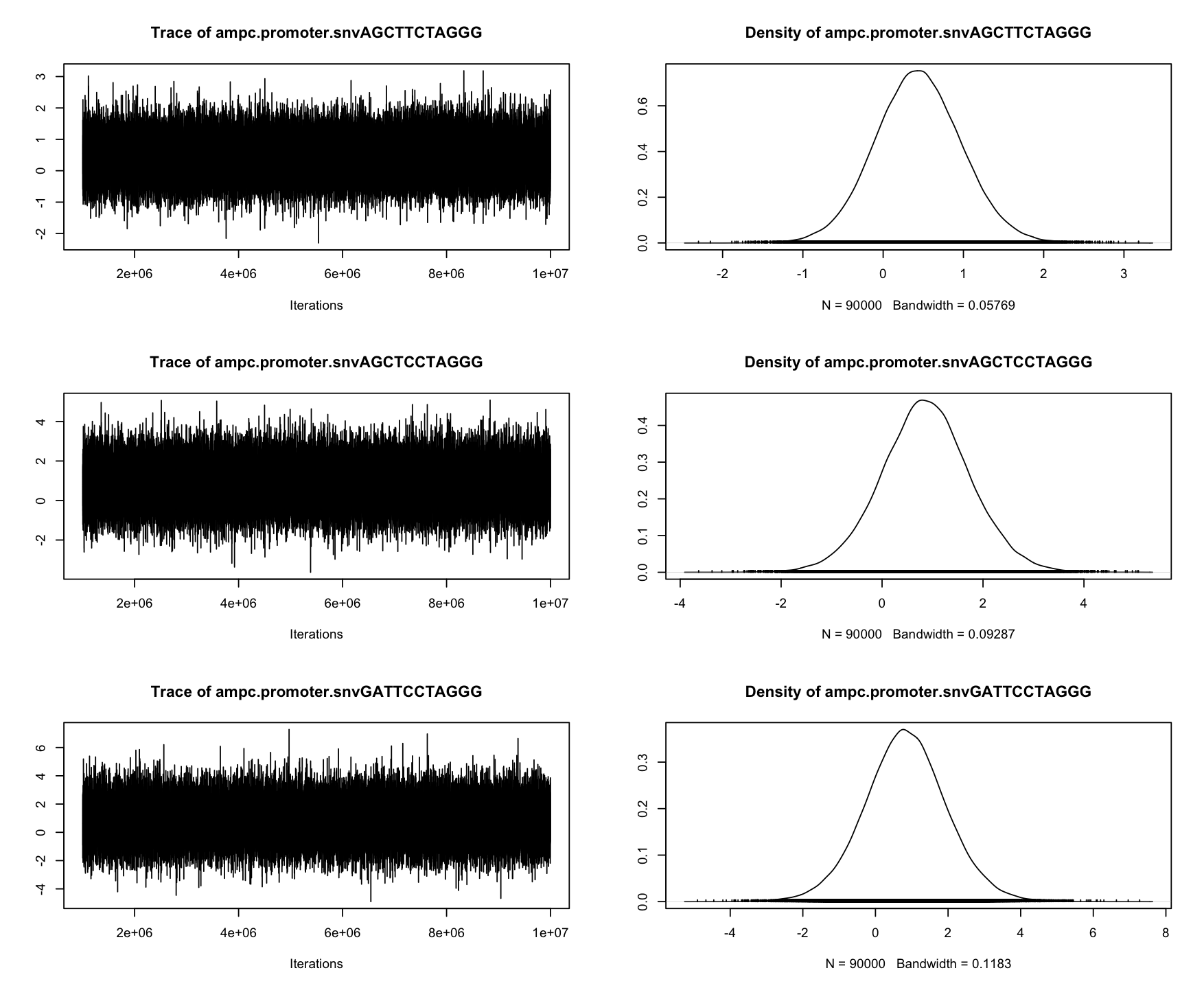

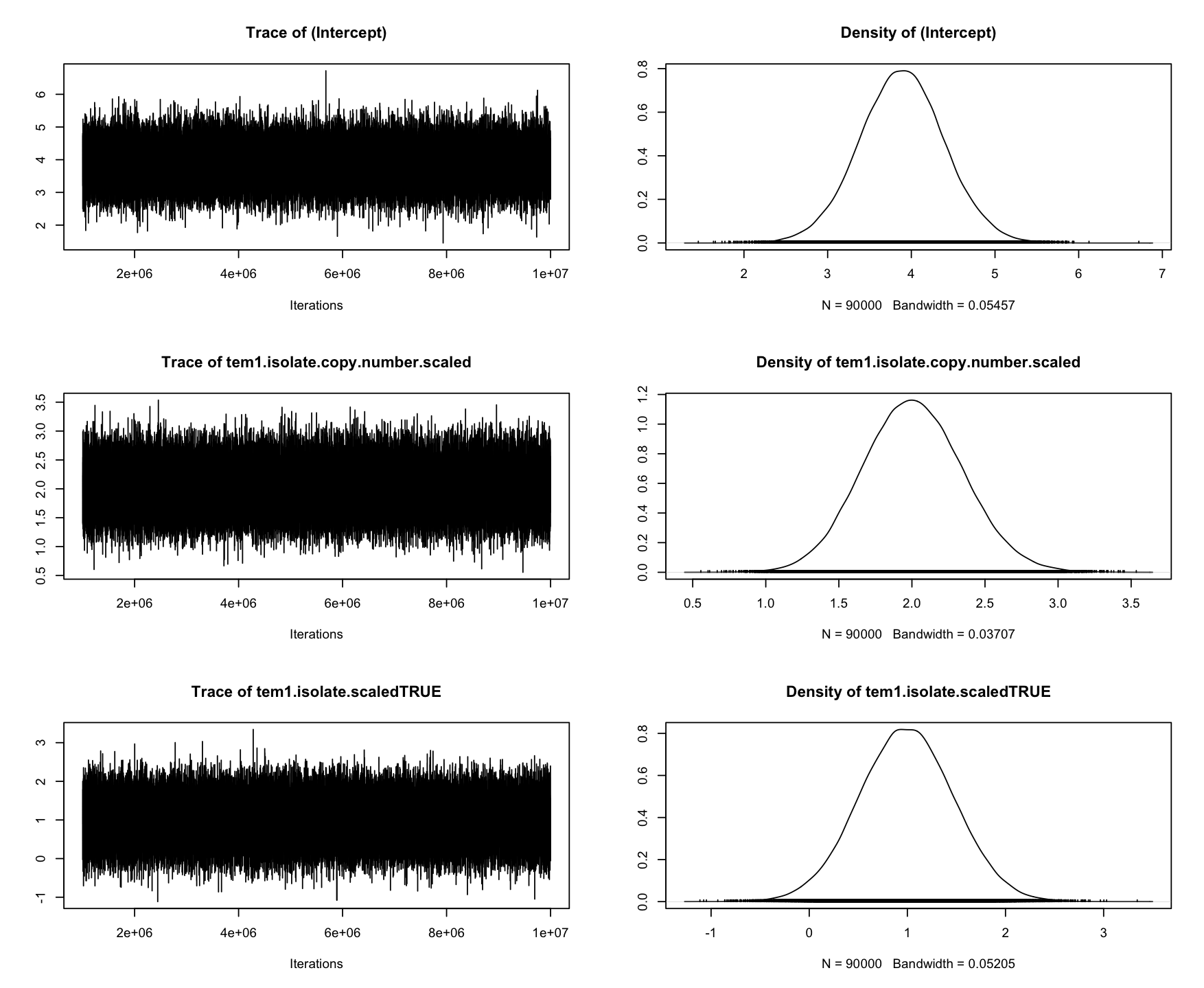

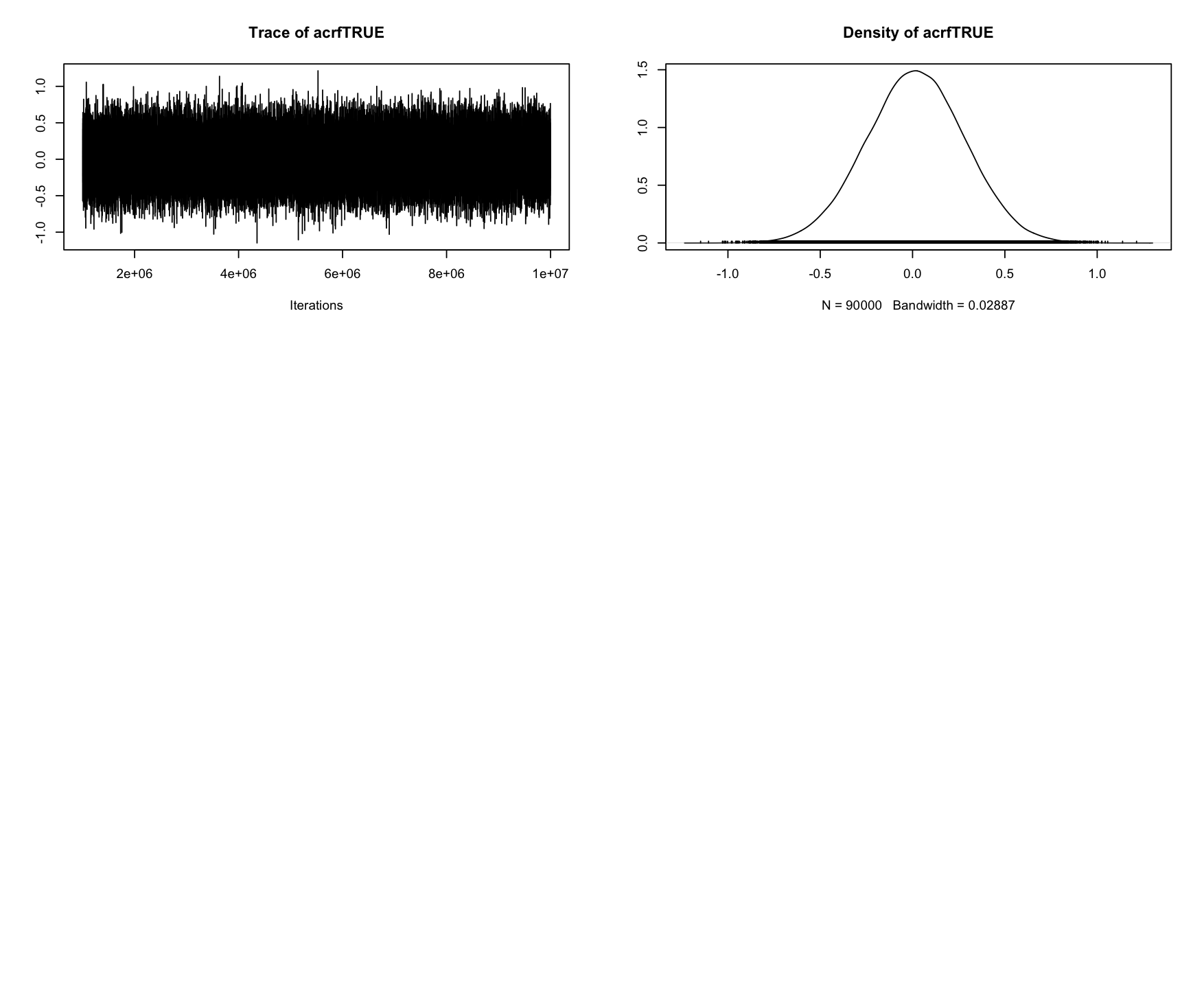

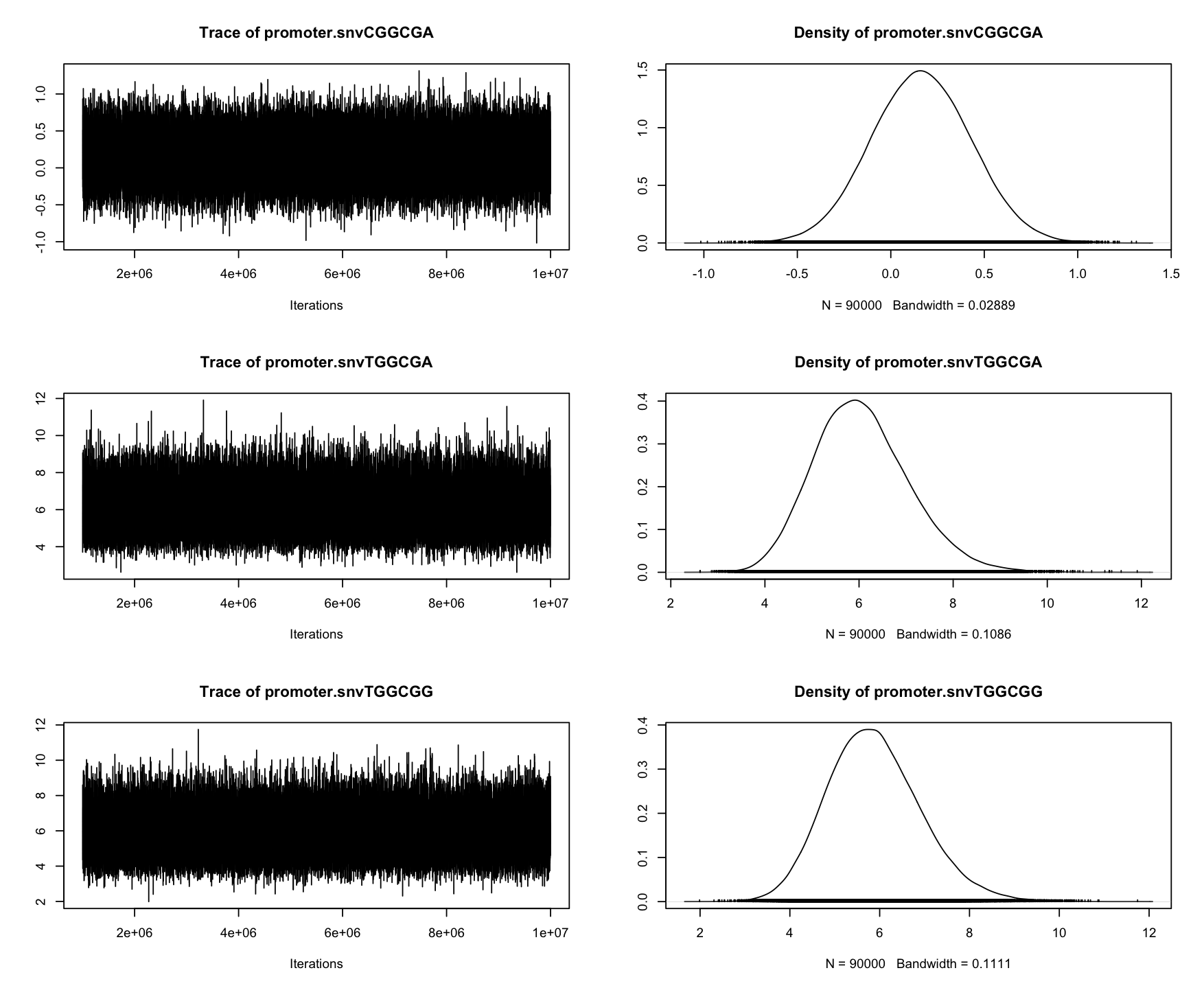

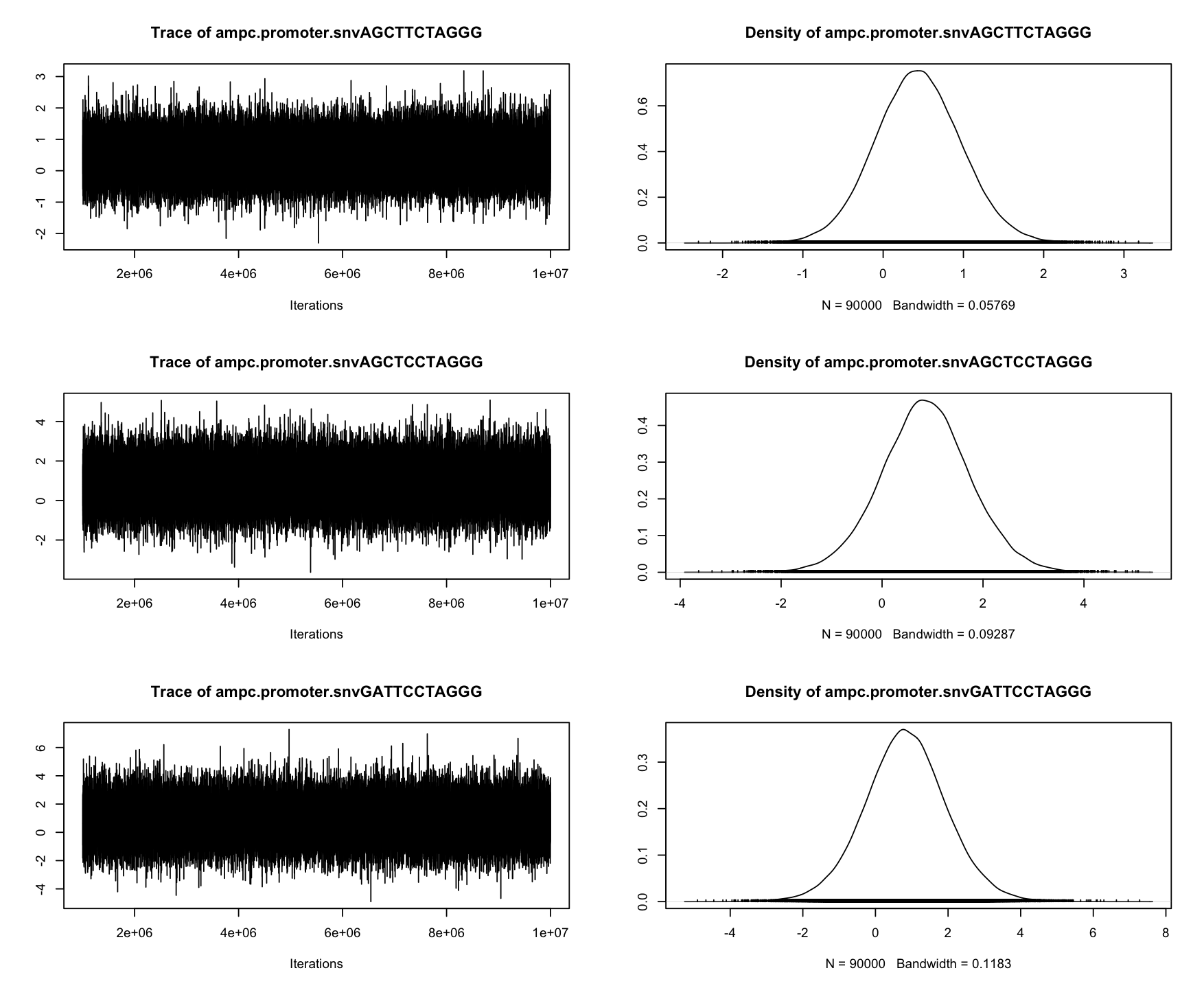

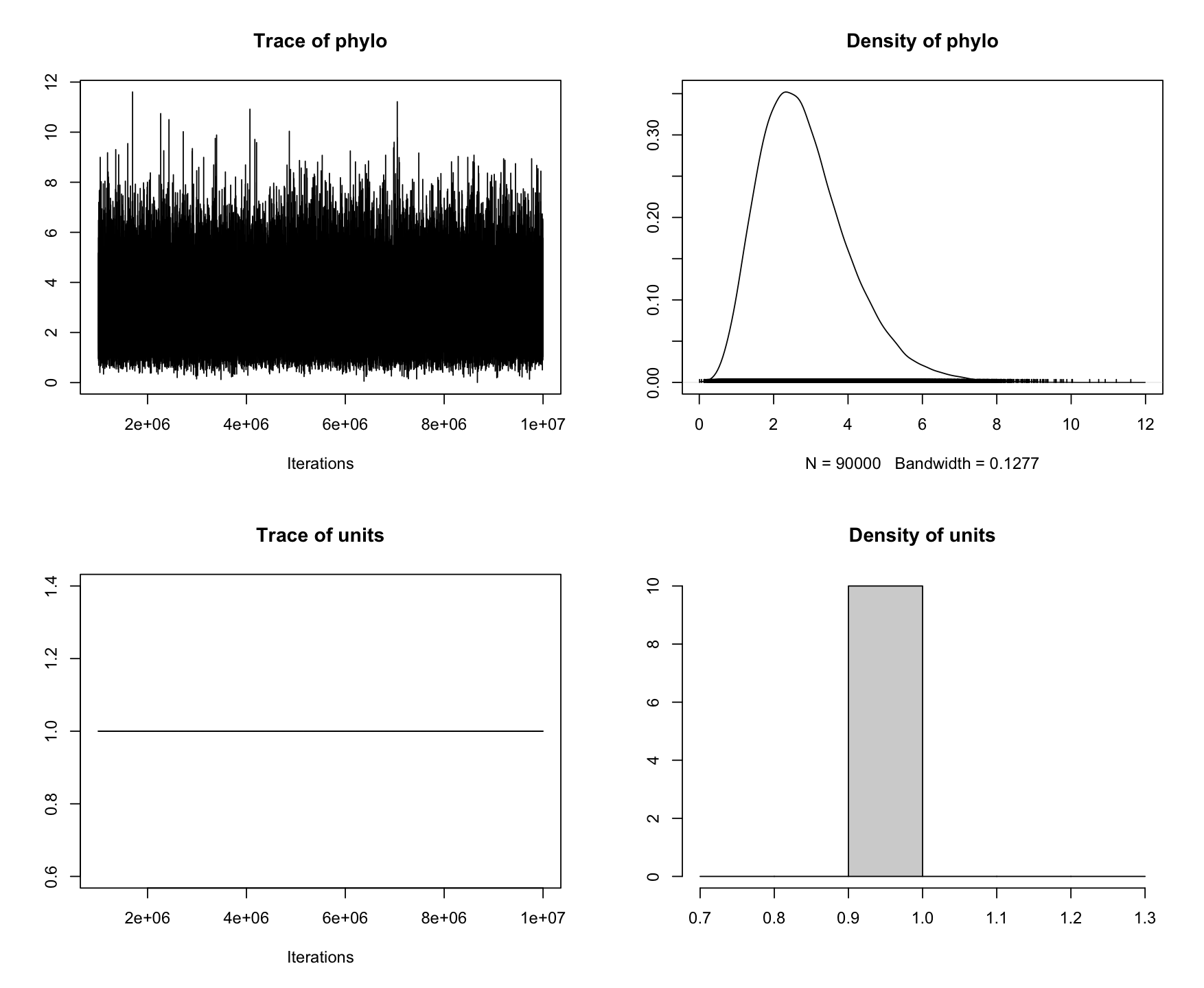

```
> summary(chain.2)

 Iterations = 1000001:9999901
 Thinning interval  = 100
 Sample size  = 90000

 DIC: NaN

 G-structure:  ~phylo

      post.mean l-95% CI u-95% CI eff.samp
phylo     2.827   0.7394    5.224    69909

 R-structure:  ~units

      post.mean l-95% CI u-95% CI eff.samp
units         1        1        1        0

 Location effects: coamox.mic ~ tem1.isolate.copy.number.scaled + tem1.isolate.scaled + ampc.promoter.snv + promoter.snv + acrf

                                post.mean l-95% CI u-95% CI eff.samp  pMCMC
(Intercept)                       3.87568  2.86385  4.85827    67288 <1e-05 ***
tem1.isolate.copy.number.scaled   2.00702  1.34054  2.68732    87830 <1e-05 ***
tem1.isolate.scaledTRUE           0.98913  0.04740  1.92915    90000  0.039 *
ampc.promoter.snvAGCTTCTAGGG      0.43856 -0.61152  1.52046    86262  0.411
ampc.promoter.snvAGCTCCTAGGG      0.87258 -0.85639  2.64130    90000  0.310
ampc.promoter.snvGATTCCTAGGG      0.82590 -1.41254  3.06937    87704  0.448
promoter.snvCGGCGA                0.16208 -0.36062  0.68443    90000  0.549
promoter.snvTGGCGA                6.05363  4.10873  8.02603    77559 <1e-05 ***
promoter.snvTGGCGG                5.87131  3.90756  7.94880    79880 <1e-05 ***
acrfTRUE                          0.01815 -0.49602  0.54566    90000  0.942
---
Signif. codes:  0 ‘***’ 0.001 ‘**’ 0.01 ‘*’ 0.05 ‘.’ 0.1 ‘ ’ 1

 Cutpoints:

                           post.mean l-95% CI u-95% CI eff.samp
cutpoint.traitcoamox.mic.1     1.222   0.7598    1.683    47758
cutpoint.traitcoamox.mic.2     4.061   3.5445    4.561    30820
cutpoint.traitcoamox.mic.3     5.612   5.0617    6.102    30353
cutpoint.traitcoamox.mic.4     6.650   6.1637    7.000    32413

> autocorr.diag(chain.2$VCV)
                 phylo units
Lag 0     1.0000000000   NaN
Lag 100   0.0754184027   NaN
Lag 500   0.0086358261   NaN
Lag 1000 -0.0005974199   NaN
Lag 5000 -0.0014404314   NaN

> plot(chain.2)
```

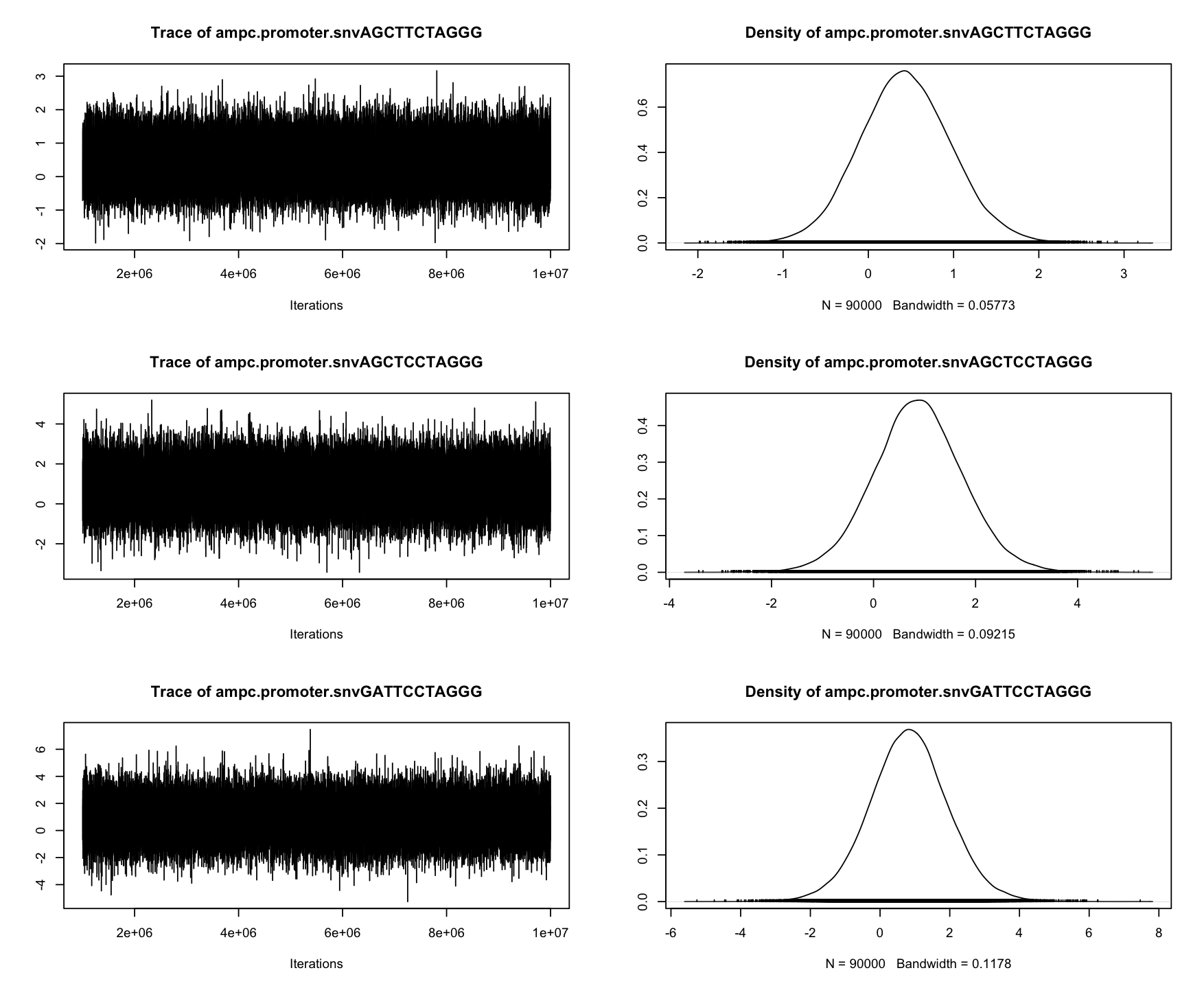

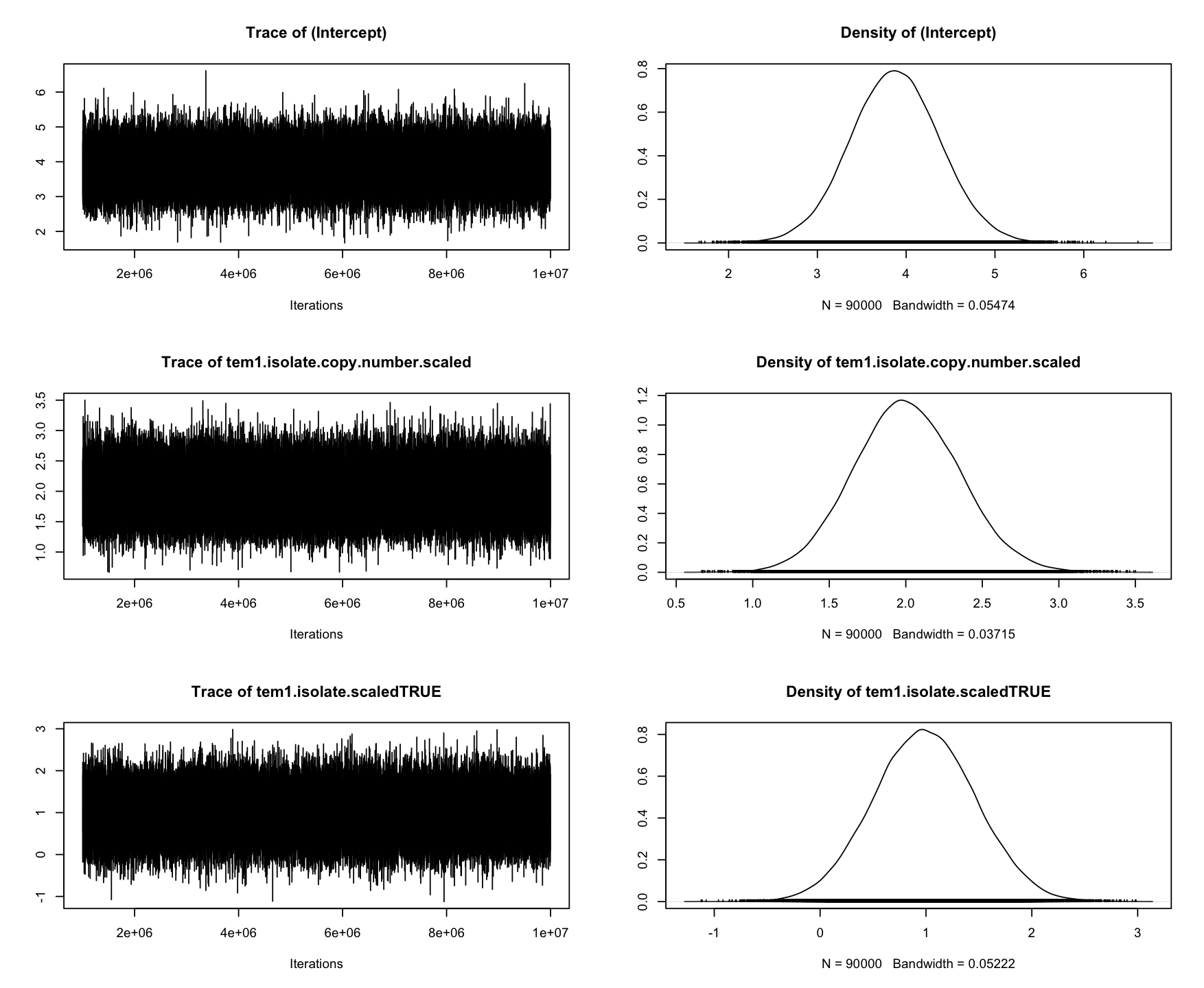

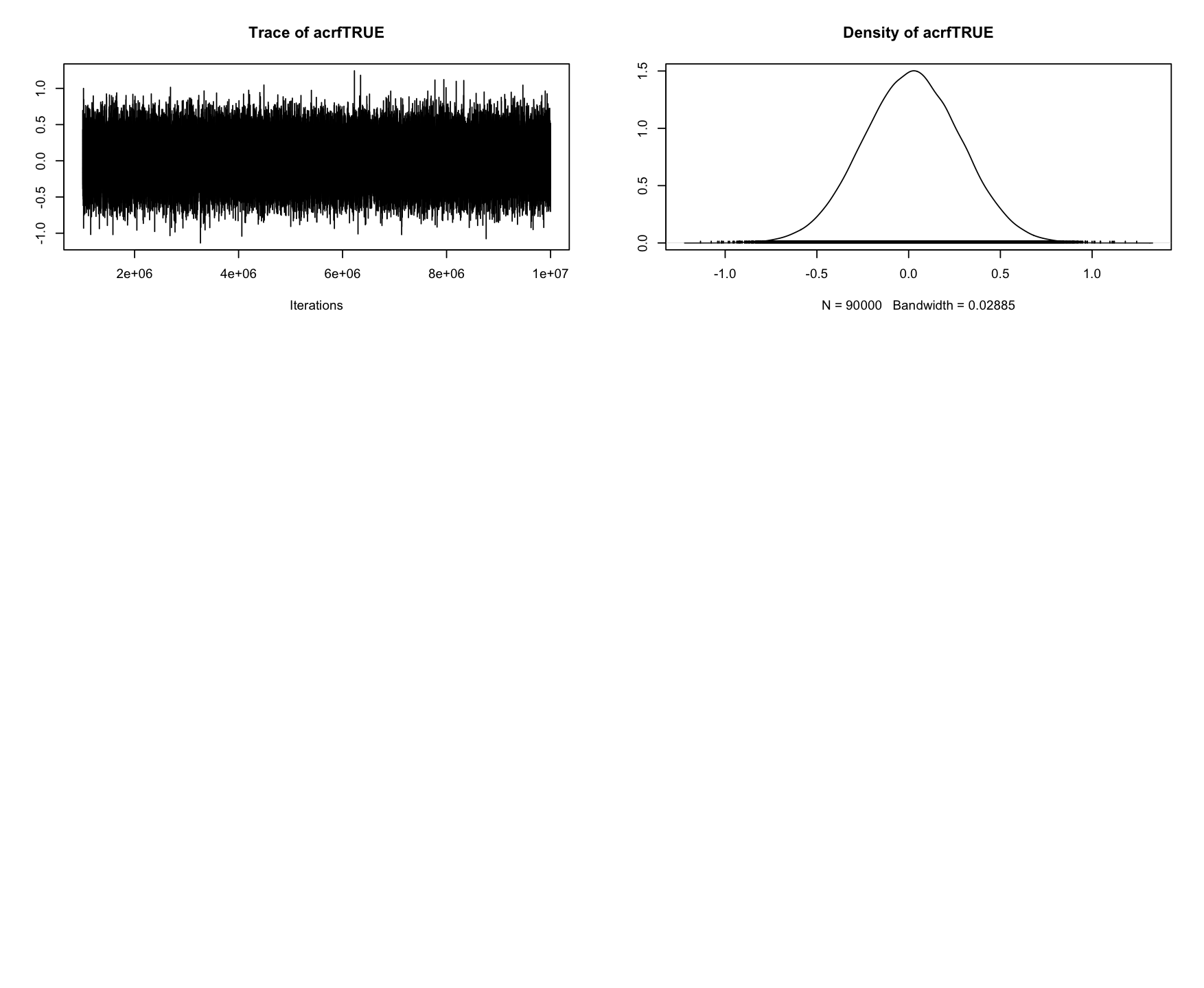

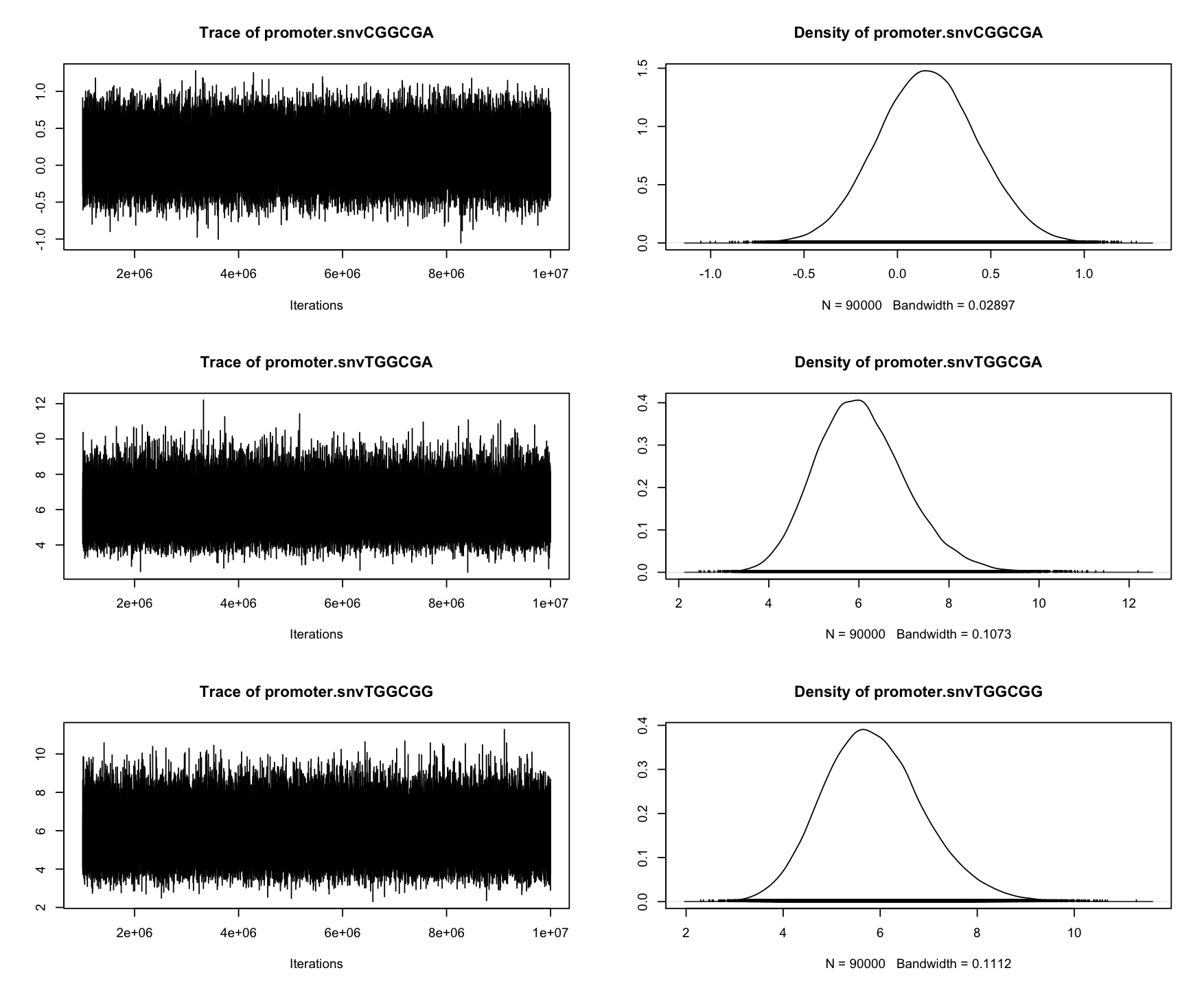

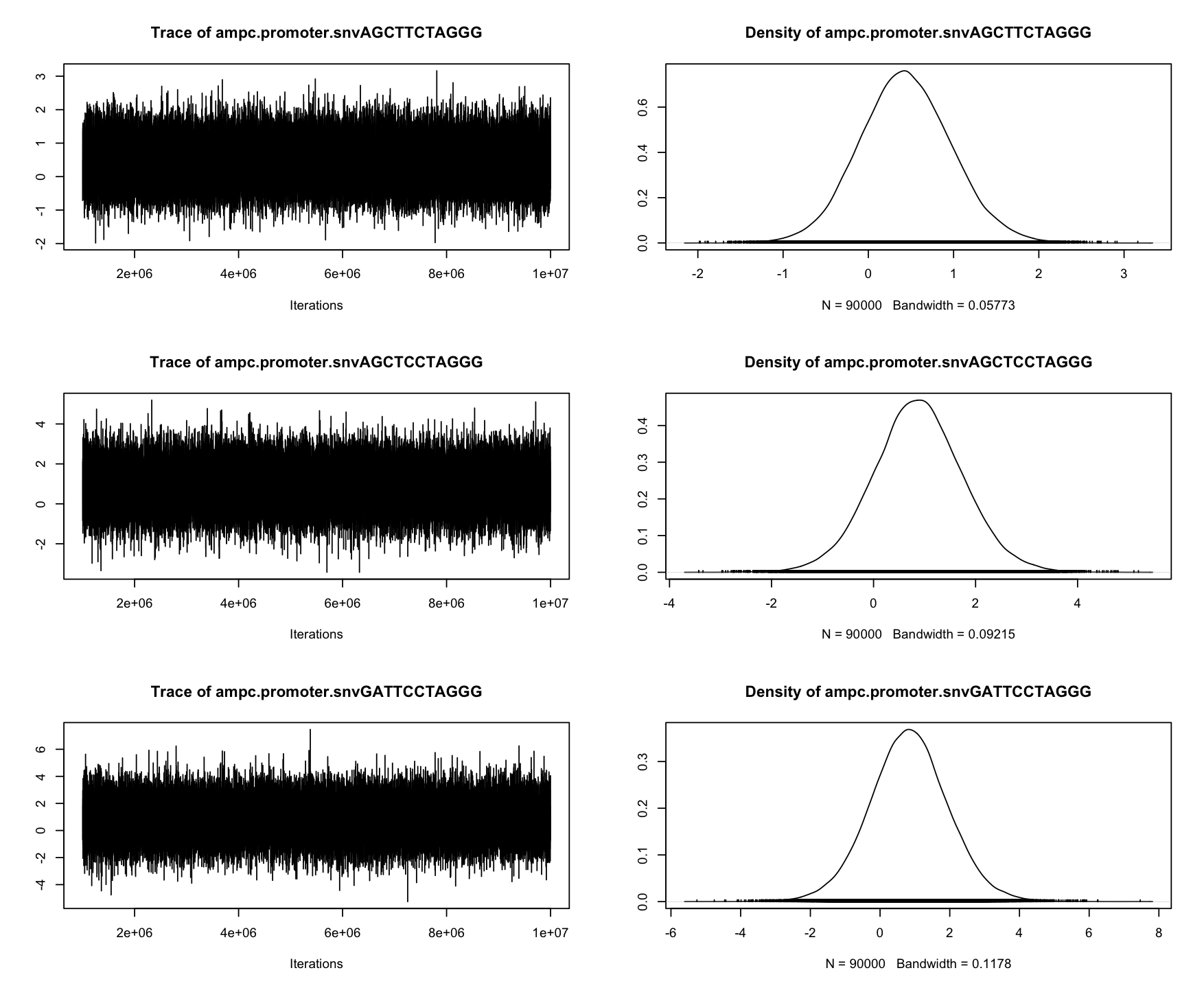

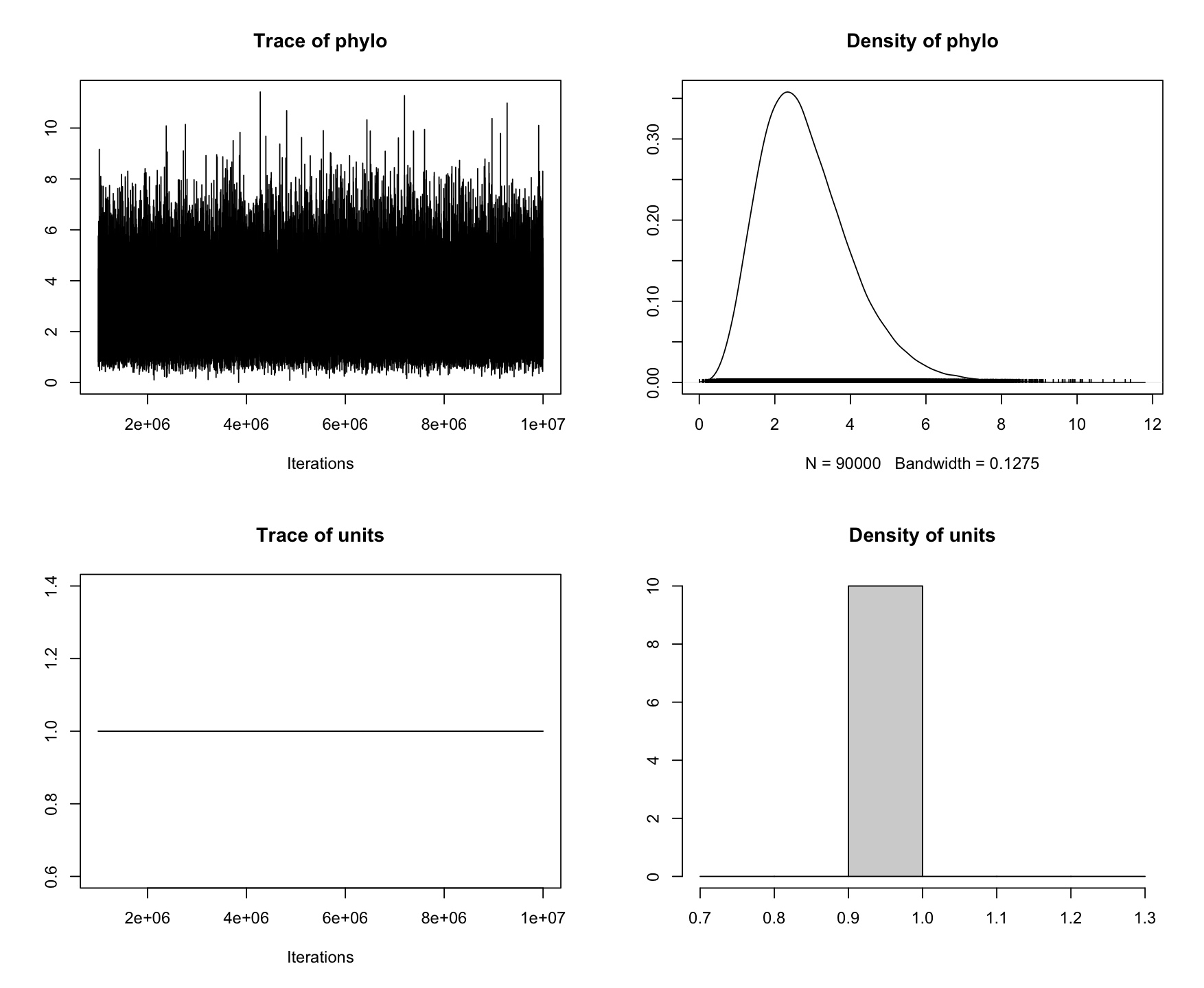

```
> mclist <- mcmc.list(chain.1$Sol, chain.2$Sol)
> gelman.diag(mclist)
Potential scale reduction factors:

                                Point est. Upper C.I.
(Intercept)                              1          1
tem1.isolate.copy.number.scaled          1          1
tem1.isolate.scaledTRUE                  1          1
ampc.promoter.snvAGCTTCTAGGG             1          1
ampc.promoter.snvAGCTCCTAGGG             1          1
ampc.promoter.snvGATTCCTAGGG             1          1
promoter.snvCGGCGA                       1          1
promoter.snvTGGCGA                       1          1
promoter.snvTGGCGG                       1          1
acrfTRUE                                 1          1
phylo.542                                1          1
phylo.416                                1          1
phylo.56                                 1          1
phylo.213                                1          1
phylo.54                                 1          1
phylo.435                                1          1
phylo.90                                 1          1
phylo.393                                1          1
phylo.72                                 1          1
phylo.217                                1          1
phylo.489                                1          1
phylo.189                                1          1
phylo.434                                1          1
phylo.194                                1          1
phylo.371                                1          1
phylo.127                                1          1
phylo.99                                 1          1
phylo.129                                1          1
phylo.8                                  1          1
phylo.414                                1          1
phylo.454                                1          1
phylo.259                                1          1
phylo.120                                1          1
phylo.145                                1          1
phylo.426                                1          1
phylo.46                                 1          1
phylo.49                                 1          1
phylo.394                                1          1
phylo.183                                1          1
phylo.30                                 1          1
phylo.151                                1          1
phylo.148                                1          1
phylo.236                                1          1
phylo.220                                1          1
phylo.117                                1          1
phylo.272                                1          1
phylo.400                                1          1
phylo.328                                1          1
phylo.415                                1          1
phylo.398                                1          1
phylo.252                                1          1
phylo.239                                1          1
phylo.246                                1          1
phylo.98                                 1          1
phylo.424                                1          1
phylo.66                                 1          1
phylo.108                                1          1
phylo.476                                1          1
phylo.468                                1          1
phylo.363                                1          1
phylo.21                                 1          1
phylo.128                                1          1
phylo.257                                1          1
phylo.242                                1          1
phylo.339                                1          1
phylo.455                                1          1
phylo.149                                1          1
phylo.417                                1          1
phylo.332                                1          1
phylo.373                                1          1
phylo.402                                1          1
phylo.52                                 1          1
phylo.498                                1          1
phylo.494                                1          1
phylo.472                                1          1
phylo.302                                1          1
phylo.445                                1          1
phylo.389                                1          1
phylo.61                                 1          1
phylo.299                                1          1
phylo.420                                1          1
phylo.103                                1          1
phylo.464                                1          1
phylo.449                                1          1
phylo.97                                 1          1
phylo.208                                1          1
phylo.329                                1          1
phylo.316                                1          1
phylo.80                                 1          1
phylo.438                                1          1
phylo.216                                1          1
phylo.324                                1          1
phylo.224                                1          1
phylo.447                                1          1
phylo.419                                1          1
phylo.175                                1          1
phylo.266                                1          1
phylo.343                                1          1
phylo.136                                1          1
phylo.156                                1          1
phylo.144                                1          1
phylo.404                                1          1
phylo.202                                1          1
phylo.439                                1          1
phylo.366                                1          1
phylo.285                                1          1
phylo.297                                1          1
phylo.133                                1          1
phylo.284                                1          1
phylo.512                                1          1
phylo.155                                1          1
phylo.253                                1          1
phylo.37                                 1          1
phylo.317                                1          1
phylo.143                                1          1
phylo.475                                1          1
phylo.342                                1          1
phylo.471                                1          1
phylo.264                                1          1
phylo.517                                1          1
phylo.507                                1          1
phylo.132                                1          1
phylo.39                                 1          1
phylo.286                                1          1
phylo.170                                1          1
phylo.76                                 1          1
phylo.500                                1          1
phylo.524                                1          1
phylo.533                                1          1
phylo.333                                1          1
phylo.410                                1          1
phylo.446                                1          1
phylo.344                                1          1
phylo.406                                1          1
phylo.433                                1          1
phylo.22                                 1          1
phylo.430                                1          1
phylo.288                                1          1
phylo.270                                1          1
phylo.511                                1          1
phylo.429                                1          1
phylo.277                                1          1
phylo.322                                1          1
phylo.482                                1          1
phylo.262                                1          1
phylo.387                                1          1
phylo.58                                 1          1
phylo.193                                1          1
phylo.388                                1          1
phylo.158                                1          1
phylo.12                                 1          1
phylo.110                                1          1
phylo.276                                1          1
phylo.412                                1          1
phylo.411                                1          1
phylo.365                                1          1
phylo.82                                 1          1
phylo.92                                 1          1
phylo.492                                1          1
phylo.370                                1          1
phylo.137                                1          1
phylo.483                                1          1
phylo.267                                1          1
phylo.421                                1          1
phylo.237                                1          1
phylo.200                                1          1
phylo.536                                1          1
phylo.260                                1          1
phylo.537                                1          1
phylo.477                                1          1
phylo.518                                1          1
phylo.522                                1          1
phylo.55                                 1          1
phylo.358                                1          1
phylo.197                                1          1
phylo.493                                1          1
phylo.41                                 1          1
phylo.40                                 1          1
phylo.34                                 1          1
phylo.94                                 1          1
phylo.408                                1          1
phylo.292                                1          1
phylo.248                                1          1
phylo.109                                1          1
phylo.1                                  1          1
phylo.214                                1          1
phylo.480                                1          1
phylo.423                                1          1
phylo.247                                1          1
phylo.268                                1          1
phylo.157                                1          1
phylo.385                                1          1
phylo.353                                1          1
phylo.53                                 1          1
phylo.211                                1          1
phylo.265                                1          1
phylo.139                                1          1
phylo.198                                1          1
phylo.396                                1          1
phylo.456                                1          1
phylo.245                                1          1
phylo.263                                1          1
phylo.134                                1          1
phylo.3                                  1          1
phylo.95                                 1          1
phylo.514                                1          1
phylo.525                                1          1
phylo.529                                1          1
phylo.162                                1          1
phylo.515                                1          1
phylo.282                                1          1
phylo.427                                1          1
phylo.240                                1          1
phylo.166                                1          1
phylo.60                                 1          1
phylo.182                                1          1
phylo.532                                1          1
phylo.273                                1          1
phylo.119                                1          1
phylo.11                                 1          1
phylo.355                                1          1
phylo.241                                1          1
phylo.315                                1          1
phylo.24                                 1          1
phylo.203                                1          1
phylo.121                                1          1
phylo.305                                1          1
phylo.534                                1          1
phylo.167                                1          1
phylo.336                                1          1
phylo.16                                 1          1
phylo.330                                1          1
phylo.320                                1          1
phylo.188                                1          1
phylo.274                                1          1
phylo.280                                1          1
phylo.281                                1          1
phylo.89                                 1          1
phylo.531                                1          1
phylo.106                                1          1
phylo.227                                1          1
phylo.164                                1          1
phylo.382                                1          1
phylo.192                                1          1
phylo.301                                1          1
phylo.506                                1          1
phylo.334                                1          1
phylo.177                                1          1
phylo.469                                1          1
phylo.2                                  1          1
phylo.470                                1          1
phylo.351                                1          1
phylo.31                                 1          1
phylo.159                                1          1
phylo.187                                1          1
phylo.488                                1          1
phylo.486                                1          1
phylo.225                                1          1
phylo.32                                 1          1
phylo.541                                1          1
phylo.380                                1          1
phylo.462                                1          1
phylo.451                                1          1
phylo.403                                1          1
phylo.115                                1          1
phylo.250                                1          1
phylo.93                                 1          1
phylo.230                                1          1
phylo.457                                1          1
phylo.313                                1          1
phylo.331                                1          1
phylo.176                                1          1
phylo.75                                 1          1
phylo.296                                1          1
phylo.205                                1          1
phylo.256                                1          1
phylo.88                                 1          1
phylo.487                                1          1
phylo.347                                1          1
phylo.442                                1          1
phylo.495                                1          1
phylo.116                                1          1
phylo.233                                1          1
phylo.539                                1          1
phylo.83                                 1          1
phylo.444                                1          1
phylo.508                                1          1
phylo.33                                 1          1
phylo.57                                 1          1
phylo.142                                1          1
phylo.311                                1          1
phylo.386                                1          1
phylo.135                                1          1
phylo.19                                 1          1
phylo.59                                 1          1
phylo.287                                1          1
phylo.375                                1          1
phylo.490                                1          1
phylo.448                                1          1
phylo.10                                 1          1
phylo.399                                1          1
phylo.466                                1          1
phylo.310                                1          1
phylo.543                                1          1
phylo.504                                1          1
phylo.174                                1          1
phylo.499                                1          1
phylo.126                                1          1
phylo.413                                1          1
phylo.125                                1          1
phylo.201                                1          1
phylo.513                                1          1
phylo.69                                 1          1
phylo.340                                1          1
phylo.179                                1          1
phylo.25                                 1          1
phylo.436                                1          1
phylo.546                                1          1
phylo.279                                1          1
phylo.485                                1          1
phylo.38                                 1          1
phylo.422                                1          1
phylo.229                                1          1
phylo.35                                 1          1
phylo.131                                1          1
phylo.71                                 1          1
phylo.6                                  1          1
phylo.28                                 1          1
phylo.401                                1          1
phylo.222                                1          1
phylo.243                                1          1
phylo.42                                 1          1
phylo.63                                 1          1
phylo.18                                 1          1
phylo.123                                1          1
phylo.140                                1          1
phylo.502                                1          1
phylo.503                                1          1
phylo.341                                1          1
phylo.238                                1          1
phylo.478                                1          1
phylo.443                                1          1
phylo.44                                 1          1
phylo.544                                1          1
phylo.204                                1          1
phylo.510                                1          1
phylo.314                                1          1
phylo.124                                1          1
phylo.64                                 1          1
phylo.219                                1          1
phylo.178                                1          1
phylo.345                                1          1
phylo.479                                1          1
phylo.307                                1          1
phylo.275                                1          1
phylo.540                                1          1
phylo.23                                 1          1
phylo.73                                 1          1
phylo.530                                1          1
phylo.244                                1          1
phylo.191                                1          1
phylo.474                                1          1
phylo.460                                1          1
phylo.528                                1          1
phylo.295                                1          1
phylo.459                                1          1
phylo.463                                1          1
phylo.79                                 1          1
phylo.481                                1          1
phylo.190                                1          1
phylo.461                                1          1
phylo.535                                1          1
phylo.223                                1          1
phylo.29                                 1          1
phylo.65                                 1          1
phylo.283                                1          1
phylo.425                                1          1
phylo.255                                1          1
phylo.521                                1          1
phylo.199                                1          1
phylo.107                                1          1
phylo.294                                1          1
phylo.122                                1          1
phylo.87                                 1          1
phylo.105                                1          1
phylo.138                                1          1
phylo.440                                1          1

Multivariate psrf

1
```

### **Combined phenotype model specification**

The combined model investigates how *bla*_TEM-1_ expression affects MIC, accounting for phylogenetic effects and scaling these effects by a parameter *θ*. The random effects structure for expression for isolate $i$ is given by

$$\text{exp}_{i}= \text{p}_{i} + \text{a}_{i}$$

where $\text{p}_{i}$ is the phylogenetic effect and $\text{a}_{i}$ is the isolate main effect. The random effects structure for MIC for isolate $i$ is given by

$$\text{MIC}_{i}= {\theta(\text{p}}_{i} + \text{a}_{i}) + \text{u}_{i}$$

where $\theta$ is the scaling factor and $\text{u}_{i}$ phylogenetic effect specifically fitted for MIC.

The covariance structure can be represented as a series of regressions based on the random effects. The variance of phylogenetic effect on expression is given by

$$\text{Var}(p)= \sigma_{p}^{2}$$

the covariance between expression and MIC is given by

$$\text{Cov}(exp, MIC)=\theta\times\sigma_{p}^{2}$$

and the variance of MIC is given by

$$\text{Var}(MIC)=\text{Var}(u) +\theta^{2}\times\sigma_{p}^{2}$$

This approach demonstrates that the MIC phenotype is influenced by the expression levels, with the relationship modulated by the scaling parameter *θ*. The model assumes a causal relationship where expression impacts MIC, and this impact is consistent across all random effects. The combined model with fixed and random effects is then given by

$$Y_{i}=\left\{ \begin{aligned} X_{i}^{exp}\boldsymbol{\beta}+\text{exp}_{i} + \varepsilon_{i}^{exp}\text{ }\text{ }\text{for}\text{ expression} \\ X_{i}^{MIC}\boldsymbol{\beta}+\text{MIC}_{i} + \varepsilon_{i}^{MIC}\text{ }\text{for}\text{ MIC} \end{aligned} \right.$$

Note, that due to model complexity, we used ordinal encoding for MIC categories.

### **Combined phenotype model outputs**

```
> summary(chain.1)

 Iterations = 1000001:9999901
 Thinning interval  = 100
 Sample size  = 90000

 DIC: NaN

 G-structure:  ~phylo

      post.mean  l-95% CI u-95% CI eff.samp
phylo   0.09938 1.489e-08   0.2427    57787

               ~isolate.id

           post.mean l-95% CI u-95% CI eff.samp
isolate.id    0.1818   0.1039   0.2705    50740

               ~us(at.level(trait, "mic")):phylo

                                                    post.mean l-95% CI u-95% CI eff.samp
at.level(trait, "mic"):at.level(trait, "mic").phylo    0.5906   0.1418      1.1    83991

 R-structure:  ~idh(trait):units

               post.mean l-95% CI u-95% CI eff.samp
traitexp.units   0.04027  0.03245  0.04857    15534
traitmic.units   0.41560  0.23881  0.60102    57755

 Location effects: y ~ -1 + trait:(1 + tem1.isolate.scaled + tem1.isolate.copy.number.scaled + promoter.snv)

                                         post.mean l-95% CI u-95% CI eff.samp  pMCMC
(Intercept)                                4.32938  3.87446  4.78219    90000 <1e-05 ***
traitexp:tem1.isolate.scaledFALSE         -4.54617 -5.19684 -3.92338    90000 <1e-05 ***
traitmic:tem1.isolate.scaledFALSE         -0.46167 -0.91183 -0.00569    90000 0.0465 *
traitexp:tem1.isolate.copy.number.scaled  -1.02103 -1.96612 -0.09322    88080 0.0349 *
traitmic:tem1.isolate.copy.number.scaled   1.03191  0.72891  1.34028    88658 <1e-05 ***
traitexp:promoter.snvCGGCGA               -0.24776 -0.50319  0.01327    86917 0.0623 .
traitmic:promoter.snvCGGCGA                0.03181 -0.23004  0.28456    90000 0.8064
traitexp:promoter.snvTGGCGA               -1.66270 -2.01689 -1.30631    87829 <1e-05 ***
traitmic:promoter.snvTGGCGA                2.35052  1.83735  2.88213    90000 <1e-05 ***
traitexp:promoter.snvTGGCGG               -1.68334 -2.06755 -1.30517    87857 <1e-05 ***
traitmic:promoter.snvTGGCGG                2.23015  1.69807  2.77481    90000 <1e-05 ***
---
Signif. codes:  0 ‘***’ 0.001 ‘**’ 0.01 ‘*’ 0.05 ‘.’ 0.1 ‘ ’ 1

 Theta scale parameter:

            post.mean l-95% CI u-95% CI eff.samp   pMCMC
theta_scale   -1.1267  -1.4853  -0.7196     8506 0.00231 **
---
Signif. codes:  0 ‘***’ 0.001 ‘**’ 0.01 ‘*’ 0.05 ‘.’ 0.1 ‘ ’ 1

> autocorr.diag(chain.1$VCV)
               phylo   isolate.id at.level(trait, "mic"):at.level(trait, "mic").phylo traitexp.units traitmic.units
Lag 0    1.000000000  1.000000000                                        1.000000e+00   1.0000000000   1.0000000000
Lag 100  0.112367643  0.072780947                                        1.409324e-02   0.2537680979   0.0829974214
Lag 500  0.015074121  0.022672690                                        6.601437e-03   0.1825237734   0.0196155673
Lag 1000 0.006205217  0.009637130                                       -1.172422e-03   0.1062380205   0.0085013712
Lag 5000 0.002281869 -0.001722614                                        5.754047e-05   0.0001475553  -0.0005311794

> plot(chain.1)
```

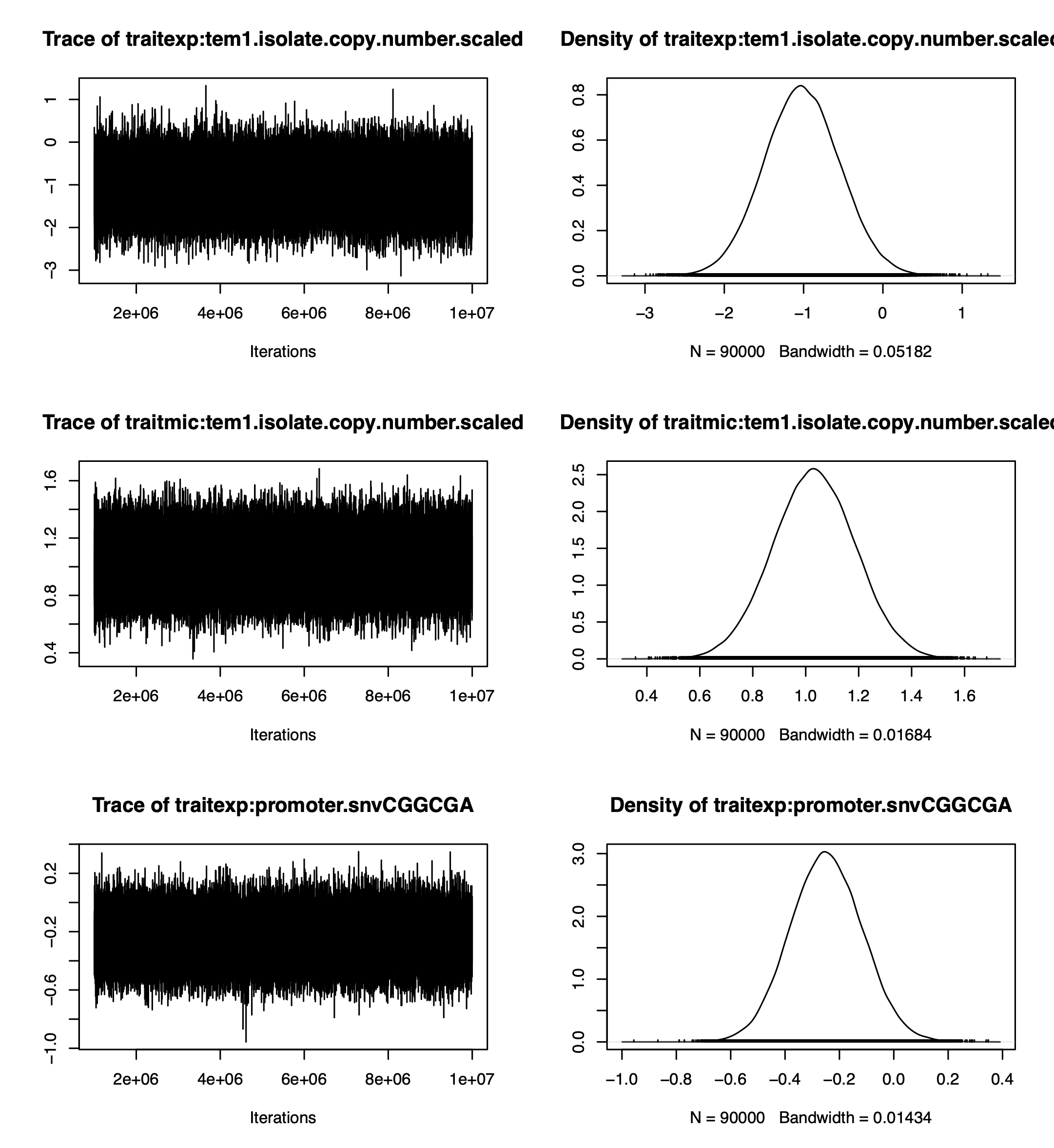

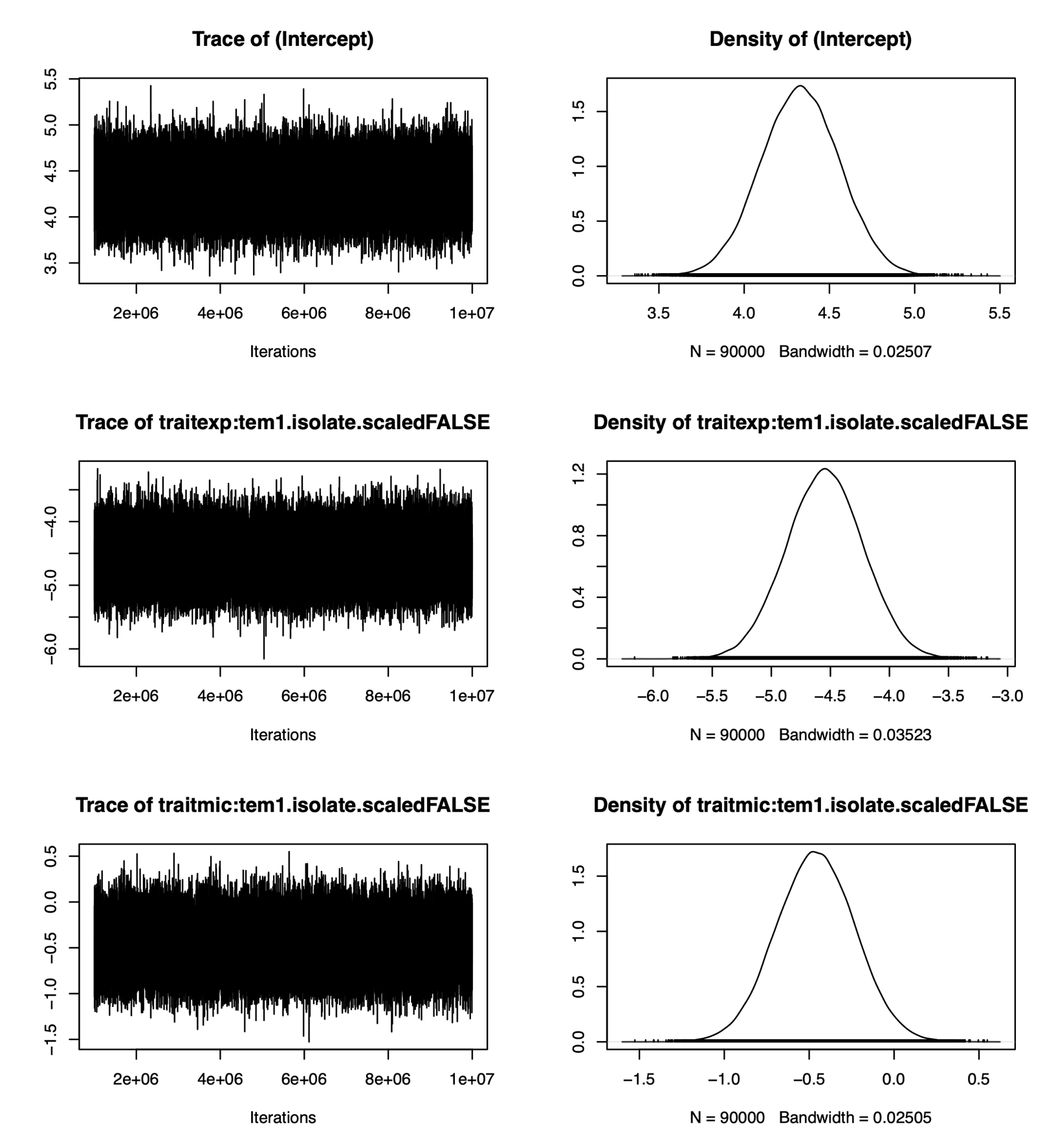

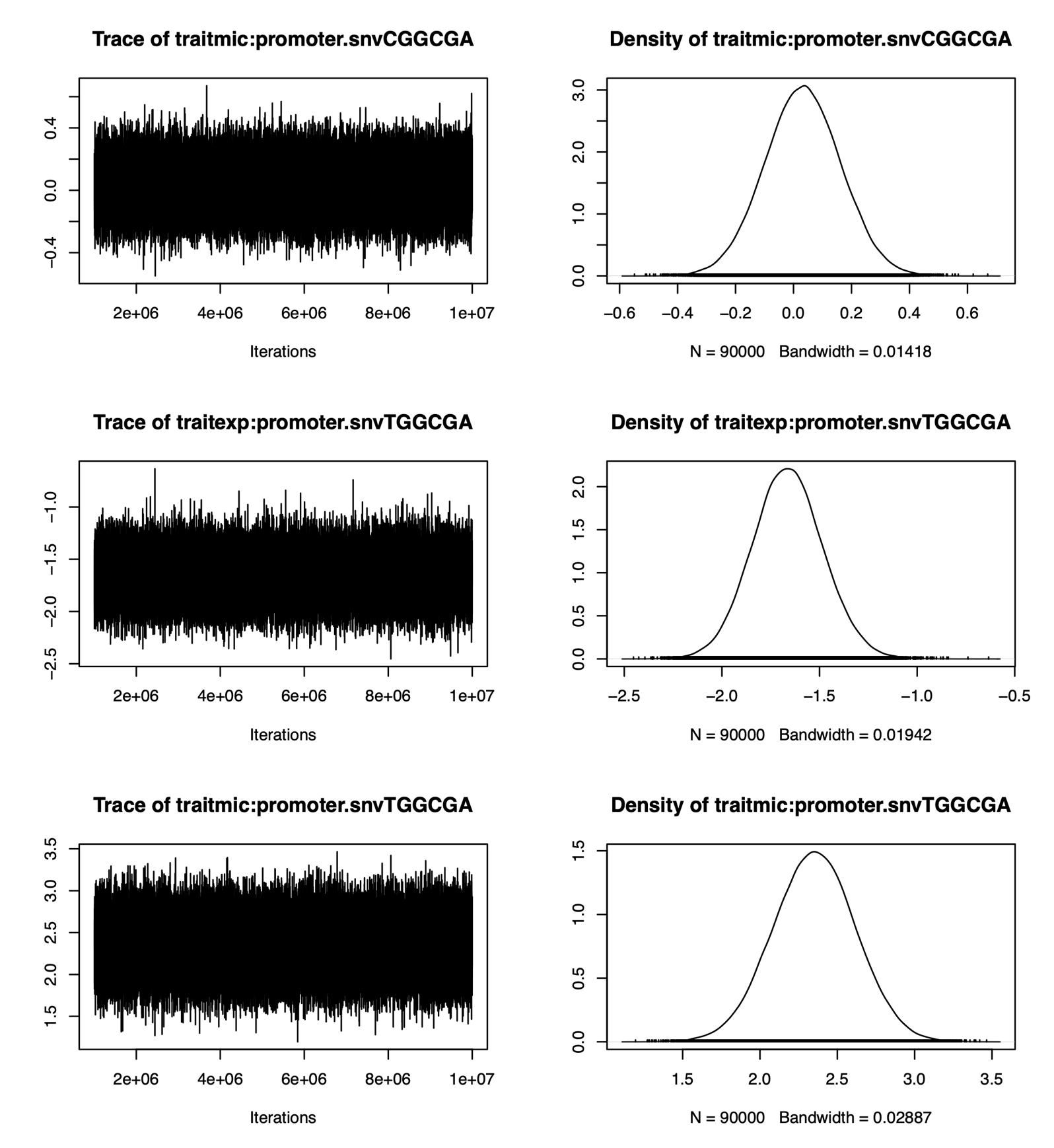

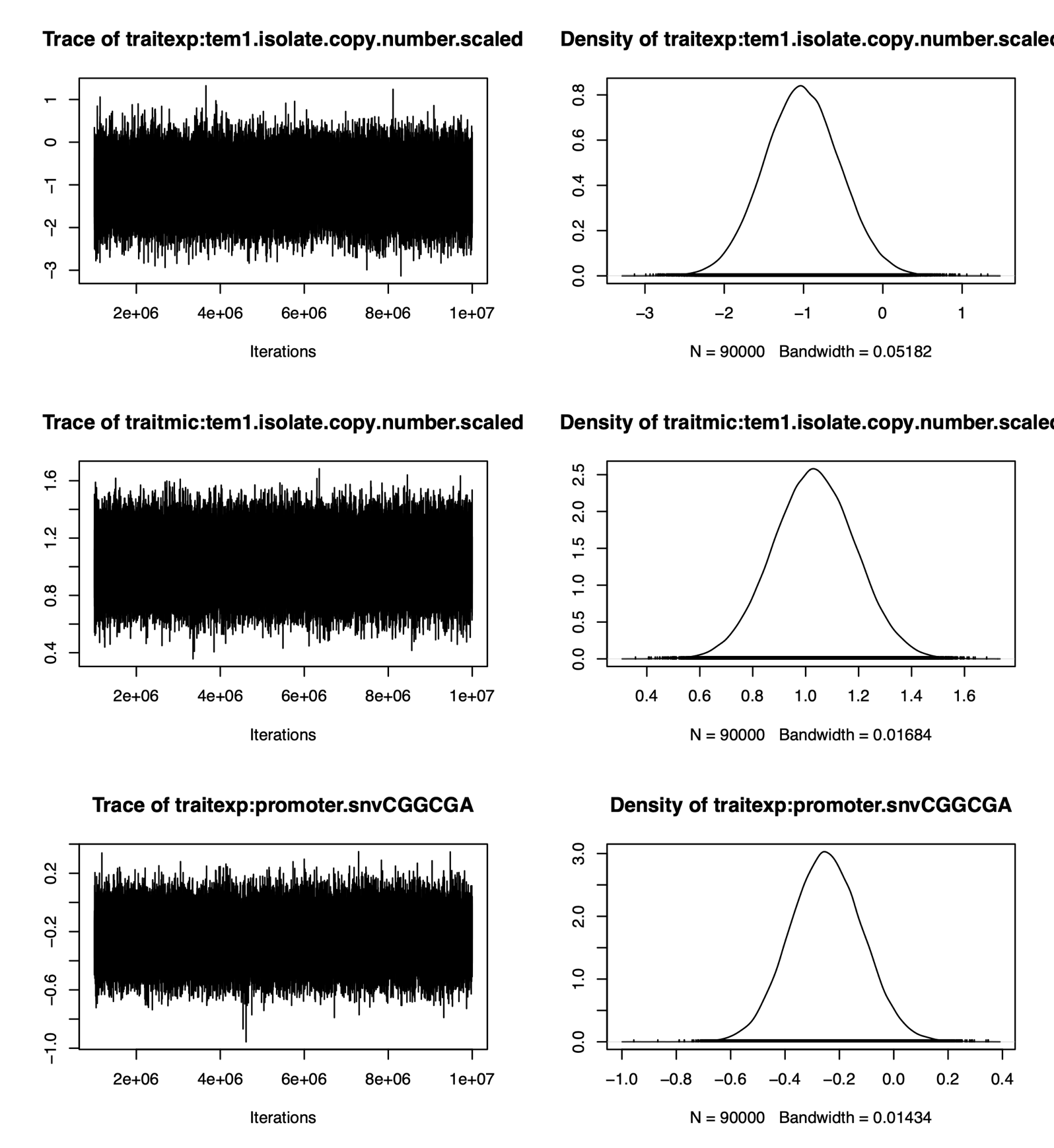

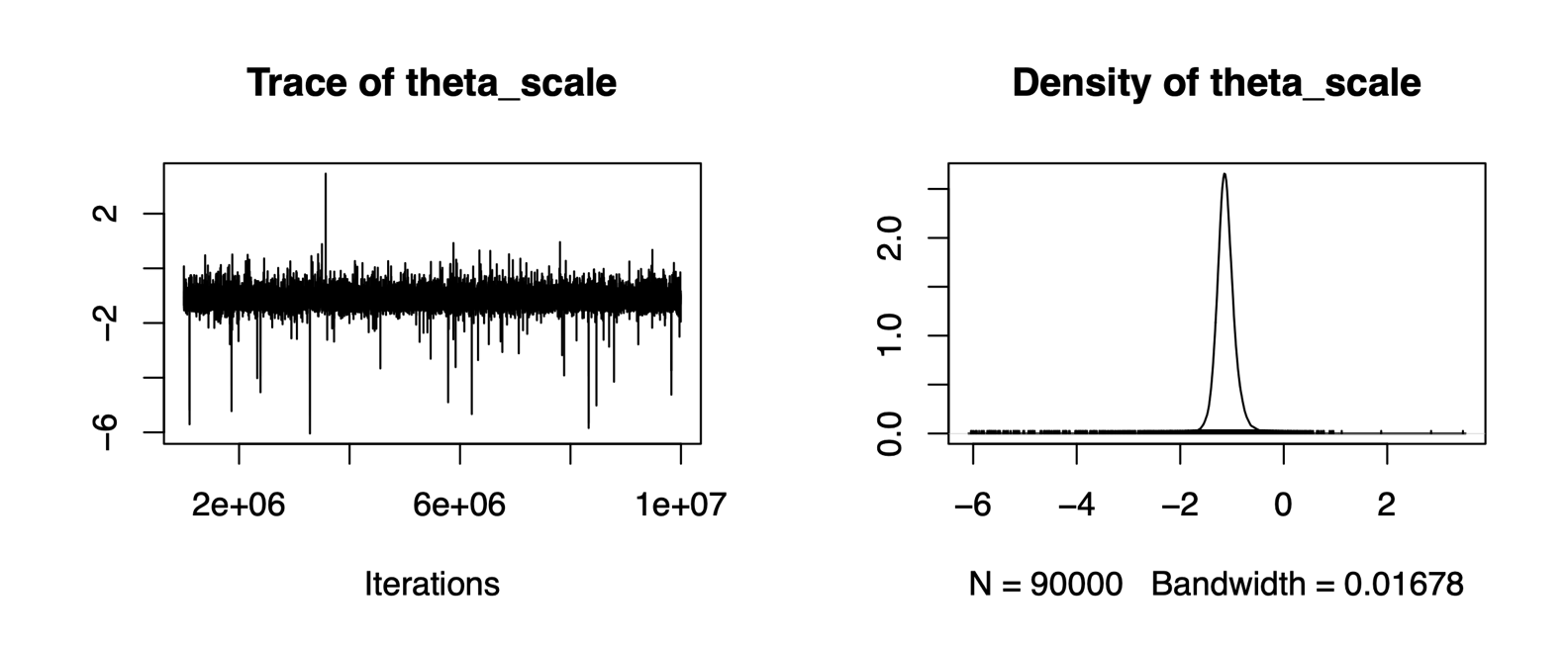

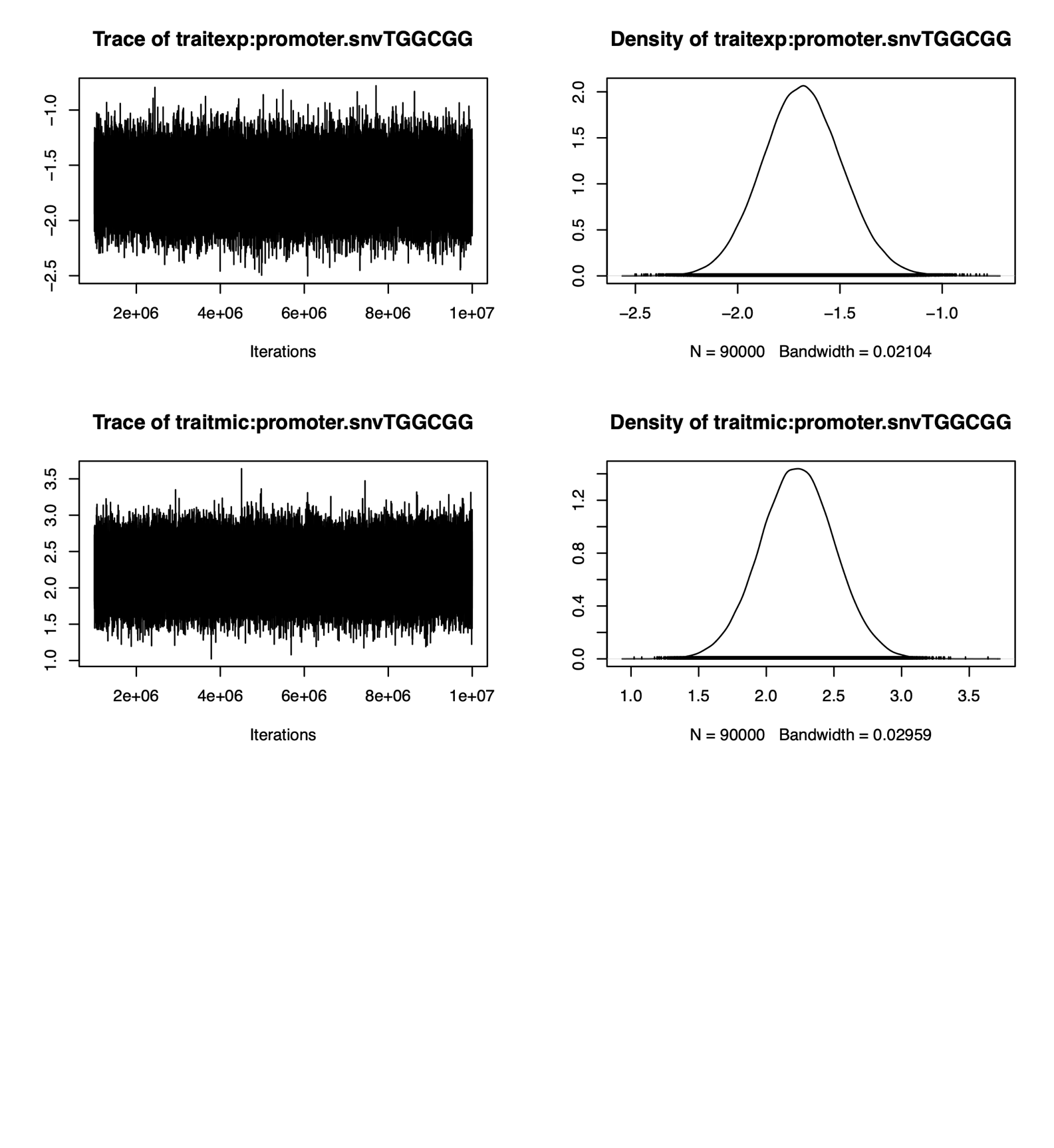

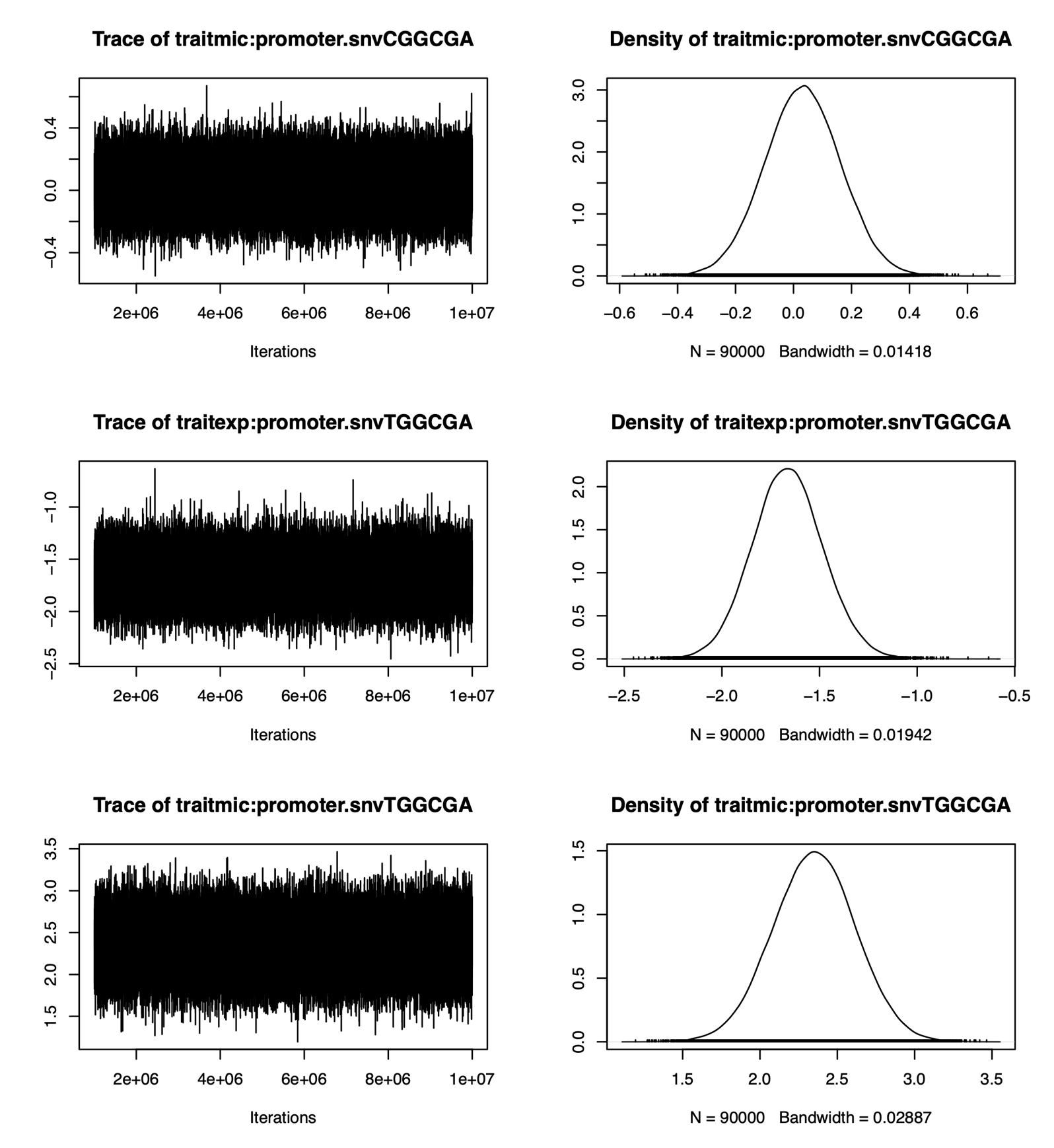

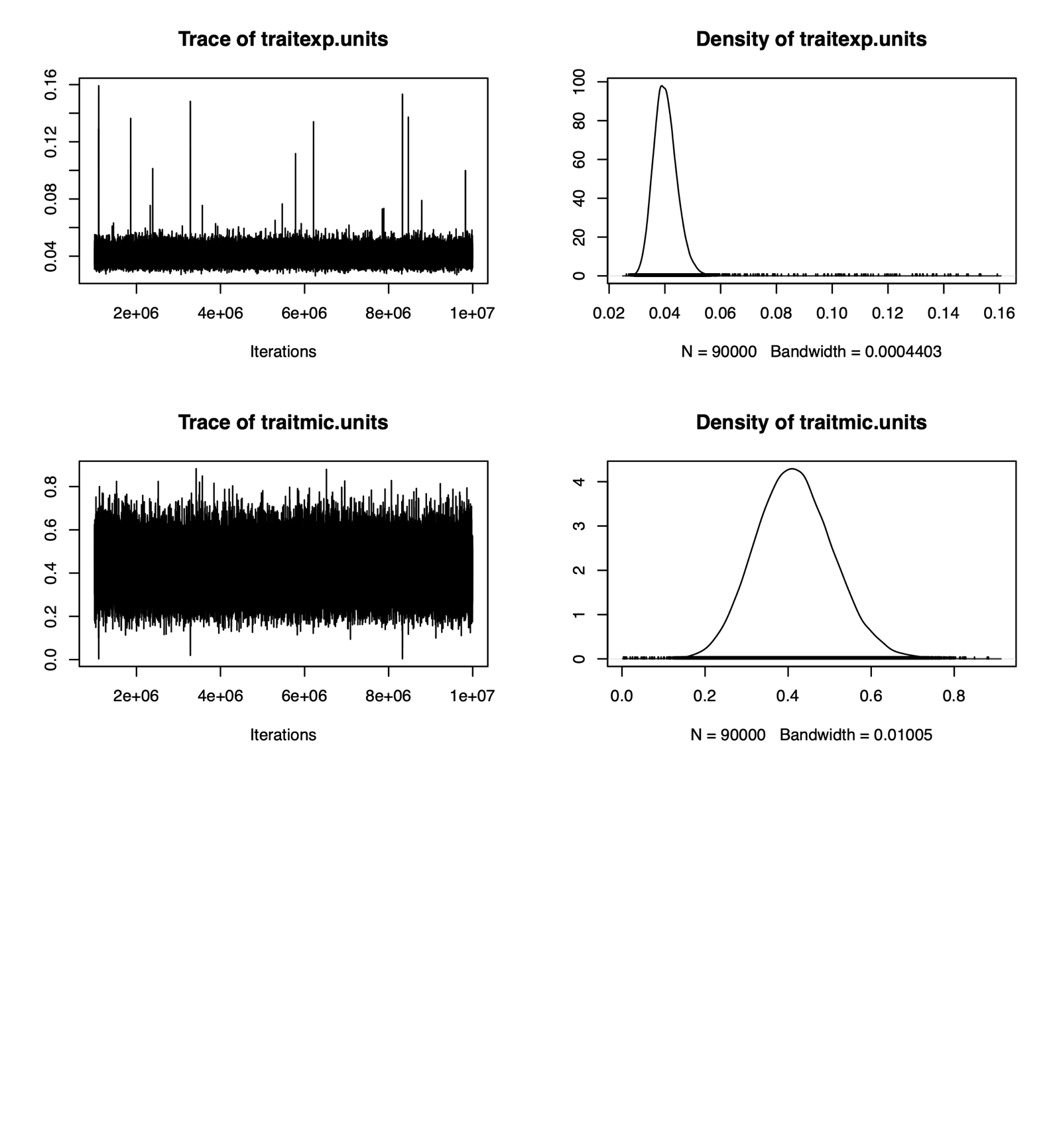

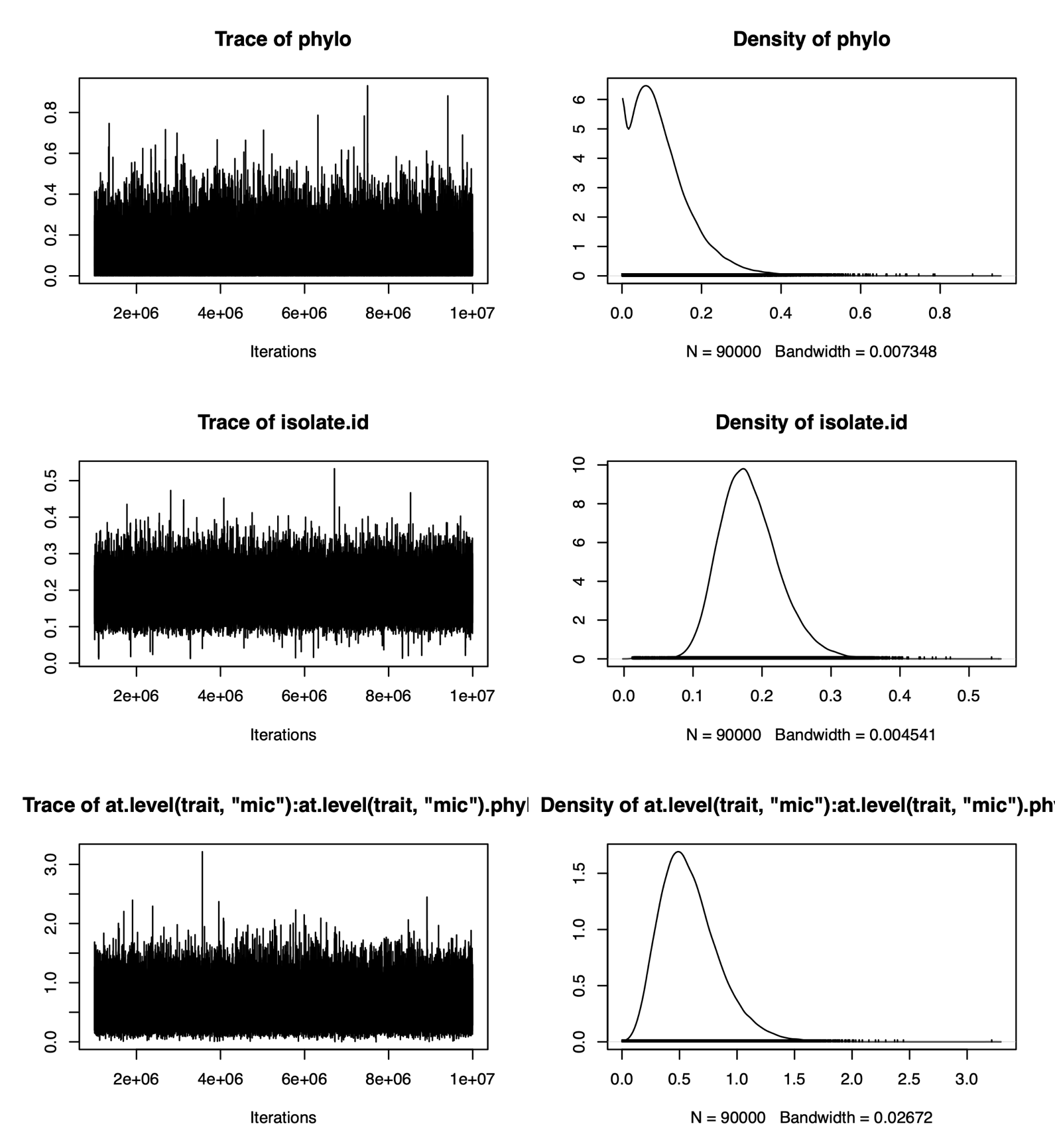

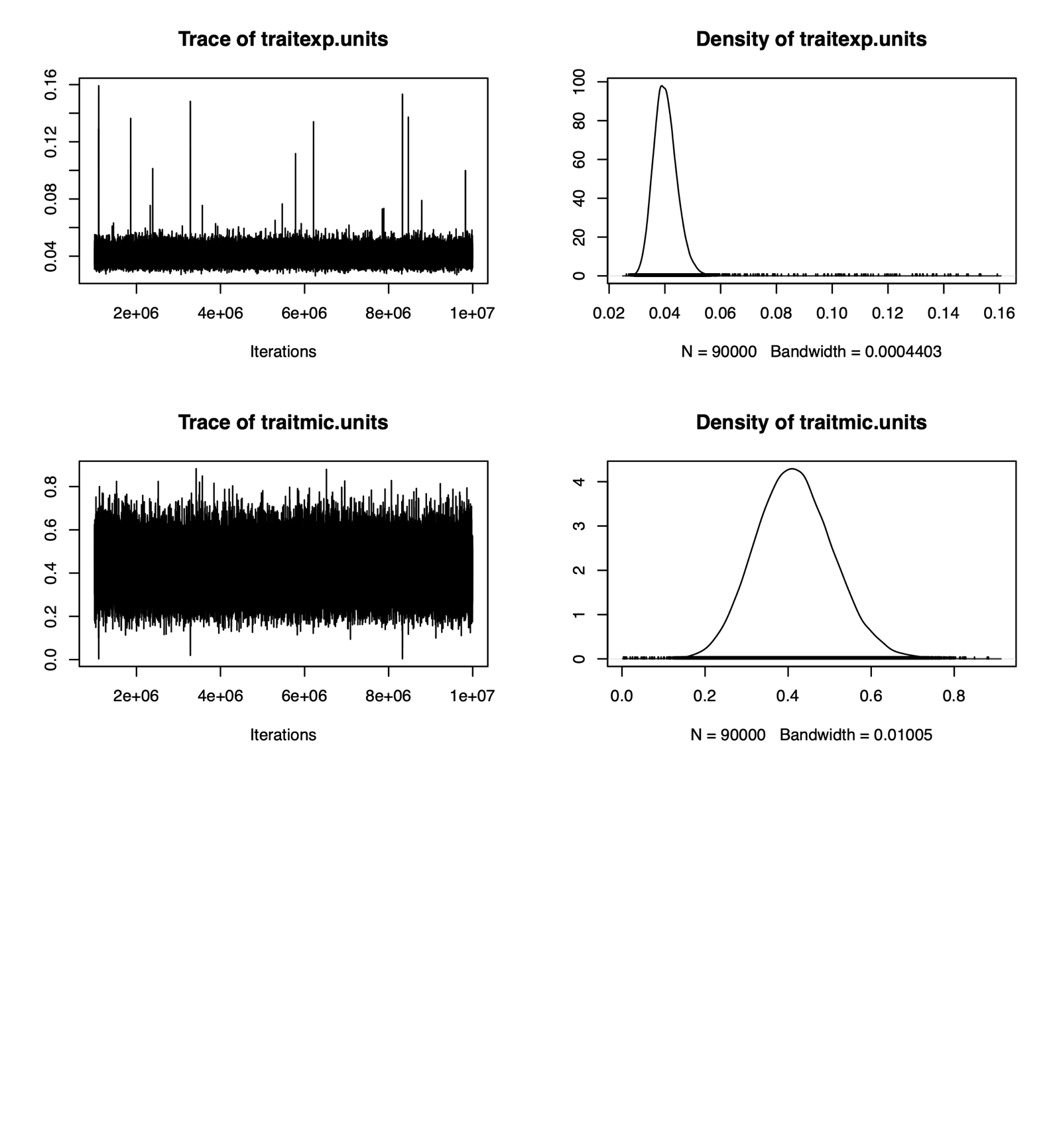

```
> summary(chain.2)

 Iterations = 1000001:9999901
 Thinning interval  = 100
 Sample size  = 90000

 DIC: NaN

 G-structure:  ~phylo

      post.mean  l-95% CI u-95% CI eff.samp
phylo    0.1003 6.355e-11   0.2435    61115

               ~isolate.id

           post.mean l-95% CI u-95% CI eff.samp
isolate.id    0.1818   0.1024   0.2703    48151

               ~us(at.level(trait, "mic")):phylo

                                                    post.mean l-95% CI u-95% CI eff.samp
at.level(trait, "mic"):at.level(trait, "mic").phylo    0.5897   0.1465    1.095    84462

 R-structure:  ~idh(trait):units

               post.mean l-95% CI u-95% CI eff.samp
traitexp.units   0.04026  0.03233   0.0484    13141
traitmic.units   0.41494  0.23108   0.5954    56352

 Location effects: y ~ -1 + trait:(1 + tem1.isolate.scaled + tem1.isolate.copy.number.scaled + promoter.snv)

                                         post.mean  l-95% CI  u-95% CI eff.samp  pMCMC
(Intercept)                               4.328525  3.874485  4.782199    90000 <1e-05 ***
traitexp:tem1.isolate.scaledFALSE        -4.544511 -5.171154 -3.902730    90000 <1e-05 ***
traitmic:tem1.isolate.scaledFALSE        -0.461352 -0.899921  0.002839    90000 0.0448 *
traitexp:tem1.isolate.copy.number.scaled -1.019478 -1.946873 -0.062113    90000 0.0352 *
traitmic:tem1.isolate.copy.number.scaled  1.033209  0.729652  1.336409    90092 <1e-05 ***
traitexp:promoter.snvCGGCGA              -0.248080 -0.507897  0.008901    90000 0.0627 .
traitmic:promoter.snvCGGCGA               0.032095 -0.223837  0.290353    90000 0.8044
traitexp:promoter.snvTGGCGA              -1.662544 -2.014827 -1.300562    83716 <1e-05 ***
traitmic:promoter.snvTGGCGA               2.351595  1.831656  2.880991    91212 <1e-05 ***
traitexp:promoter.snvTGGCGG              -1.683006 -2.056777 -1.290550    87655 <1e-05 ***
traitmic:promoter.snvTGGCGG               2.230071  1.706237  2.771809    90000 <1e-05 ***
---
Signif. codes:  0 ‘***’ 0.001 ‘**’ 0.01 ‘*’ 0.05 ‘.’ 0.1 ‘ ’ 1

 Theta scale parameter:

            post.mean l-95% CI u-95% CI eff.samp   pMCMC
theta_scale   -1.1272  -1.4934  -0.7211     8575 0.00198 **
---
Signif. codes:  0 ‘***’ 0.001 ‘**’ 0.01 ‘*’ 0.05 ‘.’ 0.1 ‘ ’ 1

> autocorr.diag(chain.2$VCV)
                phylo   isolate.id at.level(trait, "mic"):at.level(trait, "mic").phylo traitexp.units traitmic.units
Lag 0    1.0000000000  1.000000000                                         1.000000000    1.000000000    1.000000000
Lag 100  0.1147704162  0.076378596                                         0.008282837    0.307716514    0.086258143
Lag 500  0.0125540987  0.032054763                                        -0.002772776    0.191916176    0.023457348
Lag 1000 0.0003901405  0.015048908                                         0.007131786    0.120163028    0.014677994
Lag 5000 0.0022367820 -0.001287983                                         0.003862192   -0.001317448    0.001987818

> plot(chain.2)
```

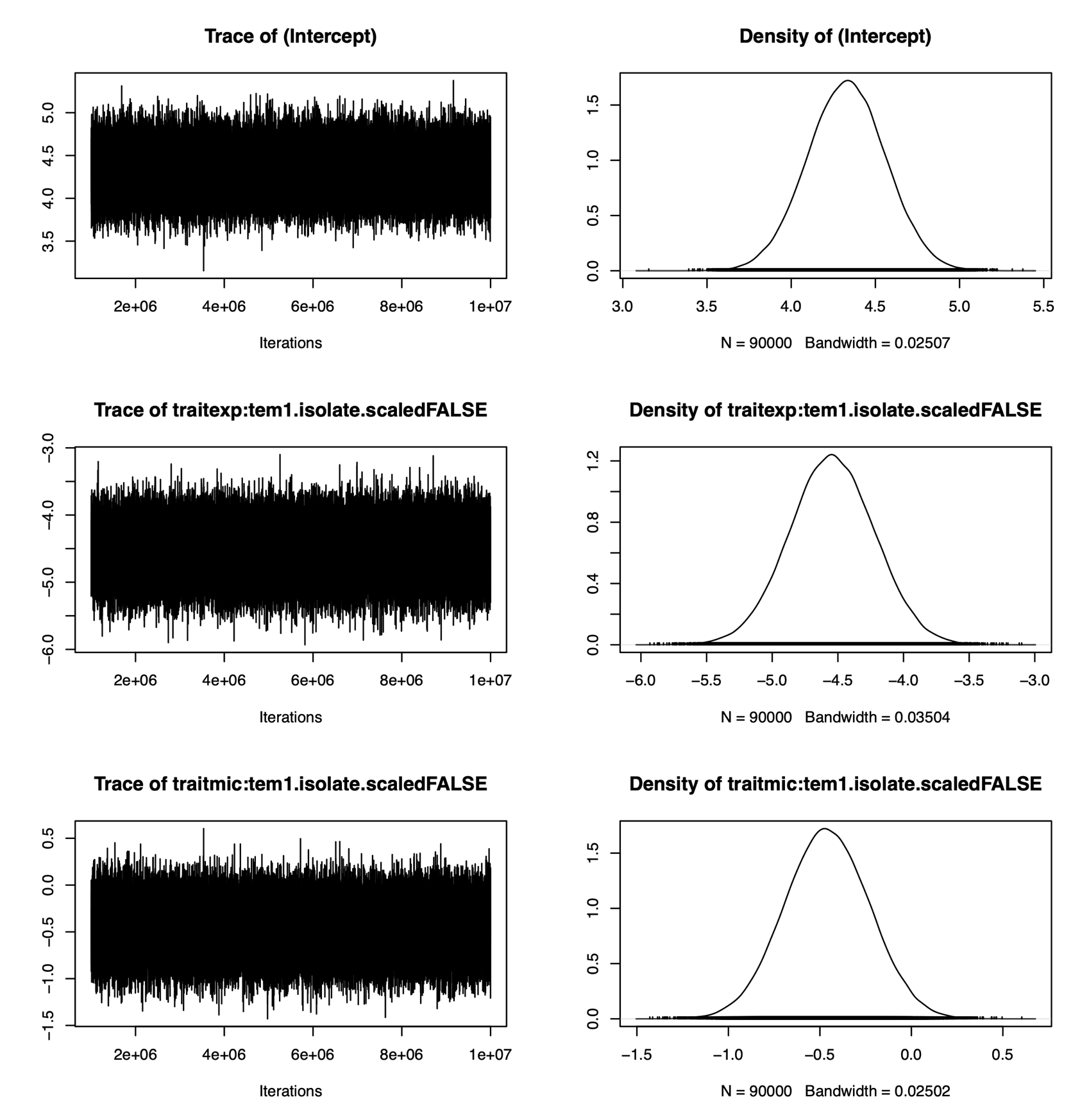

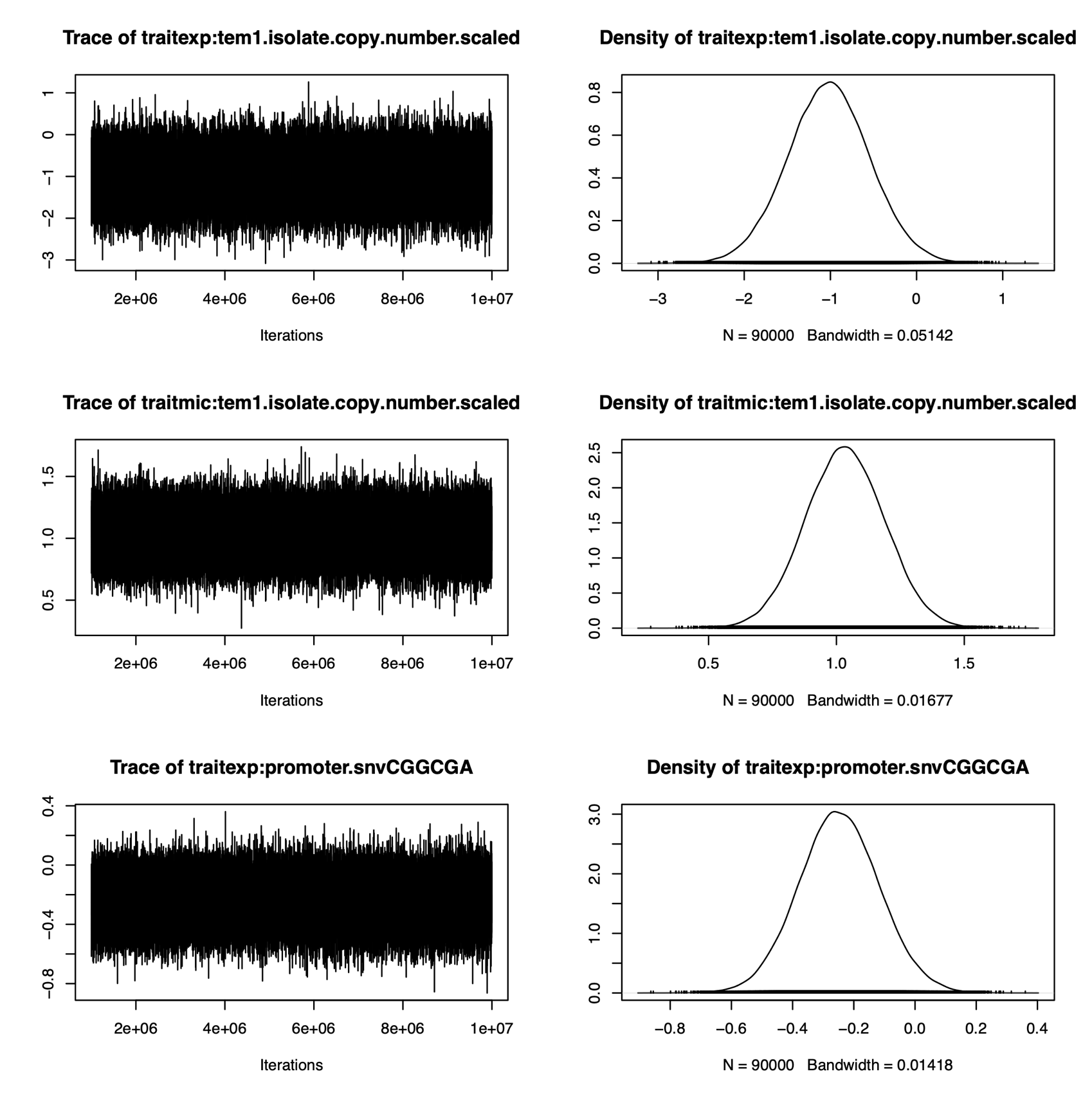

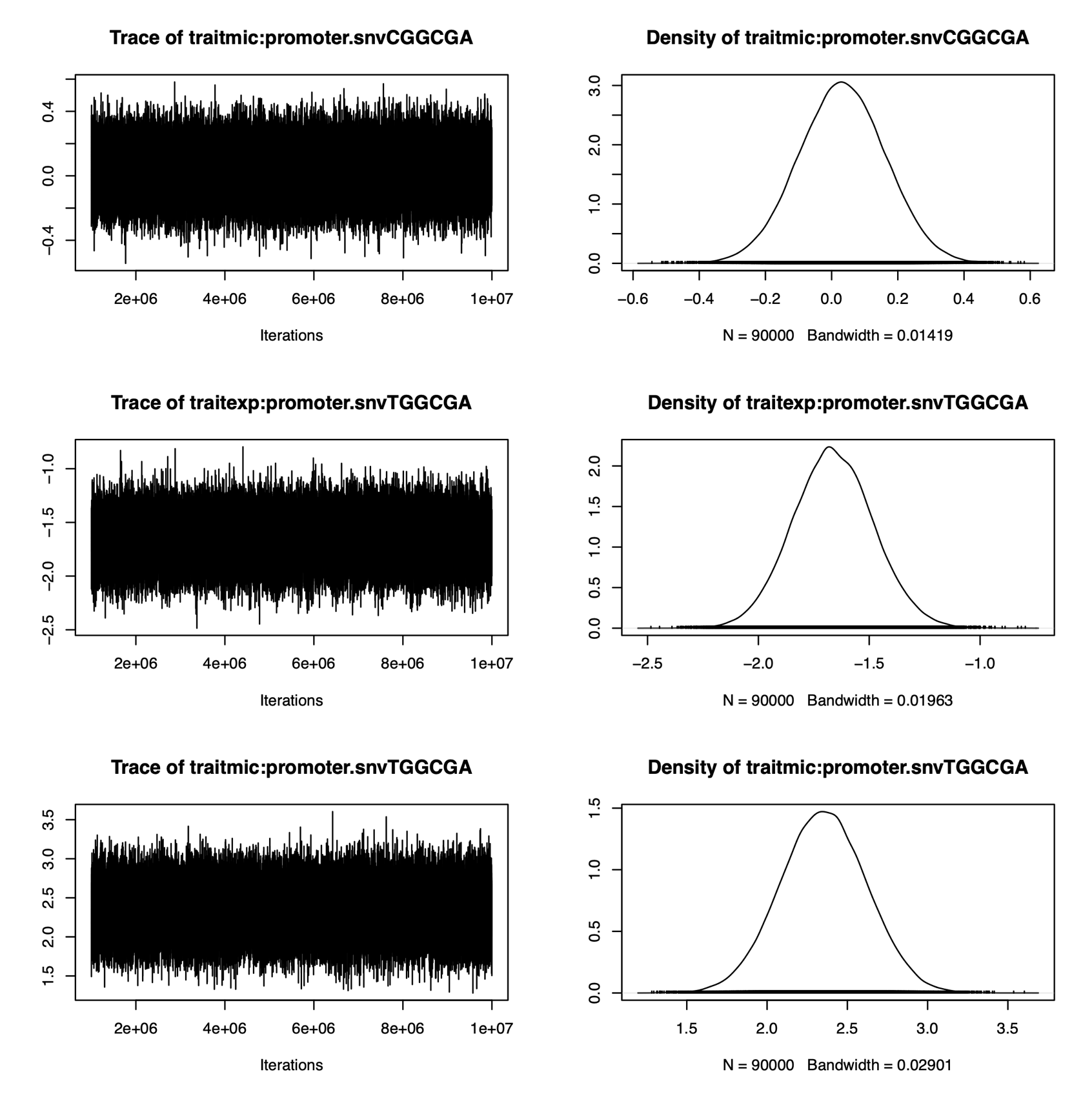

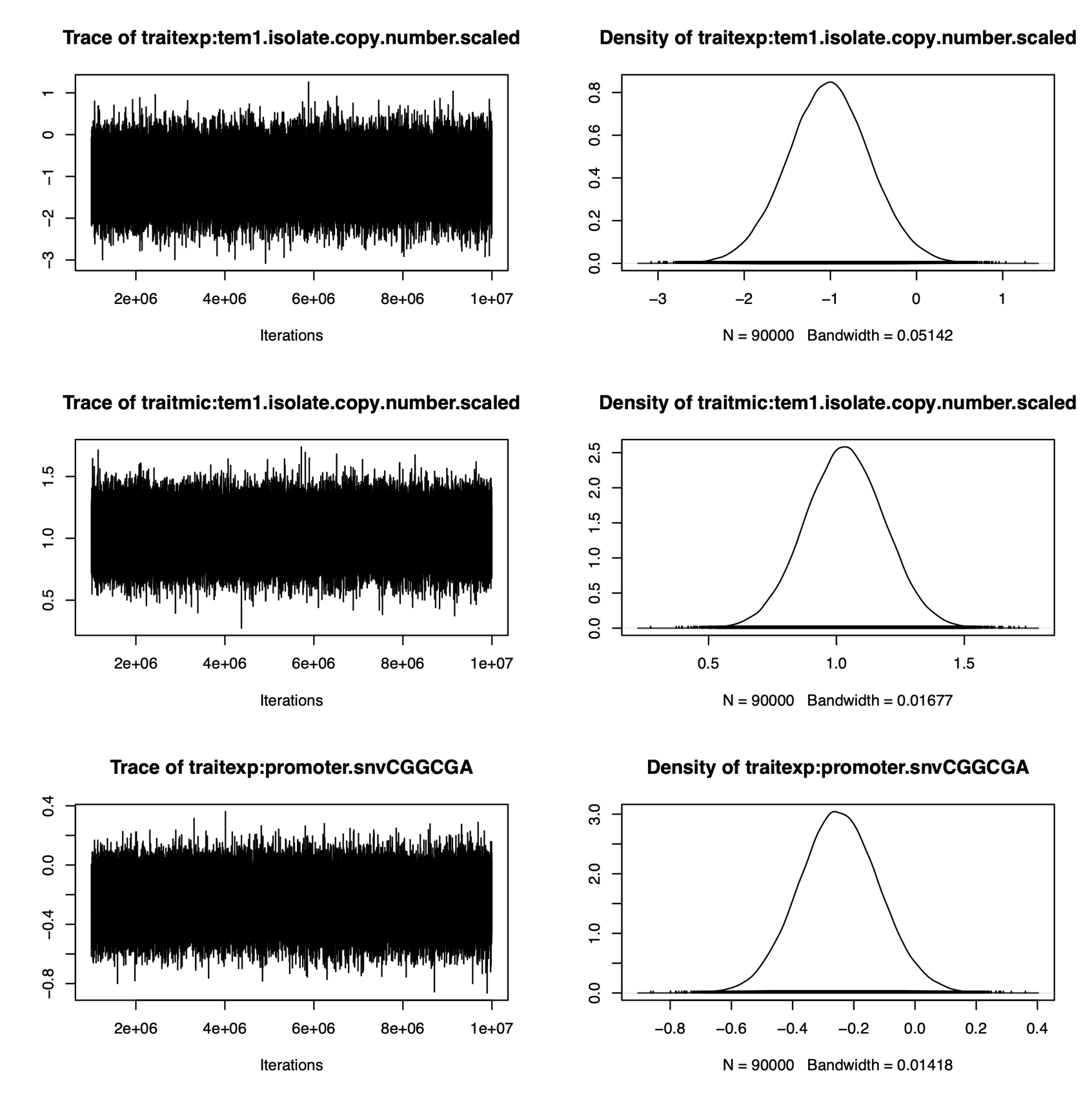

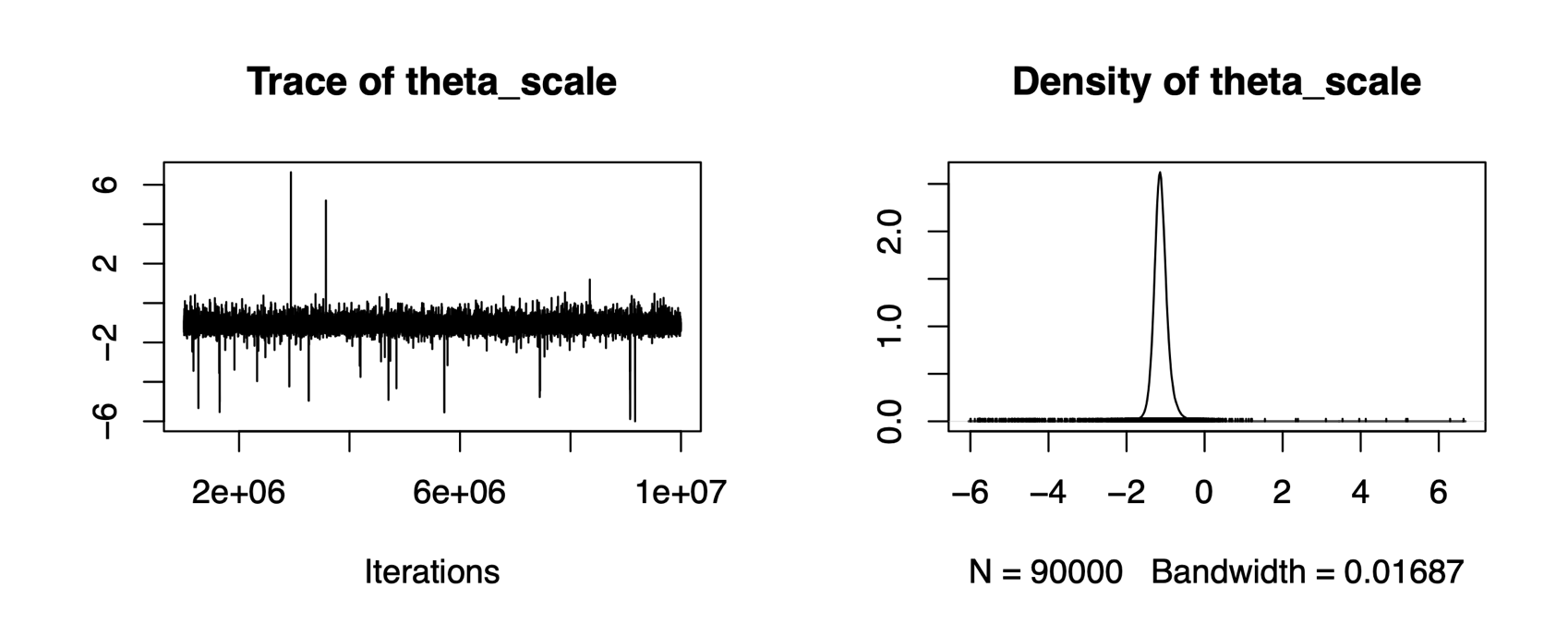

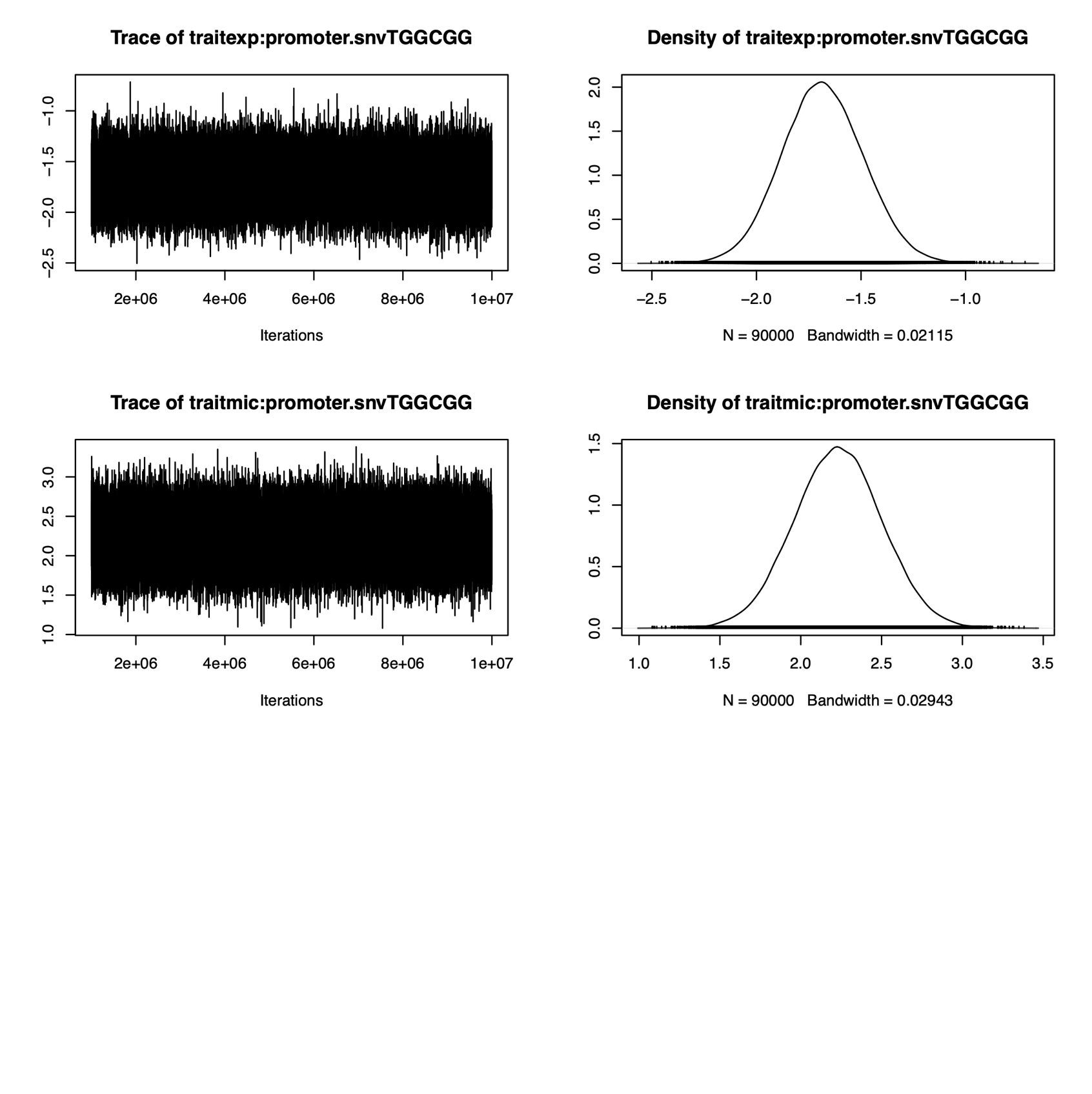

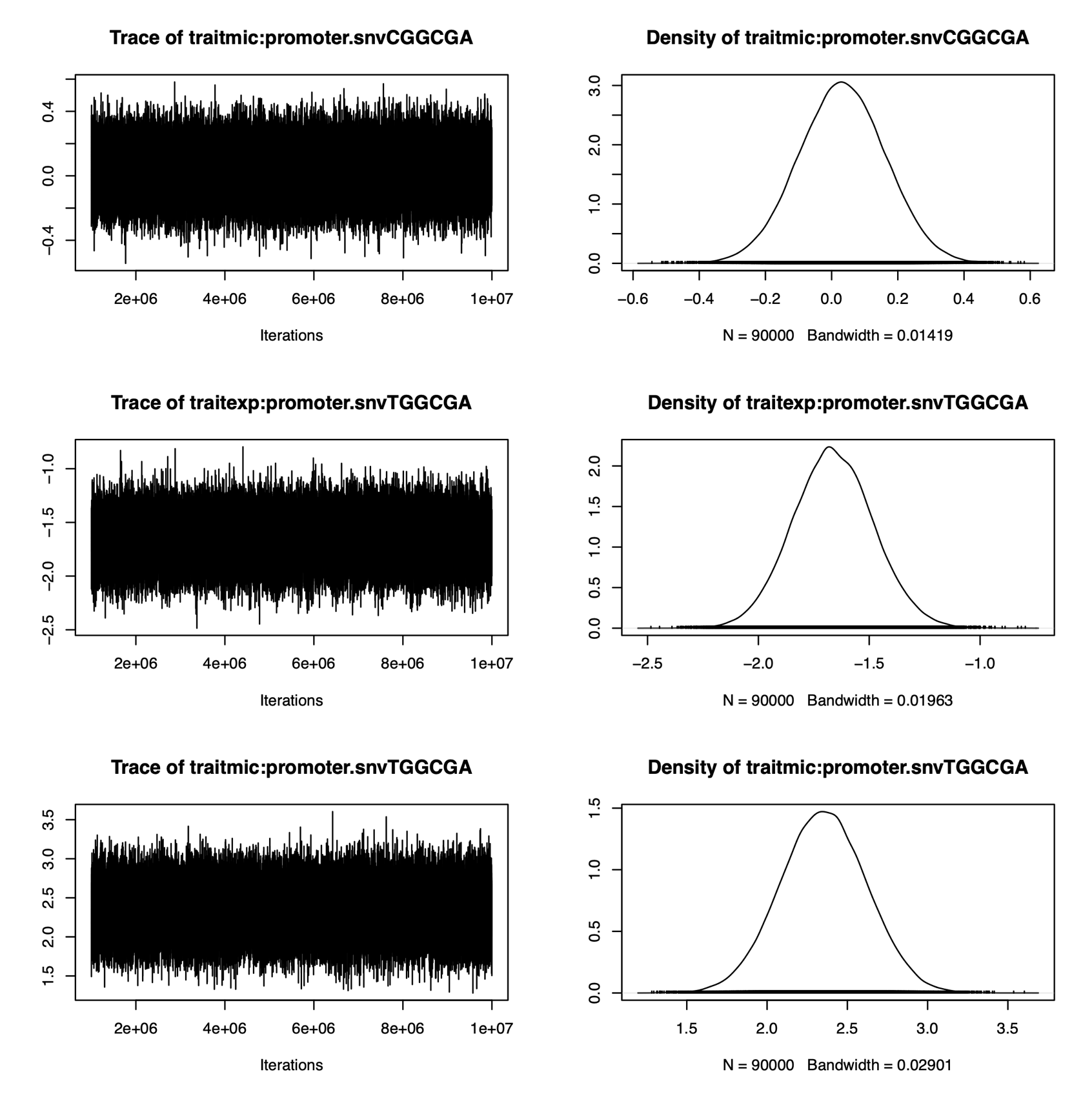

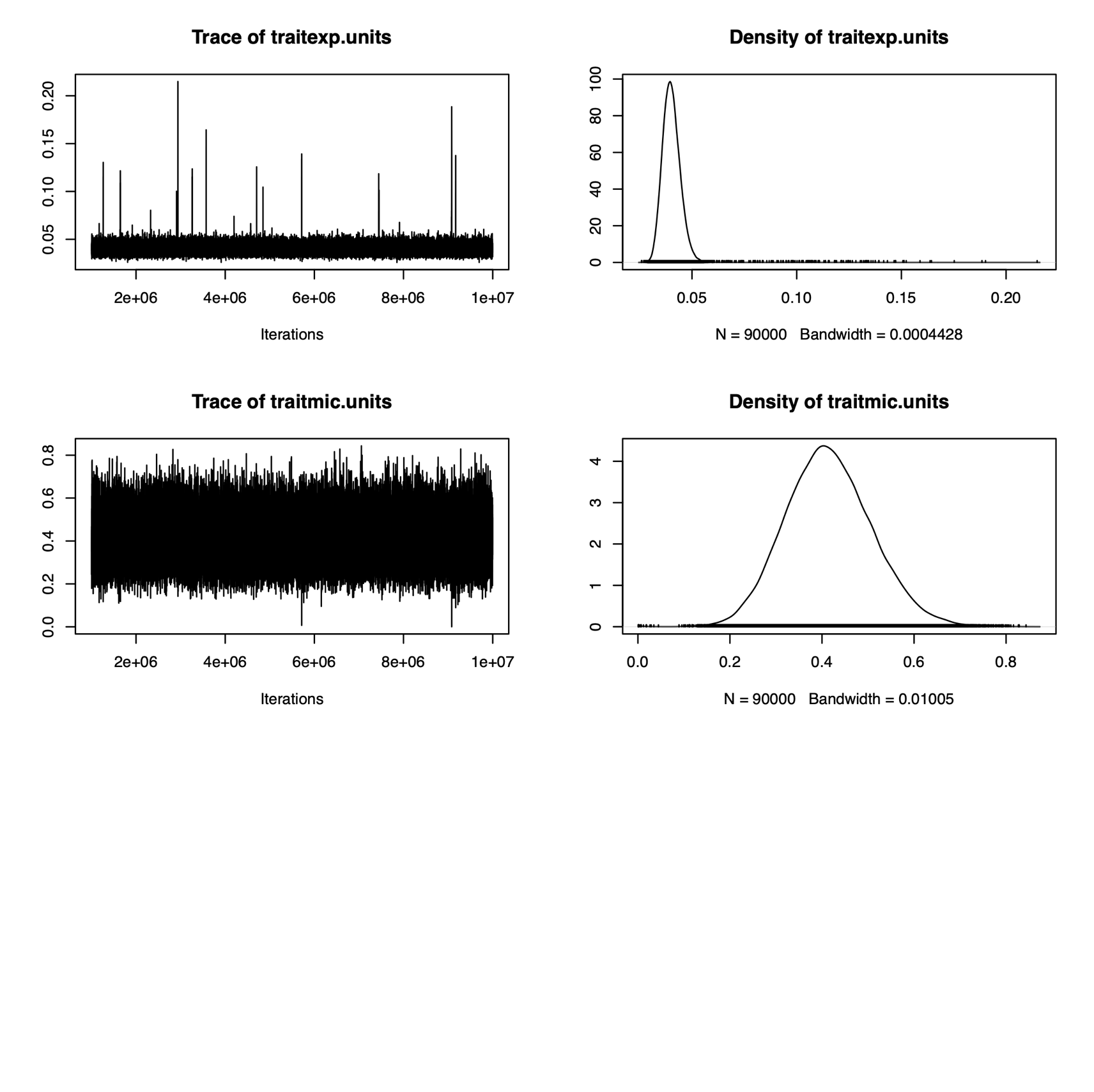

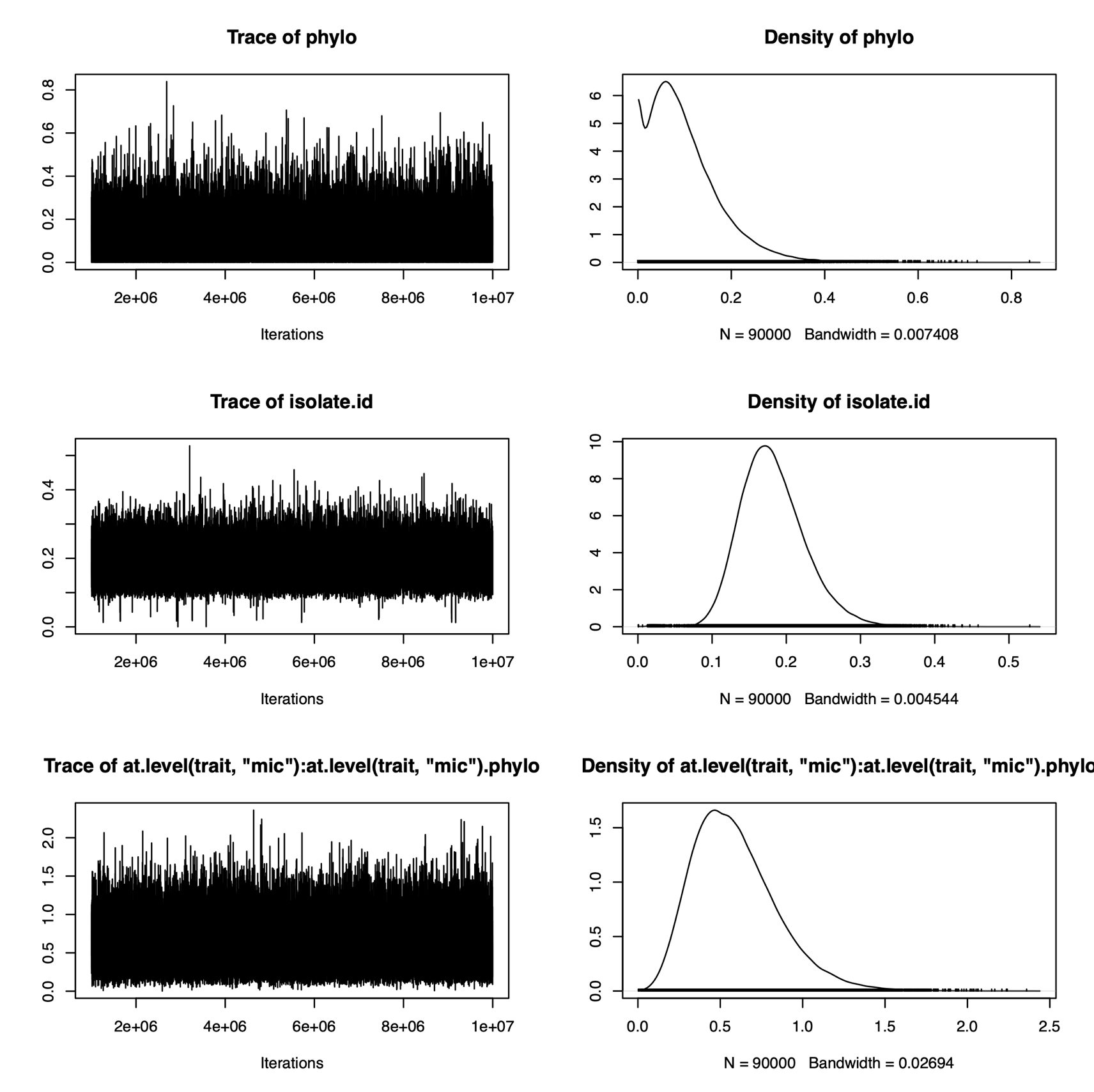

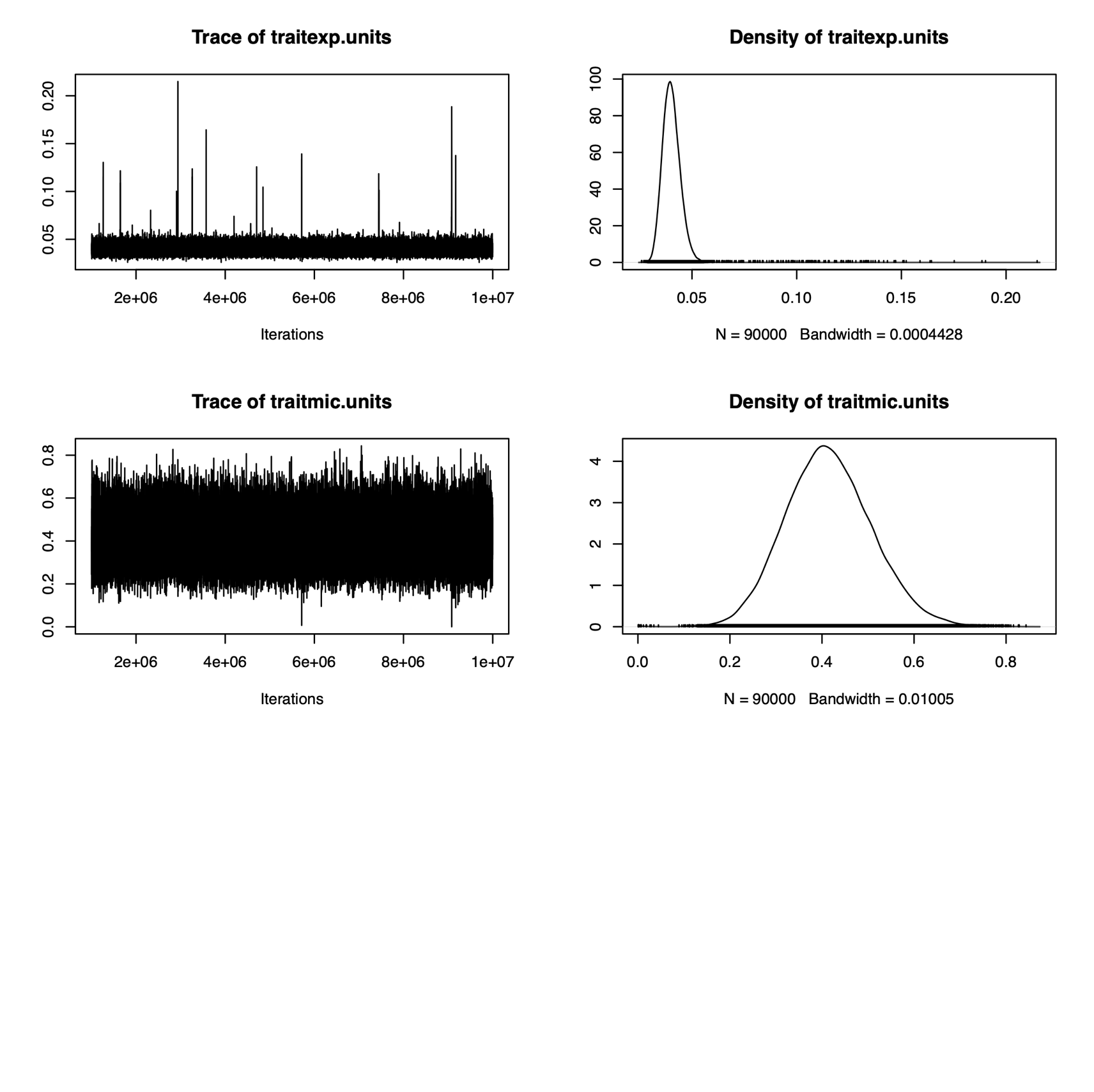

```
> mclist <- mcmc.list(chain.1$Sol, chain.2$Sol)
> gelman.diag(mclist)

Potential scale reduction factors:

                                         Point est. Upper C.I.
(Intercept)                                       1          1
traitexp:tem1.isolate.scaledFALSE                 1          1
traitmic:tem1.isolate.scaledFALSE                 1          1
traitexp:tem1.isolate.copy.number.scaled          1          1
traitmic:tem1.isolate.copy.number.scaled          1          1
traitexp:promoter.snvCGGCGA                       1          1
traitmic:promoter.snvCGGCGA                       1          1
traitexp:promoter.snvTGGCGA                       1          1
traitmic:promoter.snvTGGCGA                       1          1
traitexp:promoter.snvTGGCGG                       1          1
traitmic:promoter.snvTGGCGG                       1          1
phylo.542                                         1          1
phylo.416                                         1          1
phylo.56                                          1          1
phylo.213                                         1          1
phylo.54                                          1          1
phylo.435                                         1          1
phylo.90                                          1          1
phylo.393                                         1          1
phylo.72                                          1          1
phylo.489                                         1          1
phylo.189                                         1          1
phylo.434                                         1          1
phylo.194                                         1          1
phylo.371                                         1          1
phylo.127                                         1          1
phylo.99                                          1          1
phylo.129                                         1          1
phylo.8                                           1          1
phylo.414                                         1          1
phylo.454                                         1          1
phylo.120                                         1          1
phylo.145                                         1          1
phylo.426                                         1          1
phylo.394                                         1          1
phylo.183                                         1          1
phylo.30                                          1          1
phylo.151                                         1          1
phylo.148                                         1          1
phylo.236                                         1          1
phylo.220                                         1          1
phylo.117                                         1          1
phylo.272                                         1          1
phylo.400                                         1          1
phylo.328                                         1          1
phylo.415                                         1          1
phylo.398                                         1          1
phylo.252                                         1          1
phylo.246                                         1          1
phylo.424                                         1          1
phylo.476                                         1          1
phylo.468                                         1          1
phylo.363                                         1          1
phylo.21                                          1          1
phylo.128                                         1          1
phylo.257                                         1          1
phylo.242                                         1          1
phylo.339                                         1          1
phylo.149                                         1          1
phylo.445                                         1          1
phylo.61                                          1          1
phylo.299                                         1          1
phylo.420                                         1          1
phylo.103                                         1          1
phylo.464                                         1          1
phylo.449                                         1          1
phylo.97                                          1          1
phylo.208                                         1          1
phylo.329                                         1          1
phylo.316                                         1          1
phylo.80                                          1          1
phylo.438                                         1          1
phylo.216                                         1          1
phylo.324                                         1          1
phylo.224                                         1          1
phylo.447                                         1          1
phylo.266                                         1          1
phylo.136                                         1          1
phylo.156                                         1          1
phylo.144                                         1          1
phylo.404                                         1          1
phylo.202                                         1          1
phylo.439                                         1          1
phylo.366                                         1          1
phylo.285                                         1          1
phylo.133                                         1          1
phylo.284                                         1          1
phylo.512                                         1          1
phylo.155                                         1          1
phylo.37                                          1          1
phylo.317                                         1          1
phylo.143                                         1          1
phylo.475                                         1          1
phylo.342                                         1          1
phylo.471                                         1          1
phylo.264                                         1          1
phylo.517                                         1          1
phylo.507                                         1          1
phylo.132                                         1          1
phylo.39                                          1          1
phylo.286                                         1          1
phylo.524                                         1          1
phylo.533                                         1          1
phylo.333                                         1          1
phylo.410                                         1          1
phylo.446                                         1          1
phylo.344                                         1          1
phylo.406                                         1          1
phylo.22                                          1          1
phylo.430                                         1          1
phylo.288                                         1          1
phylo.511                                         1          1
phylo.429                                         1          1
phylo.277                                         1          1
phylo.322                                         1          1
phylo.262                                         1          1
phylo.387                                         1          1
phylo.58                                          1          1
phylo.388                                         1          1
phylo.158                                         1          1
phylo.12                                          1          1
phylo.276                                         1          1
phylo.412                                         1          1
phylo.411                                         1          1
phylo.365                                         1          1
phylo.82                                          1          1
phylo.92                                          1          1
phylo.370                                         1          1
phylo.137                                         1          1
phylo.267                                         1          1
phylo.421                                         1          1
phylo.237                                         1          1
phylo.200                                         1          1
phylo.536                                         1          1
phylo.260                                         1          1
phylo.537                                         1          1
phylo.477                                         1          1
phylo.518                                         1          1
phylo.55                                          1          1
phylo.358                                         1          1
phylo.197                                         1          1
phylo.493                                         1          1
phylo.41                                          1          1
phylo.40                                          1          1
phylo.408                                         1          1
phylo.292                                         1          1
phylo.248                                         1          1
phylo.1                                           1          1
phylo.214                                         1          1
phylo.480                                         1          1
phylo.423                                         1          1
phylo.247                                         1          1
phylo.268                                         1          1
phylo.353                                         1          1
phylo.211                                         1          1
phylo.265                                         1          1
phylo.139                                         1          1
phylo.198                                         1          1
phylo.396                                         1          1
phylo.263                                         1          1
phylo.95                                          1          1
phylo.162                                         1          1
phylo.515                                         1          1
phylo.282                                         1          1
phylo.427                                         1          1
phylo.240                                         1          1
phylo.166                                         1          1
phylo.182                                         1          1
phylo.532                                         1          1
phylo.273                                         1          1
phylo.119                                         1          1
phylo.355                                         1          1
phylo.241                                         1          1
phylo.203                                         1          1
phylo.121                                         1          1
phylo.305                                         1          1
phylo.534                                         1          1
phylo.167                                         1          1
phylo.336                                         1          1
phylo.188                                         1          1
phylo.274                                         1          1
phylo.281                                         1          1
phylo.89                                          1          1
phylo.531                                         1          1
phylo.164                                         1          1
phylo.382                                         1          1
phylo.192                                         1          1
phylo.301                                         1          1
phylo.506                                         1          1
phylo.334                                         1          1
phylo.469                                         1          1
phylo.2                                           1          1
phylo.470                                         1          1
phylo.351                                         1          1
phylo.31                                          1          1
phylo.159                                         1          1
phylo.187                                         1          1
phylo.488                                         1          1
phylo.486                                         1          1
phylo.225                                         1          1
phylo.32                                          1          1
phylo.541                                         1          1
phylo.380                                         1          1
phylo.462                                         1          1
phylo.451                                         1          1
phylo.403                                         1          1
phylo.115                                         1          1
phylo.250                                         1          1
phylo.93                                          1          1
phylo.230                                         1          1
phylo.457                                         1          1
phylo.313                                         1          1
phylo.331                                         1          1
phylo.176                                         1          1
phylo.75                                          1          1
phylo.296                                         1          1
phylo.205                                         1          1
phylo.256                                         1          1
phylo.487                                         1          1
phylo.347                                         1          1
phylo.442                                         1          1
phylo.495                                         1          1
phylo.116                                         1          1
phylo.233                                         1          1
phylo.539                                         1          1
phylo.83                                          1          1
phylo.444                                         1          1
phylo.508                                         1          1
phylo.33                                          1          1
phylo.142                                         1          1
phylo.311                                         1          1
phylo.386                                         1          1
phylo.135                                         1          1
phylo.19                                          1          1
phylo.59                                          1          1
phylo.287                                         1          1
phylo.375                                         1          1
phylo.490                                         1          1
phylo.448                                         1          1
phylo.10                                          1          1
phylo.399                                         1          1
phylo.466                                         1          1
phylo.310                                         1          1
phylo.543                                         1          1
phylo.504                                         1          1
phylo.174                                         1          1
phylo.499                                         1          1
phylo.126                                         1          1
phylo.413                                         1          1
phylo.125                                         1          1
phylo.201                                         1          1
phylo.513                                         1          1
phylo.69                                          1          1
phylo.340                                         1          1
phylo.179                                         1          1
phylo.25                                          1          1
phylo.436                                         1          1
phylo.546                                         1          1
phylo.279                                         1          1
phylo.485                                         1          1
phylo.38                                          1          1
phylo.422                                         1          1
phylo.229                                         1          1
phylo.35                                          1          1
phylo.131                                         1          1
phylo.71                                          1          1
phylo.6                                           1          1
phylo.401                                         1          1
phylo.222                                         1          1
phylo.243                                         1          1
phylo.42                                          1          1
phylo.63                                          1          1
phylo.18                                          1          1
phylo.123                                         1          1
phylo.140                                         1          1
phylo.502                                         1          1
phylo.503                                         1          1
phylo.341                                         1          1
phylo.238                                         1          1
phylo.478                                         1          1
phylo.443                                         1          1
phylo.44                                          1          1
phylo.544                                         1          1
phylo.204                                         1          1
phylo.510                                         1          1
phylo.314                                         1          1
phylo.124                                         1          1
phylo.64                                          1          1
phylo.219                                         1          1
phylo.178                                         1          1
phylo.345                                         1          1
phylo.479                                         1          1
phylo.307                                         1          1
phylo.275                                         1          1
phylo.540                                         1          1
phylo.23                                          1          1
phylo.73                                          1          1
phylo.530                                         1          1
phylo.244                                         1          1
phylo.191                                         1          1
phylo.474                                         1          1
phylo.460                                         1          1
phylo.528                                         1          1
phylo.295                                         1          1
phylo.459                                         1          1
phylo.463                                         1          1
phylo.79                                          1          1
phylo.481                                         1          1
phylo.190                                         1          1
phylo.461                                         1          1
phylo.535                                         1          1
phylo.223                                         1          1
phylo.29                                          1          1
phylo.65                                          1          1
phylo.283                                         1          1
phylo.425                                         1          1
phylo.107                                         1          1
phylo.294                                         1          1
phylo.122                                         1          1
phylo.105                                         1          1
phylo.138                                         1          1
phylo.440                                         1          1
isolate.id.OXEC-1                                 1          1
isolate.id.OXEC-10                                1          1
isolate.id.OXEC-103                               1          1
isolate.id.OXEC-105                               1          1
isolate.id.OXEC-107                               1          1
isolate.id.OXEC-115                               1          1
isolate.id.OXEC-116                               1          1
isolate.id.OXEC-117                               1          1
isolate.id.OXEC-119                               1          1
isolate.id.OXEC-12                                1          1
isolate.id.OXEC-120                               1          1
isolate.id.OXEC-121                               1          1
isolate.id.OXEC-122                               1          1
isolate.id.OXEC-123                               1          1
isolate.id.OXEC-124                               1          1
isolate.id.OXEC-125                               1          1
isolate.id.OXEC-126                               1          1
isolate.id.OXEC-127                               1          1
isolate.id.OXEC-128                               1          1
isolate.id.OXEC-129                               1          1
isolate.id.OXEC-131                               1          1
isolate.id.OXEC-132                               1          1
isolate.id.OXEC-133                               1          1
isolate.id.OXEC-135                               1          1
isolate.id.OXEC-136                               1          1
isolate.id.OXEC-137                               1          1
isolate.id.OXEC-138                               1          1
isolate.id.OXEC-139                               1          1
isolate.id.OXEC-140                               1          1
isolate.id.OXEC-142                               1          1
isolate.id.OXEC-143                               1          1
isolate.id.OXEC-144                               1          1
isolate.id.OXEC-145                               1          1
isolate.id.OXEC-148                               1          1
isolate.id.OXEC-149                               1          1
isolate.id.OXEC-151                               1          1
isolate.id.OXEC-155                               1          1
isolate.id.OXEC-156                               1          1
isolate.id.OXEC-158                               1          1
isolate.id.OXEC-159                               1          1
isolate.id.OXEC-162                               1          1
isolate.id.OXEC-164                               1          1
isolate.id.OXEC-166                               1          1
isolate.id.OXEC-167                               1          1
isolate.id.OXEC-174                               1          1
isolate.id.OXEC-176                               1          1
isolate.id.OXEC-178                               1          1
isolate.id.OXEC-179                               1          1
isolate.id.OXEC-18                                1          1
isolate.id.OXEC-182                               1          1
isolate.id.OXEC-183                               1          1
isolate.id.OXEC-187                               1          1
isolate.id.OXEC-188                               1          1
isolate.id.OXEC-189                               1          1
isolate.id.OXEC-19                                1          1
isolate.id.OXEC-190                               1          1
isolate.id.OXEC-191                               1          1
isolate.id.OXEC-192                               1          1
isolate.id.OXEC-194                               1          1
isolate.id.OXEC-197                               1          1
isolate.id.OXEC-198                               1          1
isolate.id.OXEC-2                                 1          1
isolate.id.OXEC-200                               1          1
isolate.id.OXEC-201                               1          1
isolate.id.OXEC-202                               1          1
isolate.id.OXEC-203                               1          1
isolate.id.OXEC-204                               1          1
isolate.id.OXEC-205                               1          1
isolate.id.OXEC-208                               1          1
isolate.id.OXEC-21                                1          1
isolate.id.OXEC-211                               1          1
isolate.id.OXEC-213                               1          1
isolate.id.OXEC-214                               1          1
isolate.id.OXEC-216                               1          1
isolate.id.OXEC-219                               1          1
isolate.id.OXEC-22                                1          1
isolate.id.OXEC-220                               1          1
isolate.id.OXEC-222                               1          1
isolate.id.OXEC-223                               1          1
isolate.id.OXEC-224                               1          1
isolate.id.OXEC-225                               1          1
isolate.id.OXEC-229                               1          1
isolate.id.OXEC-23                                1          1
isolate.id.OXEC-230                               1          1
isolate.id.OXEC-233                               1          1
isolate.id.OXEC-236                               1          1
isolate.id.OXEC-237                               1          1
isolate.id.OXEC-238                               1          1
isolate.id.OXEC-240                               1          1
isolate.id.OXEC-241                               1          1
isolate.id.OXEC-242                               1          1
isolate.id.OXEC-243                               1          1
isolate.id.OXEC-244                               1          1
isolate.id.OXEC-246                               1          1
isolate.id.OXEC-247                               1          1
isolate.id.OXEC-248                               1          1
isolate.id.OXEC-25                                1          1
isolate.id.OXEC-250                               1          1
isolate.id.OXEC-252                               1          1
isolate.id.OXEC-256                               1          1
isolate.id.OXEC-257                               1          1
isolate.id.OXEC-260                               1          1
isolate.id.OXEC-262                               1          1
isolate.id.OXEC-263                               1          1
isolate.id.OXEC-264                               1          1
isolate.id.OXEC-265                               1          1
isolate.id.OXEC-266                               1          1
isolate.id.OXEC-267                               1          1
isolate.id.OXEC-268                               1          1
isolate.id.OXEC-272                               1          1
isolate.id.OXEC-273                               1          1
isolate.id.OXEC-274                               1          1
isolate.id.OXEC-275                               1          1
isolate.id.OXEC-276                               1          1
isolate.id.OXEC-277                               1          1
isolate.id.OXEC-279                               1          1
isolate.id.OXEC-281                               1          1
isolate.id.OXEC-282                               1          1
isolate.id.OXEC-283                               1          1
isolate.id.OXEC-284                               1          1
isolate.id.OXEC-285                               1          1
isolate.id.OXEC-286                               1          1
isolate.id.OXEC-287                               1          1
isolate.id.OXEC-288                               1          1
isolate.id.OXEC-29                                1          1
isolate.id.OXEC-292                               1          1
isolate.id.OXEC-294                               1          1
isolate.id.OXEC-295                               1          1
isolate.id.OXEC-296                               1          1
isolate.id.OXEC-299                               1          1
isolate.id.OXEC-30                                1          1
isolate.id.OXEC-301                               1          1
isolate.id.OXEC-305                               1          1
isolate.id.OXEC-307                               1          1
isolate.id.OXEC-31                                1          1
isolate.id.OXEC-310                               1          1
isolate.id.OXEC-311                               1          1
isolate.id.OXEC-313                               1          1
isolate.id.OXEC-314                               1          1
isolate.id.OXEC-316                               1          1
isolate.id.OXEC-317                               1          1
isolate.id.OXEC-32                                1          1
isolate.id.OXEC-322                               1          1
isolate.id.OXEC-324                               1          1
isolate.id.OXEC-328                               1          1
isolate.id.OXEC-329                               1          1
isolate.id.OXEC-33                                1          1
isolate.id.OXEC-331                               1          1
isolate.id.OXEC-333                               1          1
isolate.id.OXEC-334                               1          1
isolate.id.OXEC-336                               1          1
isolate.id.OXEC-339                               1          1
isolate.id.OXEC-340                               1          1
isolate.id.OXEC-341                               1          1
isolate.id.OXEC-342                               1          1
isolate.id.OXEC-344                               1          1
isolate.id.OXEC-345                               1          1
isolate.id.OXEC-347                               1          1
isolate.id.OXEC-35                                1          1
isolate.id.OXEC-351                               1          1
isolate.id.OXEC-353                               1          1
isolate.id.OXEC-355                               1          1
isolate.id.OXEC-358                               1          1
isolate.id.OXEC-363                               1          1
isolate.id.OXEC-365                               1          1
isolate.id.OXEC-366                               1          1
isolate.id.OXEC-37                                1          1
isolate.id.OXEC-370                               1          1
isolate.id.OXEC-371                               1          1
isolate.id.OXEC-375                               1          1
isolate.id.OXEC-38                                1          1
isolate.id.OXEC-380                               1          1
isolate.id.OXEC-382                               1          1
isolate.id.OXEC-386                               1          1
isolate.id.OXEC-387                               1          1
isolate.id.OXEC-388                               1          1
isolate.id.OXEC-39                                1          1
isolate.id.OXEC-393                               1          1
isolate.id.OXEC-394                               1          1
isolate.id.OXEC-396                               1          1
isolate.id.OXEC-398                               1          1
isolate.id.OXEC-399                               1          1
isolate.id.OXEC-40                                1          1
isolate.id.OXEC-400                               1          1
isolate.id.OXEC-401                               1          1
isolate.id.OXEC-403                               1          1
isolate.id.OXEC-404                               1          1
isolate.id.OXEC-406                               1          1
isolate.id.OXEC-408                               1          1
isolate.id.OXEC-41                                1          1
isolate.id.OXEC-410                               1          1
isolate.id.OXEC-411                               1          1
isolate.id.OXEC-412                               1          1
isolate.id.OXEC-413                               1          1
isolate.id.OXEC-414                               1          1
isolate.id.OXEC-415                               1          1
isolate.id.OXEC-416                               1          1
isolate.id.OXEC-42                                1          1
isolate.id.OXEC-420                               1          1
isolate.id.OXEC-421                               1          1
isolate.id.OXEC-422                               1          1
isolate.id.OXEC-423                               1          1
isolate.id.OXEC-424                               1          1
isolate.id.OXEC-425                               1          1
isolate.id.OXEC-426                               1          1
isolate.id.OXEC-427                               1          1
isolate.id.OXEC-429                               1          1
isolate.id.OXEC-430                               1          1
isolate.id.OXEC-434                               1          1
isolate.id.OXEC-435                               1          1
isolate.id.OXEC-436                               1          1
isolate.id.OXEC-438                               1          1
isolate.id.OXEC-439                               1          1
isolate.id.OXEC-44                                1          1
isolate.id.OXEC-440                               1          1
isolate.id.OXEC-442                               1          1
isolate.id.OXEC-443                               1          1
isolate.id.OXEC-444                               1          1
isolate.id.OXEC-445                               1          1
isolate.id.OXEC-446                               1          1
isolate.id.OXEC-447                               1          1
isolate.id.OXEC-448                               1          1
isolate.id.OXEC-449                               1          1
isolate.id.OXEC-451                               1          1
isolate.id.OXEC-454                               1          1
isolate.id.OXEC-457                               1          1
isolate.id.OXEC-459                               1          1
isolate.id.OXEC-460                               1          1
isolate.id.OXEC-461                               1          1
isolate.id.OXEC-462                               1          1
isolate.id.OXEC-463                               1          1
isolate.id.OXEC-464                               1          1
isolate.id.OXEC-466                               1          1
isolate.id.OXEC-468                               1          1
isolate.id.OXEC-469                               1          1
isolate.id.OXEC-470                               1          1
isolate.id.OXEC-471                               1          1
isolate.id.OXEC-474                               1          1
isolate.id.OXEC-475                               1          1
isolate.id.OXEC-476                               1          1
isolate.id.OXEC-477                               1          1
isolate.id.OXEC-478                               1          1
isolate.id.OXEC-479                               1          1
isolate.id.OXEC-480                               1          1
isolate.id.OXEC-481                               1          1
isolate.id.OXEC-485                               1          1
isolate.id.OXEC-486                               1          1
isolate.id.OXEC-487                               1          1
isolate.id.OXEC-488                               1          1
isolate.id.OXEC-489                               1          1
isolate.id.OXEC-490                               1          1
isolate.id.OXEC-493                               1          1
isolate.id.OXEC-495                               1          1
isolate.id.OXEC-499                               1          1
isolate.id.OXEC-502                               1          1
isolate.id.OXEC-503                               1          1
isolate.id.OXEC-504                               1          1
isolate.id.OXEC-506                               1          1
isolate.id.OXEC-507                               1          1
isolate.id.OXEC-508                               1          1
isolate.id.OXEC-510                               1          1
isolate.id.OXEC-511                               1          1
isolate.id.OXEC-512                               1          1
isolate.id.OXEC-513                               1          1
isolate.id.OXEC-515                               1          1
isolate.id.OXEC-517                               1          1
isolate.id.OXEC-518                               1          1
isolate.id.OXEC-524                               1          1
isolate.id.OXEC-528                               1          1
isolate.id.OXEC-530                               1          1
isolate.id.OXEC-531                               1          1
isolate.id.OXEC-532                               1          1
isolate.id.OXEC-533                               1          1
isolate.id.OXEC-534                               1          1
isolate.id.OXEC-535                               1          1
isolate.id.OXEC-536                               1          1
isolate.id.OXEC-537                               1          1
isolate.id.OXEC-539                               1          1
isolate.id.OXEC-54                                1          1
isolate.id.OXEC-540                               1          1
isolate.id.OXEC-541                               1          1
isolate.id.OXEC-542                               1          1
isolate.id.OXEC-543                               1          1
isolate.id.OXEC-544                               1          1
isolate.id.OXEC-546                               1          1
isolate.id.OXEC-55                                1          1
isolate.id.OXEC-56                                1          1
isolate.id.OXEC-58                                1          1
isolate.id.OXEC-59                                1          1
isolate.id.OXEC-6                                 1          1
isolate.id.OXEC-61                                1          1
isolate.id.OXEC-63                                1          1
isolate.id.OXEC-64                                1          1
isolate.id.OXEC-65                                1          1
isolate.id.OXEC-69                                1          1
isolate.id.OXEC-71                                1          1
isolate.id.OXEC-72                                1          1
isolate.id.OXEC-73                                1          1
isolate.id.OXEC-75                                1          1
isolate.id.OXEC-79                                1          1
isolate.id.OXEC-8                                 1          1
isolate.id.OXEC-80                                1          1
isolate.id.OXEC-82                                1          1
isolate.id.OXEC-83                                1          1
isolate.id.OXEC-89                                1          1
isolate.id.OXEC-90                                1          1
isolate.id.OXEC-92                                1          1
isolate.id.OXEC-93                                1          1
isolate.id.OXEC-95                                1          1
isolate.id.OXEC-97                                1          1
isolate.id.OXEC-99                                1          1
at.level(trait, "mic").phylo.542                  1          1
at.level(trait, "mic").phylo.416                  1          1
at.level(trait, "mic").phylo.56                   1          1
at.level(trait, "mic").phylo.213                  1          1
at.level(trait, "mic").phylo.54                   1          1
at.level(trait, "mic").phylo.435                  1          1
at.level(trait, "mic").phylo.90                   1          1
at.level(trait, "mic").phylo.393                  1          1
at.level(trait, "mic").phylo.72                   1          1
at.level(trait, "mic").phylo.489                  1          1
at.level(trait, "mic").phylo.189                  1          1
at.level(trait, "mic").phylo.434                  1          1
at.level(trait, "mic").phylo.194                  1          1
at.level(trait, "mic").phylo.371                  1          1
at.level(trait, "mic").phylo.127                  1          1
at.level(trait, "mic").phylo.99                   1          1
at.level(trait, "mic").phylo.129                  1          1
at.level(trait, "mic").phylo.8                    1          1
at.level(trait, "mic").phylo.414                  1          1
at.level(trait, "mic").phylo.454                  1          1
at.level(trait, "mic").phylo.120                  1          1
at.level(trait, "mic").phylo.145                  1          1
at.level(trait, "mic").phylo.426                  1          1
at.level(trait, "mic").phylo.394                  1          1
at.level(trait, "mic").phylo.183                  1          1
at.level(trait, "mic").phylo.30                   1          1
at.level(trait, "mic").phylo.151                  1          1
at.level(trait, "mic").phylo.148                  1          1
at.level(trait, "mic").phylo.236                  1          1
at.level(trait, "mic").phylo.220                  1          1
at.level(trait, "mic").phylo.117                  1          1
at.level(trait, "mic").phylo.272                  1          1
at.level(trait, "mic").phylo.400                  1          1
at.level(trait, "mic").phylo.328                  1          1
at.level(trait, "mic").phylo.415                  1          1
at.level(trait, "mic").phylo.398                  1          1
at.level(trait, "mic").phylo.252                  1          1
at.level(trait, "mic").phylo.246                  1          1
at.level(trait, "mic").phylo.424                  1          1
at.level(trait, "mic").phylo.476                  1          1
at.level(trait, "mic").phylo.468                  1          1
at.level(trait, "mic").phylo.363                  1          1
at.level(trait, "mic").phylo.21                   1          1
at.level(trait, "mic").phylo.128                  1          1
at.level(trait, "mic").phylo.257                  1          1
at.level(trait, "mic").phylo.242                  1          1
at.level(trait, "mic").phylo.339                  1          1
at.level(trait, "mic").phylo.149                  1          1
at.level(trait, "mic").phylo.445                  1          1
at.level(trait, "mic").phylo.61                   1          1
at.level(trait, "mic").phylo.299                  1          1
at.level(trait, "mic").phylo.420                  1          1
at.level(trait, "mic").phylo.103                  1          1
at.level(trait, "mic").phylo.464                  1          1
at.level(trait, "mic").phylo.449                  1          1
at.level(trait, "mic").phylo.97                   1          1
at.level(trait, "mic").phylo.208                  1          1
at.level(trait, "mic").phylo.329                  1          1
at.level(trait, "mic").phylo.316                  1          1
at.level(trait, "mic").phylo.80                   1          1
at.level(trait, "mic").phylo.438                  1          1
at.level(trait, "mic").phylo.216                  1          1
at.level(trait, "mic").phylo.324                  1          1
at.level(trait, "mic").phylo.224                  1          1
at.level(trait, "mic").phylo.447                  1          1
at.level(trait, "mic").phylo.266                  1          1
at.level(trait, "mic").phylo.136                  1          1
at.level(trait, "mic").phylo.156                  1          1
at.level(trait, "mic").phylo.144                  1          1
at.level(trait, "mic").phylo.404                  1          1
at.level(trait, "mic").phylo.202                  1          1
at.level(trait, "mic").phylo.439                  1          1
at.level(trait, "mic").phylo.366                  1          1
at.level(trait, "mic").phylo.285                  1          1
at.level(trait, "mic").phylo.133                  1          1
at.level(trait, "mic").phylo.284                  1          1
at.level(trait, "mic").phylo.512                  1          1
at.level(trait, "mic").phylo.155                  1          1
at.level(trait, "mic").phylo.37                   1          1
at.level(trait, "mic").phylo.317                  1          1
at.level(trait, "mic").phylo.143                  1          1
at.level(trait, "mic").phylo.475                  1          1
at.level(trait, "mic").phylo.342                  1          1
at.level(trait, "mic").phylo.471                  1          1
at.level(trait, "mic").phylo.264                  1          1
at.level(trait, "mic").phylo.517                  1          1
at.level(trait, "mic").phylo.507                  1          1
at.level(trait, "mic").phylo.132                  1          1
at.level(trait, "mic").phylo.39                   1          1
at.level(trait, "mic").phylo.286                  1          1
at.level(trait, "mic").phylo.524                  1          1
at.level(trait, "mic").phylo.533                  1          1
at.level(trait, "mic").phylo.333                  1          1
at.level(trait, "mic").phylo.410                  1          1
at.level(trait, "mic").phylo.446                  1          1
at.level(trait, "mic").phylo.344                  1          1
at.level(trait, "mic").phylo.406                  1          1
at.level(trait, "mic").phylo.22                   1          1
at.level(trait, "mic").phylo.430                  1          1
at.level(trait, "mic").phylo.288                  1          1
at.level(trait, "mic").phylo.511                  1          1
at.level(trait, "mic").phylo.429                  1          1
at.level(trait, "mic").phylo.277                  1          1
at.level(trait, "mic").phylo.322                  1          1
at.level(trait, "mic").phylo.262                  1          1
at.level(trait, "mic").phylo.387                  1          1
at.level(trait, "mic").phylo.58                   1          1
at.level(trait, "mic").phylo.388                  1          1
at.level(trait, "mic").phylo.158                  1          1
at.level(trait, "mic").phylo.12                   1          1
at.level(trait, "mic").phylo.276                  1          1
at.level(trait, "mic").phylo.412                  1          1
at.level(trait, "mic").phylo.411                  1          1
at.level(trait, "mic").phylo.365                  1          1
at.level(trait, "mic").phylo.82                   1          1
at.level(trait, "mic").phylo.92                   1          1
at.level(trait, "mic").phylo.370                  1          1
at.level(trait, "mic").phylo.137                  1          1
at.level(trait, "mic").phylo.267                  1          1
at.level(trait, "mic").phylo.421                  1          1
at.level(trait, "mic").phylo.237                  1          1
at.level(trait, "mic").phylo.200                  1          1
at.level(trait, "mic").phylo.536                  1          1
at.level(trait, "mic").phylo.260                  1          1
at.level(trait, "mic").phylo.537                  1          1
at.level(trait, "mic").phylo.477                  1          1
at.level(trait, "mic").phylo.518                  1          1
at.level(trait, "mic").phylo.55                   1          1
at.level(trait, "mic").phylo.358                  1          1
at.level(trait, "mic").phylo.197                  1          1
at.level(trait, "mic").phylo.493                  1          1
at.level(trait, "mic").phylo.41                   1          1
at.level(trait, "mic").phylo.40                   1          1
at.level(trait, "mic").phylo.408                  1          1
at.level(trait, "mic").phylo.292                  1          1
at.level(trait, "mic").phylo.248                  1          1
at.level(trait, "mic").phylo.1                    1          1
at.level(trait, "mic").phylo.214                  1          1
at.level(trait, "mic").phylo.480                  1          1
at.level(trait, "mic").phylo.423                  1          1
at.level(trait, "mic").phylo.247                  1          1
at.level(trait, "mic").phylo.268                  1          1
at.level(trait, "mic").phylo.353                  1          1
at.level(trait, "mic").phylo.211                  1          1
at.level(trait, "mic").phylo.265                  1          1
at.level(trait, "mic").phylo.139                  1          1
at.level(trait, "mic").phylo.198                  1          1
at.level(trait, "mic").phylo.396                  1          1
at.level(trait, "mic").phylo.263                  1          1
at.level(trait, "mic").phylo.95                   1          1
at.level(trait, "mic").phylo.162                  1          1
at.level(trait, "mic").phylo.515                  1          1
at.level(trait, "mic").phylo.282                  1          1
at.level(trait, "mic").phylo.427                  1          1
at.level(trait, "mic").phylo.240                  1          1
at.level(trait, "mic").phylo.166                  1          1
at.level(trait, "mic").phylo.182                  1          1
at.level(trait, "mic").phylo.532                  1          1
at.level(trait, "mic").phylo.273                  1          1
at.level(trait, "mic").phylo.119                  1          1
at.level(trait, "mic").phylo.355                  1          1
at.level(trait, "mic").phylo.241                  1          1
at.level(trait, "mic").phylo.203                  1          1
at.level(trait, "mic").phylo.121                  1          1
at.level(trait, "mic").phylo.305                  1          1
at.level(trait, "mic").phylo.534                  1          1
at.level(trait, "mic").phylo.167                  1          1
at.level(trait, "mic").phylo.336                  1          1
at.level(trait, "mic").phylo.188                  1          1
at.level(trait, "mic").phylo.274                  1          1
at.level(trait, "mic").phylo.281                  1          1
at.level(trait, "mic").phylo.89                   1          1
at.level(trait, "mic").phylo.531                  1          1
at.level(trait, "mic").phylo.164                  1          1
at.level(trait, "mic").phylo.382                  1          1
at.level(trait, "mic").phylo.192                  1          1
at.level(trait, "mic").phylo.301                  1          1
at.level(trait, "mic").phylo.506                  1          1
at.level(trait, "mic").phylo.334                  1          1
at.level(trait, "mic").phylo.469                  1          1
at.level(trait, "mic").phylo.2                    1          1
at.level(trait, "mic").phylo.470                  1          1
at.level(trait, "mic").phylo.351                  1          1
at.level(trait, "mic").phylo.31                   1          1
at.level(trait, "mic").phylo.159                  1          1
at.level(trait, "mic").phylo.187                  1          1
at.level(trait, "mic").phylo.488                  1          1
at.level(trait, "mic").phylo.486                  1          1
at.level(trait, "mic").phylo.225                  1          1
at.level(trait, "mic").phylo.32                   1          1
at.level(trait, "mic").phylo.541                  1          1
at.level(trait, "mic").phylo.380                  1          1
at.level(trait, "mic").phylo.462                  1          1
at.level(trait, "mic").phylo.451                  1          1
at.level(trait, "mic").phylo.403                  1          1
at.level(trait, "mic").phylo.115                  1          1
at.level(trait, "mic").phylo.250                  1          1
at.level(trait, "mic").phylo.93                   1          1
at.level(trait, "mic").phylo.230                  1          1
at.level(trait, "mic").phylo.457                  1          1
at.level(trait, "mic").phylo.313                  1          1
at.level(trait, "mic").phylo.331                  1          1
at.level(trait, "mic").phylo.176                  1          1
at.level(trait, "mic").phylo.75                   1          1
at.level(trait, "mic").phylo.296                  1          1
at.level(trait, "mic").phylo.205                  1          1
at.level(trait, "mic").phylo.256                  1          1
at.level(trait, "mic").phylo.487                  1          1
at.level(trait, "mic").phylo.347                  1          1
at.level(trait, "mic").phylo.442                  1          1
at.level(trait, "mic").phylo.495                  1          1
at.level(trait, "mic").phylo.116                  1          1
at.level(trait, "mic").phylo.233                  1          1
at.level(trait, "mic").phylo.539                  1          1
at.level(trait, "mic").phylo.83                   1          1
at.level(trait, "mic").phylo.444                  1          1
at.level(trait, "mic").phylo.508                  1          1
at.level(trait, "mic").phylo.33                   1          1
at.level(trait, "mic").phylo.142                  1          1
at.level(trait, "mic").phylo.311                  1          1
at.level(trait, "mic").phylo.386                  1          1
at.level(trait, "mic").phylo.135                  1          1
at.level(trait, "mic").phylo.19                   1          1
at.level(trait, "mic").phylo.59                   1          1
at.level(trait, "mic").phylo.287                  1          1
at.level(trait, "mic").phylo.375                  1          1
at.level(trait, "mic").phylo.490                  1          1
at.level(trait, "mic").phylo.448                  1          1
at.level(trait, "mic").phylo.10                   1          1
at.level(trait, "mic").phylo.399                  1          1
at.level(trait, "mic").phylo.466                  1          1
at.level(trait, "mic").phylo.310                  1          1
at.level(trait, "mic").phylo.543                  1          1
at.level(trait, "mic").phylo.504                  1          1
at.level(trait, "mic").phylo.174                  1          1
at.level(trait, "mic").phylo.499                  1          1
at.level(trait, "mic").phylo.126                  1          1
at.level(trait, "mic").phylo.413                  1          1
at.level(trait, "mic").phylo.125                  1          1
at.level(trait, "mic").phylo.201                  1          1
at.level(trait, "mic").phylo.513                  1          1
at.level(trait, "mic").phylo.69                   1          1
at.level(trait, "mic").phylo.340                  1          1
at.level(trait, "mic").phylo.179                  1          1
at.level(trait, "mic").phylo.25                   1          1
at.level(trait, "mic").phylo.436                  1          1
at.level(trait, "mic").phylo.546                  1          1
at.level(trait, "mic").phylo.279                  1          1
at.level(trait, "mic").phylo.485                  1          1
at.level(trait, "mic").phylo.38                   1          1
at.level(trait, "mic").phylo.422                  1          1
at.level(trait, "mic").phylo.229                  1          1
at.level(trait, "mic").phylo.35                   1          1
at.level(trait, "mic").phylo.131                  1          1
at.level(trait, "mic").phylo.71                   1          1
at.level(trait, "mic").phylo.6                    1          1
at.level(trait, "mic").phylo.401                  1          1
at.level(trait, "mic").phylo.222                  1          1
at.level(trait, "mic").phylo.243                  1          1
at.level(trait, "mic").phylo.42                   1          1
at.level(trait, "mic").phylo.63                   1          1
at.level(trait, "mic").phylo.18                   1          1
at.level(trait, "mic").phylo.123                  1          1
at.level(trait, "mic").phylo.140                  1          1
at.level(trait, "mic").phylo.502                  1          1
at.level(trait, "mic").phylo.503                  1          1
at.level(trait, "mic").phylo.341                  1          1
at.level(trait, "mic").phylo.238                  1          1
at.level(trait, "mic").phylo.478                  1          1
at.level(trait, "mic").phylo.443                  1          1
at.level(trait, "mic").phylo.44                   1          1
at.level(trait, "mic").phylo.544                  1          1
at.level(trait, "mic").phylo.204                  1          1
at.level(trait, "mic").phylo.510                  1          1
at.level(trait, "mic").phylo.314                  1          1
at.level(trait, "mic").phylo.124                  1          1
at.level(trait, "mic").phylo.64                   1          1
at.level(trait, "mic").phylo.219                  1          1
at.level(trait, "mic").phylo.178                  1          1
at.level(trait, "mic").phylo.345                  1          1
at.level(trait, "mic").phylo.479                  1          1
at.level(trait, "mic").phylo.307                  1          1
at.level(trait, "mic").phylo.275                  1          1
at.level(trait, "mic").phylo.540                  1          1
at.level(trait, "mic").phylo.23                   1          1
at.level(trait, "mic").phylo.73                   1          1
at.level(trait, "mic").phylo.530                  1          1
at.level(trait, "mic").phylo.244                  1          1
at.level(trait, "mic").phylo.191                  1          1
at.level(trait, "mic").phylo.474                  1          1
at.level(trait, "mic").phylo.460                  1          1
at.level(trait, "mic").phylo.528                  1          1
at.level(trait, "mic").phylo.295                  1          1
at.level(trait, "mic").phylo.459                  1          1
at.level(trait, "mic").phylo.463                  1          1
at.level(trait, "mic").phylo.79                   1          1
at.level(trait, "mic").phylo.481                  1          1
at.level(trait, "mic").phylo.190                  1          1
at.level(trait, "mic").phylo.461                  1          1
at.level(trait, "mic").phylo.535                  1          1
at.level(trait, "mic").phylo.223                  1          1
at.level(trait, "mic").phylo.29                   1          1
at.level(trait, "mic").phylo.65                   1          1
at.level(trait, "mic").phylo.283                  1          1
at.level(trait, "mic").phylo.425                  1          1
at.level(trait, "mic").phylo.107                  1          1
at.level(trait, "mic").phylo.294                  1          1
at.level(trait, "mic").phylo.122                  1          1
at.level(trait, "mic").phylo.105                  1          1
at.level(trait, "mic").phylo.138                  1          1
at.level(trait, "mic").phylo.440                  1          1

Multivariate psrf

1.01
```
